# Diverse cell junctions with unique molecular composition in tissues of a sponge (Porifera)

**DOI:** 10.1101/685875

**Authors:** Jennyfer M. Mitchell, Scott A. Nichols

**Affiliations:** Department of Biological Sciences, 2101 E. Wesley Ave. SGM 203, University of Denver, Denver, CO 80208; University of Colorado, Anschutz Medical Campus, 12801 E. 17th Ave. RC1S, 11501G, Aurora, CO 80045

## Abstract

The integrity and organization of animal tissues depends upon specialized protein complexes that mediate adhesion between cells with each other (cadherin-based adherens junctions), and with the extracellular matrix (integrin-based focal adhesions). Reconstructing how and when these cell junctions evolved is central to understanding early tissue evolution in animals. We examined focal adhesion protein homologs in tissues of the freshwater sponge, *Ephydatia muelleri* (phylum Porifera). We found that sponge homologs of focal adhesion proteins co-precipitate as a complex and localize to cell junctions in sponge tissues. These data support that the adhesion roles of these proteins evolved early, prior to the divergence of sponges and other animals. However, in contrast to the spatially partitioned distribution of cell junctions in epithelia of other animals, focal adhesion proteins were found to be co-distributed with the adherens junction protein Emβ-catenin in sponge tissues; both at certain cell-cell and cell-extracellular matrix (ECM) adhesions. Sponge adhesion structures were found to be unique in other ways, too. The basopinacoderm (substrate-attachment epithelium) lacks typical polarity in that cell-ECM adhesions form on both basal and apical surfaces, and compositionally unique cell junctions form at the interface between cells with spicules (siliceous skeletal elements) and between cells and environmental bacteria. These results clarify the diversity, distribution and molecular composition of cell junctions in tissues of *E. muelleri*, but raise new questions about their function and homology with cell junctions in other animals.

## Introduction

Beyond simply gluing cells together, cell adhesion molecules are dynamically regulated during development and cell migration, spatially regulated in polarized tissues, and involved in cell signaling and mechanotransduction (Collins and Nelson, 2015; Halbleib and Nelson, 2006; Hynes, 2007, 1992; Kuo, 2013; Pinheiro and Bellaϊche, 2018; Zaman, 2008). Consequently, myriad adhesion mechanisms have evolved to function in different contexts in animals (e.g., (Ebnet, 2017)). Of these, two predominate: 1) the adherens junction, which is involved in cell-cell adhesion and is composed of cadherin receptors, p120-, α- and β-catenin, and 2) focal adhesions, which are involved in cell-extracellular matrix (ECM) adhesion and composed of protein such as integrins, vinculin, paxillin, talin and focal adhesion kinase (FAK).

The molecular components of both the adherens junction and focal adhesions are widely conserved in animals, and some of their components have origins outside of animals (Dickinson et al., 2011; Grau-Bové et al., 2017; Grimson et al., 2000; Murray and Zaidel-Bar, 2014; Sebé-Pedrós et al., 2010). However, experimental studies of cell junction composition and function are largely restricted to bilaterian animals, such as the roundworm *Caenorhabditis elegans*, the fruit fly *Drosophila melanogaster*, and vertebrates. Recent studies demonstrate conserved roles for adherens junction proteins in cnidarians, as well (Clarke et al., 2019, 2016; Pukhlyakova et al., n.d.).

Organisms of critical importance for reconstructing early steps in the evolution of animal cell adhesion mechanisms are the sponges (Porifera). They are one of the most phylogenetically divergent groups of animals (King and Rokas, 2017; Simion et al., 2017), their anatomy is fundamentally different from other animals (Leys and Hill, 2012), and there are long-standing questions about the structure and homology of their tissues compared to epithelia in other animals (Leys et al., 2009; Tyler, 2003). It has been argued that sponge cell adhesion relies primarily upon an extracellular proteoglycan complex termed the Aggregation Factor (Bucior and Burger, 2004; Cauldwell et al., 1973; Grice et al., 2017; Haseley et al., 2001; Henkart et al., 1973; Humphreys, 1963; Vilanova et al., 2016). Antibodies raised against the Aggregation Factor have been reported to block reaggregation of dissociated cells (Schütze et al., 2001), and purified Aggregation Factor can mediate adhesion between beads in cell-free assays (Jarchow and Burger, 1998). Consequently, the integrity of sponge tissues is thought to depend upon the interaction of cells with the Aggregation Factor, an ECM component, rather than through cell junctions like those found in epithelia of other animals (Harwood and Coates, 2004; Varner, 1995).

The singular importance of the Aggregation Factor has been challenged by sequencing studies that have revealed conserved homologs of genes encoding adherens junction and focal adhesion proteins in diverse sponges (Fahey and Degnan, 2010; Nichols et al., 2006; Srivastava et al., 2010). Moreover, there is mounting experimental evidence that these proteins have conserved adhesion roles in sponge tissues. In the homoscleromorph sponge *Oscarella pearsei* (formerly *O. carmela*), a homolog of vinculin (common to adherens junction and focal adhesions in bilaterians) was detected at cell-cell and cell-ECM adhesions, and was found to interact with actin and talin *in vitro* (Miller et al., 2018). Also, a yeast two-hybrid screen revealed conserved interactions between the adherens junction component β-catenin with a classical cadherin (Nichols et al., 2012a). Likewise, in the freshwater sponge *Ephydatia muelleri*, both a classical cadherin and α-catenin were detected as co-precipitates of β-catenin (Emβ-catenin), which localized to actin plaques at cell-cell contacts that resemble adherens junctions (Figure 1) (Schippers and Nichols, 2018). These data indicate that adherens junction and focal adhesion proteins may have conserved functions in sponge tissues.

**Figure 1.**
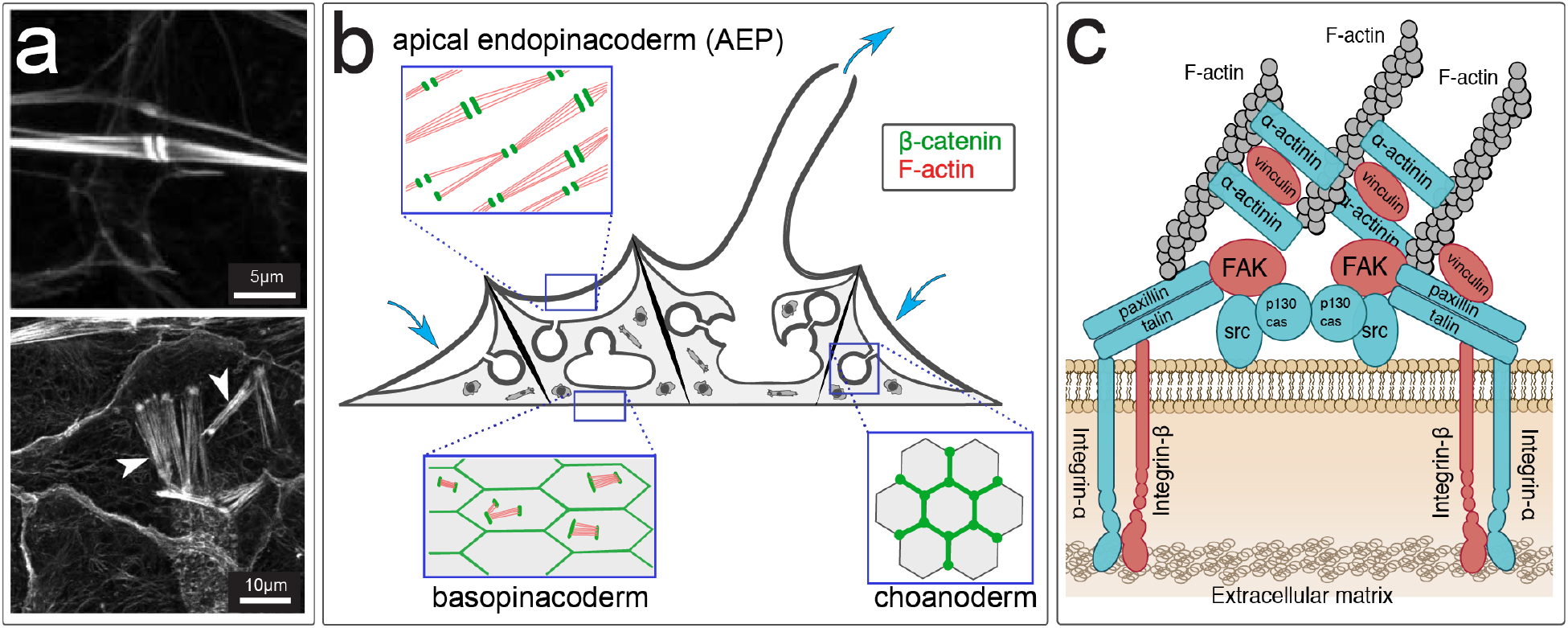
Adherens Junction- and Focal Adhesion-like structures in tissues of *Ephydatia muelleri*. (a) Top: Cells of the apical endopinacoderm contain bundles of actin filaments that culminate in dense plaques resembling spot adherens junctions at points of cell-cell contact. Bottom: Cells of the basopinacoderm contain bundles of actin filaments that resemble stress fibers of focal adhesions (white arrowheads). (b) Cross-sectional diagram of the juvenile *E. muelleri* body illustrating the distribution of Emβ-catenin (blue arrows indicate the direction of water flow in the aquiferous system). (c) Schematic illustration of the molecular organization of a focal adhesion. Proteins highlighted in red are the subject of the current study. (FAK = Focal Adhesion Kinase; Artwork in Panel b adapted from Schippers and Nichols (2018), and in Panel c adapted from Mitra et al. (2005).

However, the study of Emβ-catenin has also revealed new peculiarities of cell adhesion in sponges. Cells of the basopinacoderm (the tissue at the interface with the substrate) contain actin bundles that Max Pavans De Ceccatty (De Ceccatty, 1986) described as “devices for cell-to-substratum attachment.” If the mechanisms of cell-substrate adhesion in sponges are conserved with other animals, one might expect that these are integrin-based focal adhesions. Instead, they were found to stain positive for Emβ-catenin, an adherens junction component (Schippers and Nichols, 2018).

To better understand the composition and organization of cell junctions in sponge tissues, we examined the endogenous interactions and distribution of the focal adhesion proteins vinculin (Vcl), focal adhesion kinase (FAK) and integrin-β (ITGB) in *E. muelleri*. We found that this species has diverse adhesion structures composed of adherens junction and focal adhesion proteins, but these proteins are not as strictly partitioned to cell-cell versus cell-ECM junctions as they are in epithelial tissues of bilaterian animals. Instead, these proteins are often co-distributed in both contexts. Moreover, sponge tissues have specialized junctions not found in other animals; including cell-spicule junctions and cell-bacteria junctions. These data contribute to an increasingly complex narrative about the ancestral diversity and organization of cell junctions and their roles in early animal tissue evolution.

## Results

BLAST search (Altschul, 2014) of the *E. muelleri* transcriptome (Peña et al., 2016) revealed highly conserved homologs of the primary protein components of focal adhesions. We detected seven integrin-β homologs, six integrin-α homologs, two talin homologs, and one homolog each of vinculin, focal adhesion kinase, and paxillin (Supplement); integrins were numbered to reflect their relative expression levels, not to indicate their orthology to integrin subfamilies in other animals). We characterized the distribution of select focal adhesion proteins in sponge tissues by immunostaining with custom antibodies against EmVcl, EmFAK and EmITGB1. Each antibody was validated as specific to its target antigen under non-denaturing conditions (Supplement). Anti-EmITGB1 was also found to recognize EmITGB2 and EmITGB4. We subsequently refer to EmITGB generally, rather than EmITGB1 specifically, to reflect this cross-reactivity.

### Focal Adhesion-like structures may function in substrate attachment

To determine whether focal adhesion-like structures in the basopinacoderm (Figure 1) are involved in cell-substrate attachment, it was important to distinguish whether they form at the interface between the tissue and the substrate (coverslip), or between the tissue and the mesohyl (ECM-filled interior of the sponge). To test this we used Total Internal Reflection Fluorescence (TIRF) Microscopy (Axelrod, 1981; Fish, 2009) and found that junction-associated stress fibers (Livne and Geiger, 2016) were within 100-200 nm of the coverslip, near the substrate-adjacent cell membrane (Figure 2a) – consistent with a role in cell-substrate attachment. We further reasoned that shear forces associated with water flow and turbulence should lead to an increase in the number of focal adhesion-like structures, as mechanical tissue stress induces focal adhesion formation in cultured vertebrate cells (Davies et al., 1994). To test this we grew sponges in dishes on a rocking platform for comparison with sponges grown in dishes on a stable platform. As shown in Figure 2b, a 43% increase was detected in sponges grown on a rocking platform (n=12, p-value = 0.0058).

**Figure 2.**
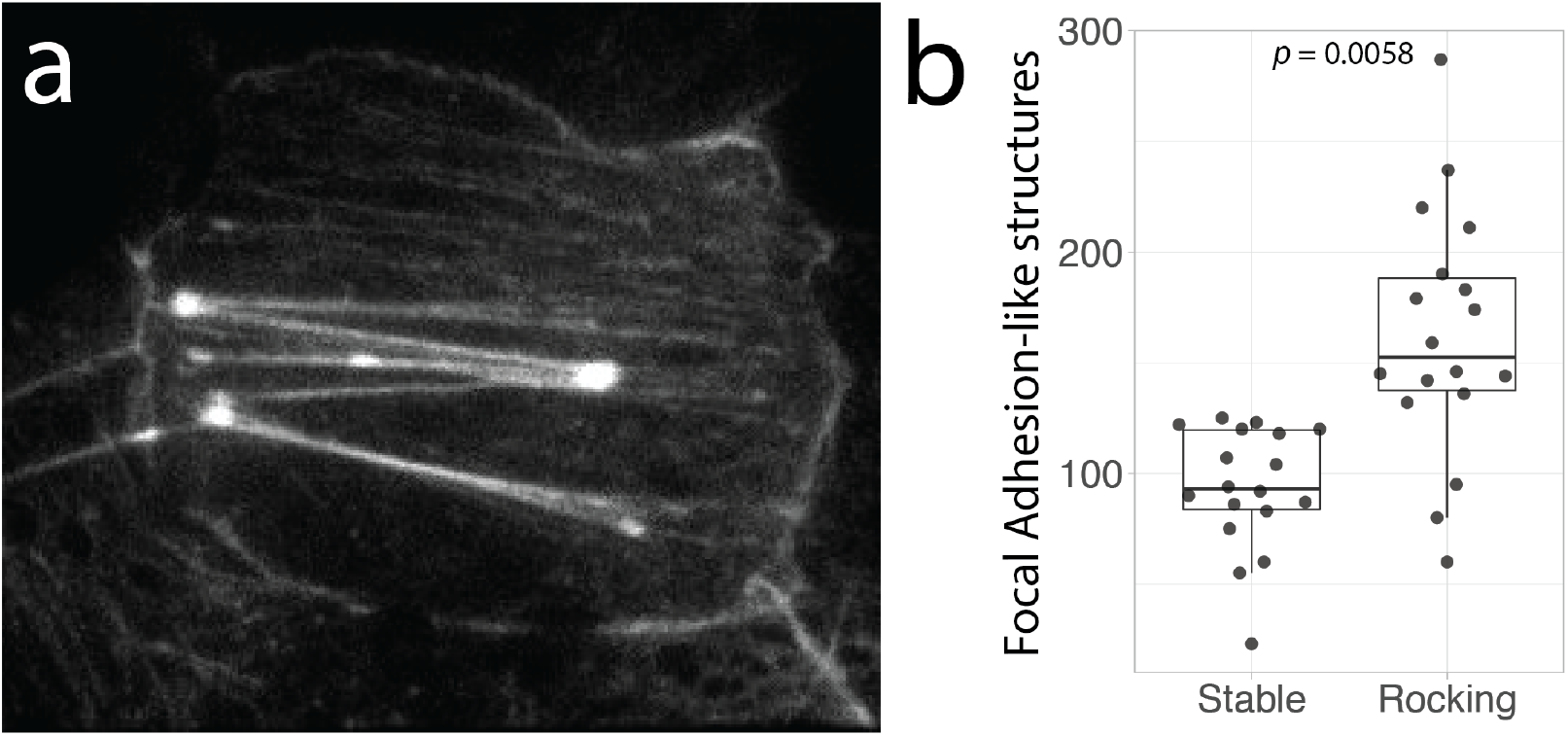
Actin stress fibers in the basopinacoderm may be associated with cell-substrate adhesions. (a) Total Internal Reflection Fluorescence (TIRF) imaging was used to determine the subcellular localization of focal adhesion-like structures in the basopinacoderm. (b) Their abundance was quantified in individuals grown on a stable surface and compared to individuals grown on a rocking platform.

### Three different types of focal adhesion-like structures in the basopinacoderm

Further examination of stress fiber-like structures in the basopinacoderm revealed three distinct categories (Figure 3). The first category included the actin filaments detected by TIRF at the substrate-adjacent cell membrane (Figure 3a), which we termed ‘ventral adhesions’. The second category resembled ventral adhesions, but with one or both ends terminating at a membrane invagination or vesicle containing bacteria (Figure 3b). We termed these ‘bacterial adhesions’. A third category of actin filaments was found to span vertically from the substrate-adjacent (ventral) cell membrane to the mesohyl-adjacent (dorsal) cell membrane. These actin filaments formed prominent plaques on the dorsal surface of the cell (Figure 3c) that we termed ‘dorsal adhesions’.

**Figure 3.**
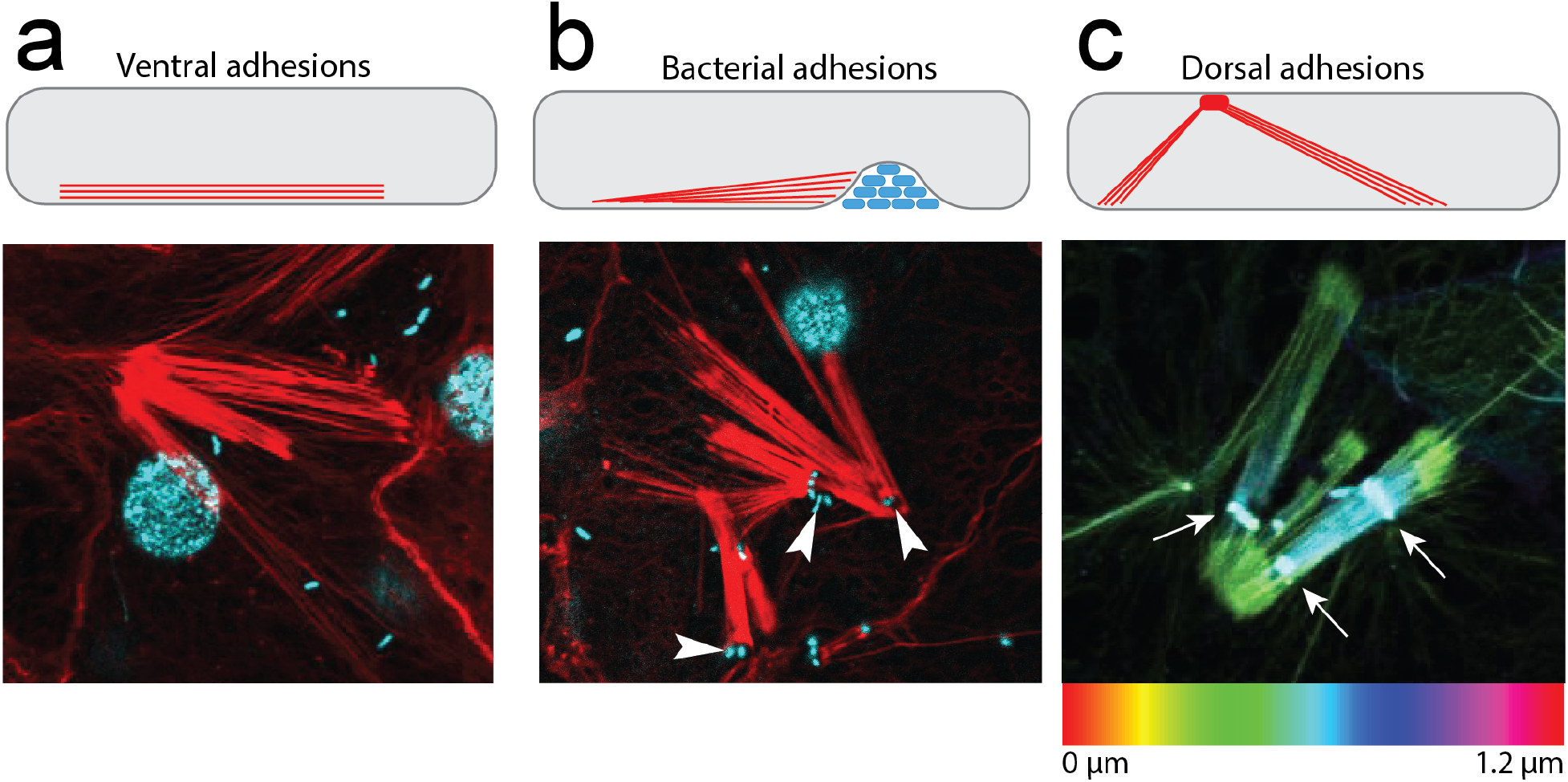
Three types of focal adhesion-like structures in basopinacoderm. (a-c) Basopinacocytes (cells of the substrate-attachment epithelium) are drawn at the top in profile view, with the mesohyl-interface (dorsal surface) at the top and substrate-interface (ventral surface) at the bottom. Bundles of actin filaments were found at the (a) ventral surface, and (b) were sometimes associated with membrane invaginations containing bacteria (white arrowheads). Panel (c) shows actin staining, colored to depict pixel depth within the confocal stack. White arrows indicate dorsal adhesions, from which actin filaments descend ventrally within cell. (panels a, b: red = actin; cyan = DNA)

Immunostaining of adhesion proteins at these three different categories of focal adhesion-like structures revealed that they were compositionally distinct. Only EmVcl was found to be associated with ventral adhesions (Figure 4), whereas both Emβ-catenin and EmVcl were consistently detected at bacterial adhesions (Figure 5), and only EmITGB was detected at dorsal adhesions (Figure 6). The staining patterns of EmFAK were inconsistent in the basopinacoderm, and difficult to discern due to high levels of cytosolic staining. For example, EmFAK was not usually detected at focal adhesion-like structures (Figures 4–6), except rarely at bacterial adhesions (Supplement).

**Figure 4.**
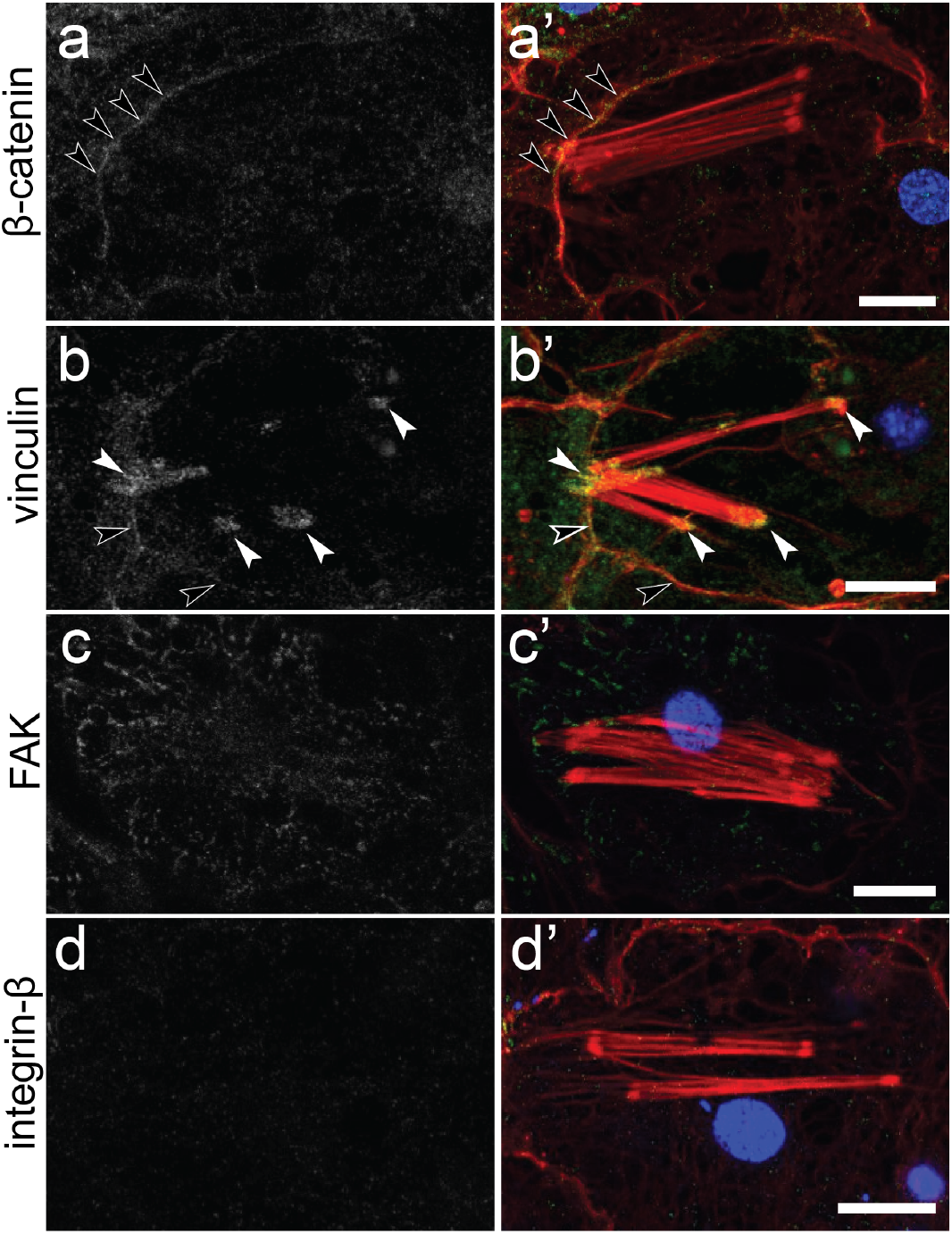
Immunostaining of ventral adhesions in the basopinacoderm. Both Emβ-catenin and EmVcl exhibited faint cell boundary staining (black arrowheads), but only EmVcl was detected in association with ventral adhesions (white arrowheads). [(a-d) antibody staining only; (a’-d’) antibody = green, DNA = blue, actin = red; scale bars = 10 µm].

**Figure 5.**
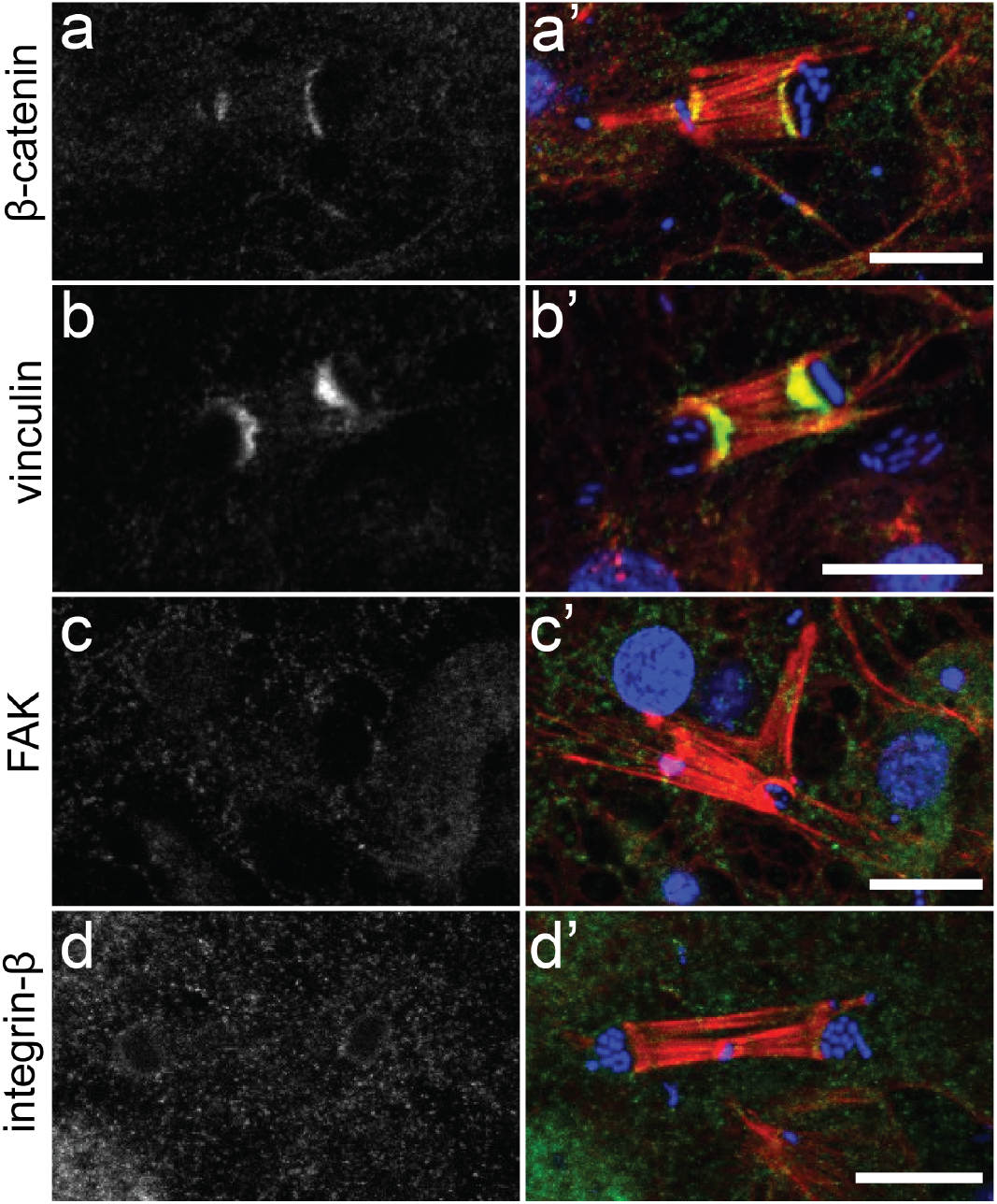
Immunostaining of bacterial adhesions in the basopinacoderm. Both (a) Emβ-catenin and (b) EmVcl were detected at the interface of stress fibers and membrane pockets containing environmental bacteria. Neither (c) EmFAK nor (d) EmITGB were detected at these structures (but see text for further discussion of EmFAK). [(a-d) antibody staining only; (a’-d’) antibody = green, DNA = blue, F-actin = red; scale bars = 10 µm].

**Figure 6.**
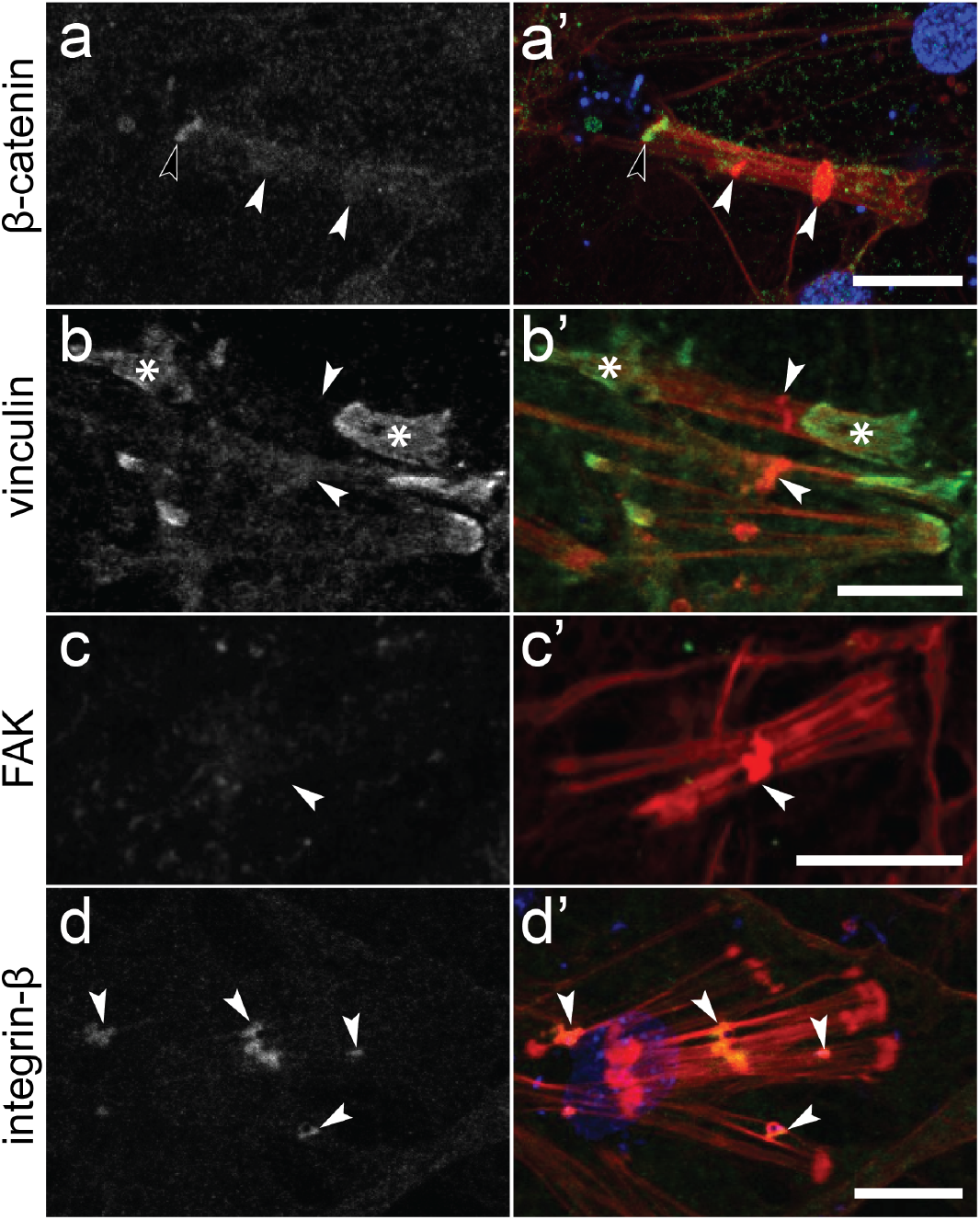
Immunostaining of dorsal adhesions. Neither (a) Emβ-catenin, (b) EmVcl, nor (c) EmFAK were detected at mesohyl-interfacing plaques (white arrowheads) of dorsal adhesion stress fibers. Visible staining of Emβ-catenin corresponds to a bacterial adhesions (black arrowhead) and of EmVcl corresponds to ventral adhesions (Asterisk). In contrast, (d) EmITGB was highly enriched at dorsal adhesions. [(a-d) antibody staining only; (a’-d’) antibody = green, DNA = blue, F-actin = red].

### Cell junctions at the spicule interface

A unique component of the ECM of many sponges are spicules; siliceous skeletal elements that form tent pole-like tissue supports. Specialized transport cells attach to spicules and move them into position (Nakayama et al., 2015) where they are anchored at rosette-shaped clusters of cells in the basopinacoderm by collagen (Nakayama et al., 2015).

We consistently detected cell junctions at the interface of cells and spicules, which stained positive for both Emβ-catenin and EmVcl. Again, EmFAK had a low signal-to-noise ratio, making it difficult to determine if it was present at these structures. EmITGB was not detected (Figure 7). It was unclear if the cells forming these structures were transport cells, basopinacocytes, or a different cell type altogether.

**Figure 7.**
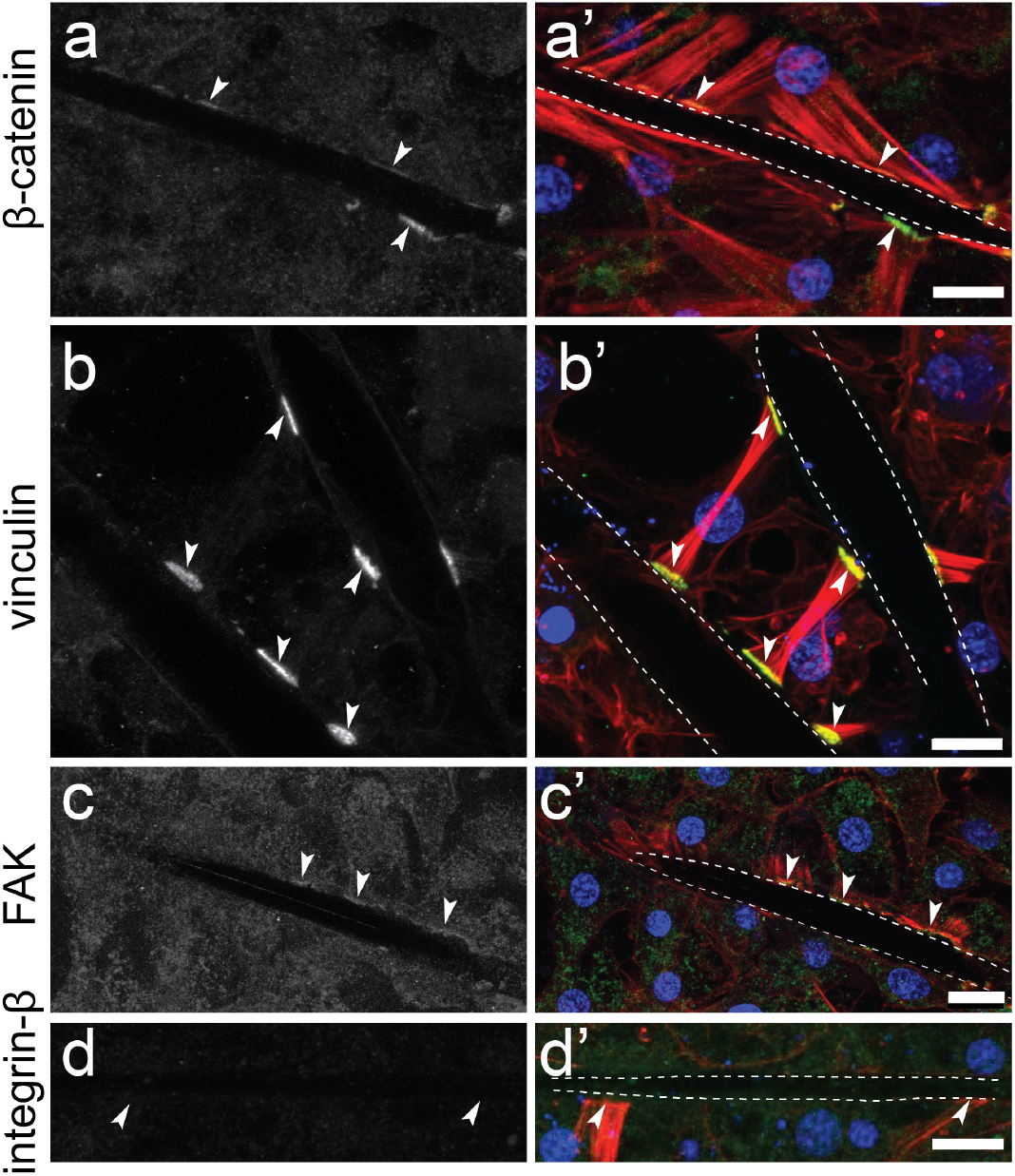
Immunostaining of adhesion proteins at cell-spicule junctions. (a) Emβ-catenin and (b) EmVcl localize to cell junctions at the interface with spicules (dotted lines mark spicules). (c) EmFAK was possibly enriched at these structures, but only marginally above background levels (arrowheads), whereas (d) EmITGB was not detected at all. [(a-d) antibody staining only; (a’-d’) antibody = green, DNA = blue, F-actin = red; scale = 10 µm].

### No evidence for focal adhesion-dependent cell migration

Focal adhesions have well characterized roles in the migration of cultured vertebrate cells, where they provide traction needed for movement across two-dimensional surfaces. Cell migration through three-dimensional ECM is often less dependent on integrin-mediated adhesion (Paluch et al., 2016). Sponges have dense populations of migratory cells in the three-dimensional environment of the mesohyl (Supplement), and we previously showed that filopodia of migratory cells in the sponge *O. pearsei* stain positive for vinculin (Miller et al., 2018). In contrast, neither EmVcl, EmFAK nor EmITGB were detected in migratory cells of *E. muelleri* (Supplement).

### Both Emβ-catenin and focal adhesion proteins are present at cell-cell junctions

We previously reported cortical staining of Emβ-catenin in the basopinacoderm, choanoderm, and the apical endopinacoderm (the inner tissue layer of the sponge surface; see Figure 1) (Schippers and Nichols, 2018). As shown in Figure 4a, Emβ-catenin was again detected at the cell cortex in the basopinacoderm, but we also detected EmVcl (Figure 4b) and less frequently EmITGB (Supplement). This staining was generally of low intensity and patchy, and we wondered whether this might reflect the developmental stage of the immature juvenile tissues examined. To test this we grew sponges for an additional three weeks and found markedly elevated levels of cortical staining and robust adherens junction-like structures that were EmVcl-positive (Supplement).

We detected focal adhesion proteins at cell-cell junctions in other tissues, as well. The most conspicuous cell-cell junctions in *E. muelleri* are found in the apical endopinacoderm at points where actin tracts align between neighboring cells (Figure 1b). Emβ-catenin was previously detected at these structures (Schippers and Nichols, 2018), and we found that EmVcl, EmFAK and EmITGB were also constitutively present (Figure 8). Cortical staining of EmITGB was also detected in an adjacent tissue, the exopinacoderm (Figure 8d) – this is the outermost sponge tissue and is so close in proximity to the apical endopinacoderm that they cannot be separately resolved by confocal microscopy.

**Figure 8.**
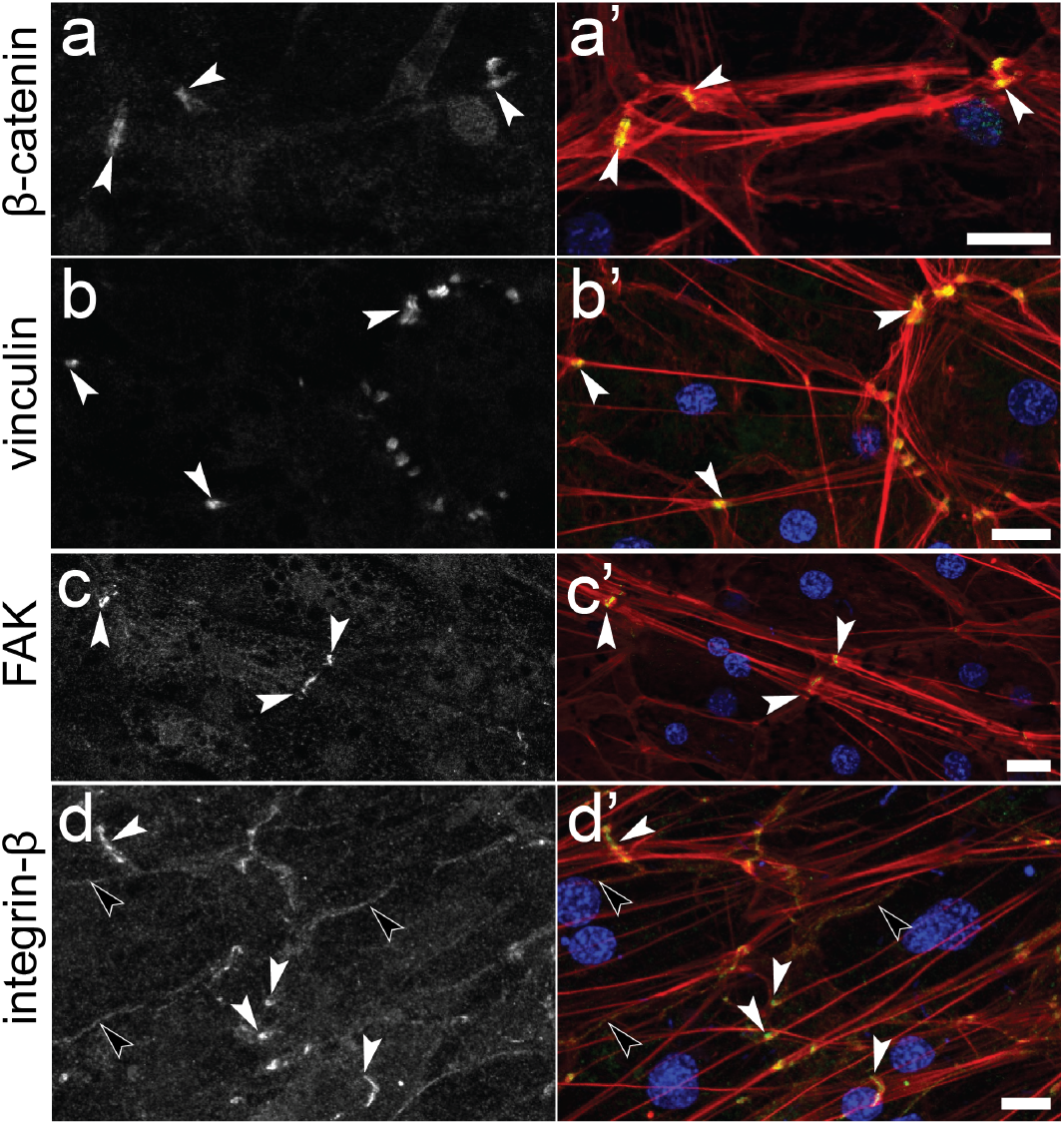
Immunostaining of cell-cell junctions in the apical endopinacoderm. Emβ-catenin, EmVcl, EmFAK and EmITGB were detected at probable adhesion plaques where F-actin tracts align between adjacent cells (white arrowheads). Low intensity staining of EmITGB was also detected at the cell cortex (black arrowheads). [(a-d) antibody staining only; (a’-d’) antibody = green, DNA = blue, F-actin = red; scale = 10 µm].

The detection of EmVcl and EmFAK at cell-cell junctions is not unprecedented. Applied force on E-cadherin has been shown to lead to phosphorylation of human vinculin at Y822 and recruitment to the adherens junction (Bays et al., 2014; Mui et al., 2016). However, alignment of EmVcl to human vinculin revealed low conservation in the region that contains Y822 (Supplement), making it difficult to predict if this mechanism for regulating vinculin function is conserved in *E. muelleri.* But, EmVcl was not detected as a co-precipitate of Emβ-catenin (Schippers and Nichols, 2018), nor did we detect adherens junction proteins as co-precipitates of EmVcl (Supplement).

Like vinculin, focal adhesion kinase has also been reported to function in contexts other than focal adhesions, including at the adherens junctions of vascular endothelia, where it directly binds VE-cadherin and phosphorylates β-catenin in response to VEGF activation (Chen et al., 2012). However, EmFAK was not detected as a co-precipitate of Emβ-catenin (Schippers and Nichols, 2018), whereas it did co-precipitate with EmITGB (Supplement). We treated sponges with 5µM FAK inhibitor 14 and found that this treatment abolished FAK staining at cell-cell junctions, but detected no other effects on the formation or molecular composition of adhesion structures in the apical endopinacoderm (Supplement).

The only tissue where we did not find evidence of co-distributed adherens junction- and focal adhesion-proteins at cell-cell contacts was the choanoderm (Supplement). In this tissue, Emβ-catenin alone was detected (Schippers and Nichols, 2018).

### Focal adhesion proteins co-precipitate

The localization of both adherens junction and focal adhesion proteins to cell-cell junctions, cell-spicule junctions, and bacteria-associated stress fibers raised the question of whether they interact directly, in ways not characterized from other animals. To test this, we analyzed the immunoprecipitates of EmVcl, EmITGB, and EmFAK using LC-MS/MS to identify their endogenous binding partners. EmITGB was found to co-precipitate with EmFAK, EmTalin2, EmITGA1, EmITGA2, and EmITGA3; all homologs of proteins known to function at integrin-based focal adhesions (Horton et al., 2016). Notably, EmVcl essentially had no co-precipitating proteins (it was the most abundant protein in the precipitate by nearly two orders of magnitude), and neither EmVcl nor EmPaxillin (a constitutive focal adhesion component in bilaterian animals) were detected in EmITGB precipitates.

## Discussion

Most animal cell adhesion proteins evolved early, concurrently with or before the transition to multicellularity (Abedin and King, 2010, 2008; Belahbib et al., 2018; Dickinson et al., 2011; Fahey and Degnan, 2010; Harwood and Coates, 2004; Nichols et al., 2012b, 2006; Sebé-Pedrós et al., 2010). However, our understanding of how these proteins functioned ancestrally, and when they were organized into interacting complexes (i.e., cell junctions) is limited. We examined the interactions and distribution of focal adhesion proteins in tissues of *E. muelleri*. We found that they co-precipitate as a complex from cell lysates and that EmVcl, EmFAK and EmITGB localize to apparent cell junctions, supporting that they have conserved adhesion roles. However, we also detected a critical difference in the spatial distribution of these proteins in sponge tissues compared to epithelia in other animals. Rather than being restricted to focal adhesion-like structures at cell-ECM contacts, they were also detected at adherens junction-like structures at cell-cell contacts, and were often co-distributed with the adherens junction protein Emβ-catenin.

In a prior study, we reported a similar anomaly: Emβ-catenin localizes to focal adhesion-like structures in the basopinacoderm of *E. muelleri (Schippers and Nichols, 2018)*. Here, we found that these structures can be parsed into structurally and compositionally distinct categories: ventral, dorsal, and bacterial adhesions. When taken into consideration, we found that Emβ-catenin was not actually associated with either ventral or dorsal adhesions. Instead, these structures respectively stained positive for EmVcl and EmITGB, consistent with their homology to focal adhesions in bilaterian tissues. The absence of EmITGB staining at EmVcl-positive adhesions does not necessarily indicate the absence of integrins altogether, as our antibody recognized only three of seven identified paralogs. In the future, the focal adhesion protein talin may serve as a more universal marker of integrin-distribution, as it is constitutively present at all integrin-based adhesions in other animals, and was detected as a co-precipitate of EmITGB from *E. muelleri* lysates.

The previously reported Emβ-catenin staining at focal adhesion-like structures in the basopinacoderm is actually restricted to bacterial adhesions. In this context, Emβ-catenin is co-distributed with the focal adhesion protein homologs, EmVcl and (sometimes) EmFAK. The functional significance of these junctions is unknown, but intriguing. In natural environments, bacterial biofilms are abundant and often provide settlement cues for the larvae of aquatic animals, including sponges (Whalan and Webster, 2015). Thus, it seems plausible that bacterial adhesions could be involved in environmental sensing through integrin-mediated signaling. An alternative possibility comes from the observation that the bacteria at these structures are encapsulated in membrane invaginations or vesicles. Most sponge cells are phagocytic (Wilkinson et al., 1984), and perhaps bacterial adhesions are involved in phagocytosis. This could be an undescribed mode of feeding, or function in the uptake of intracellular symbionts or pathogens. There is precedent for such a mechanism in vertebrates, where integrins are known to be involved in phagocytosis of particles, including microorganisms as part of a pathogen defense system (Dupuy and Caron, 2008; Underhill and Ozinsky, 2002). A difference is that β-catenin is not detected at these structures in vertebrates.

In the future, it will be interesting to identify the bacterial species at these structures, track their fate in sponge cells, and test for bacterial adhesions in attached larvae undergoing metamorphosis. Animals evolved in an environment dominated by bacteria (Alegado and King, 2014), and a compelling hypothesis is that cell adhesion molecules may have first evolved to mediate interactions with bacteria (Abedin and King, 2008).

A mixture of adherens junction and focal adhesion proteins was also detected in adhesion contexts other than bacterial adhesions. Specifically, focal adhesion proteins were co-distributed with Emβ-catenin at cell-spicule junctions, which essentially have the same composition as bacterial adhesions, and cell-cell junctions. The latter had previously been interpreted as probable adherens junctions (Schippers and Nichols, 2018), but until the role of focal adhesion proteins (particularly integrins) at these structures is clarified, this conclusion is less certain.

The co-distribution of Emβ-catenin with EmVcl, EmFAK and EmITGB could indicate that these proteins are part of a common adhesion complex in sponges tissues, but our immunoprecipitation results do not support this view. Focal adhesion proteins co-precipitate as a complex, just as adherens junctions proteins were found to co-precipitate (Schippers and Nichols, 2018); each to the exclusion of the other. Also, Emβ-catenin alone was detected at cell-cell contacts in the choanoderm, and focal adhesion proteins alone were detected at ventral and dorsal adhesions in the basopinacoderm. This indicates that these protein complexes are functionally separable and may have discrete roles, even where they are co-distributed.

Full characterization of sponge cell junctions will require further identification of associated adhesion receptors (e.g., cadherins), and integration of these data with models of Aggregation Factor-mediated cell adhesion. One clue to how these adhesion systems may interact is that the Aggregation Factor has reported RGD motifs, leading to the hypothesis that it may activate integrin signaling (Fernàndez-Busquets et al., 1998; Harwood and Coates, 2004). But, the Aggregation Factor has predominantly been studied *in vitro*, in cell dissociation/re-aggregation assays. Its distribution is not well-characterized in intact tissues.

An important consideration is that, until recently, hypotheses about the evolutionary origin of animal cell adhesion mechanisms have been inadvertently biased toward bilaterian models. As we begin to examine cell adhesion in non-bilaterian lineages, there appears to be more mechanistic diversity than anticipated. For example, the cnidarian *Nematostella vectensis* has a conserved classical cadherin/catenin complex (Clarke et al., 2016), but β-catenin is not always detected at cadherin-positive cell-cell adhesions in tissues (Pukhlyakova et al., n.d.; Salinas-Saavedra et al., 2018). Likewise, sequence analyses of ctenophores indicate that they lack conserved cadherin/β-catenin interaction motifs (Belahbib et al., 2018), and a recent study indicates that β-catenin is altogether absent at cell-cell contacts in *Mnemiopsis leidyi* (Salinas-Saavedra et al., n.d.). The molecular composition of cell junctions in placozoans is entirely uncharacterized, but from an ultrastructural perspective they resemble adherens junctions (Smith and Reese, 2016). Placozoans altogether lack cell-ECM junctions and a basal lamina (Hynes, 2012). A comprehensive understanding of the timing and sequence of cell junction assembly and the evolution of epithelia will require detailed studies of adhesion in diverse non-bilaterian tissues.

### Conclusions

This study supports that adherens junction and focal adhesion proteins functioned in adhesion and tissue organization in the last common ancestor of sponges and other animals. This stands in apparent contrast to studies which have emphasized the Aggregation Factor as the predominant adhesion mechanism in sponges. Sponge tissues are organized much more like epithelia in other animals than previously appreciated.

However, in contrast to cell adhesion properties that sponges share in common with other animals, we also discovered new ways in which they differ. Adherens junction and focal adhesion proteins are not always spatially partitioned into compositionally distinct cell-cell and cell-ECM junctions; rather, they are often co-distributed. This complicates narratives about the origins of these cell junctions and their ancestral roles in tissue organization.

Finally, the discovery of specialized cell-bacteria junctions raises new questions about the functional significance of these structures for sponge physiology (environmental sensing, feeding, symbiosis, or pathogen defense), and possibly about the ancestral role of cell adhesion molecules in animals. It is widely acknowledged that animals evolved in a bacteria-rich environment. If the interaction of cell adhesion proteins with bacteria is an ancient feature of animal biology, bacterial adhesions in sponges may provide clues to the nature of these interactions.

## Materials and Methods

### Identification of focal adhesion protein homologs in E. muelleri

Representative sequences of the focal adhesion proteins integrin-α, integrin-β, vinculin, talin, focal adhesion kinase and paxillin were retrieved from Uniprot (UniProt Consortium, 2019) and used to query the *Ephydatia muelleri* transcriptome (Peña et al., 2016) by BLAST search (Altschul, 1990) to identify candidate sponge homologs. The putative domain composition of *E. muelleri* sequences was then annotated using HMMER (Finn et al., 2011) and SMART (Letunic and Bork, 2018) web-servers. *E. muelleri* vinculin was previously distinguished from its close paralog α-catenin by phylogenetic analysis (Miller et al., 2018).

### Specimens

*Ephydatia muelleri* gemmules were collected from “upper” Red Rock Lake, Colorado, USA (40.0802, −105.543) in early October. This lake is several hundred meters southwest of Red Rock Lake, Boulder County and is unnamed. Gemmules were stored in autoclaved lake water, in the dark at 4°C. Before plating, gemmules were washed in 1% hydrogen peroxide for 5 minutes, washed three times in autoclaved lake water and grown at room temperature.

### Cloning and Recombinant Protein Expression

The coding sequence of target antigens were amplified by polymerase chain reaction (PCR) from an *E. muelleri* cDNA library using Phusion High-Fidelity DNA polymerase (NEB). Primer sequences and amplicons are specified in the Supplement (Supplement). PCR products were cloned into pET28a (Novagen), pET28 His6 Sumo TEV LIC (1S) #29659 or pET His6 GST TEV LIC (1G) #29655 (Addgene) for expression.

Expression constructs were validated by Sanger Sequencing (Eurofins) and transformed into a protease deficient *Escherichia coli* strain (Rosetta 2(DE3), Promega). For expression, a single colony was grown in Luria Broth at 37 ºC to an OD_600_ between 0.4-0.6, then induced with 300 mM of isopropyl-1-thio-β-d-galactopyranoside (IPTG) for 3-5 hours at 30 ºC. Bacterial pellets were collected by centrifugation, resuspended in 1X PBS pH 7.4 on ice. Cells were lysed by the addition of 1 mg/ml lysozyme and 0.2 mM phenylmethanesulfonyl fluoride (PMSF), incubation at room temperature (RT) for 15 min, then sonication for 4 x 30 seconds. Bacterial debris was removed by centrifugation and the supernatant incubated with either with HisPur Cobalt or Nickel Resin (Thermo Fisher Scientific) for His-tagged proteins or GST-agarose resin (Thermo Fisher Scientific) for GST-tagged proteins, for ~18 h at 4°C on a tube rotator. The resin was collected by centrifugation, and washed in either 1X PBS pH 7.4 (His-tagged proteins) or 50 mM Tris, 1M NaCl, pH 8.0 (GST-tagged proteins). After washing, purified recombinant protein was eluted by the addition of either 150 mM imidazole (His-tagged proteins) or 10 mM reduced glutathione (GST-tagged proteins).

### Antibody Production

Polyclonal antibodies were generated in rabbit against His-EmVcl, His-EmFAK and GST-EmITGB1 (Syd Labs) recombinant proteins. For affinity purification, two columns were made: 1) whole *E. coli* lysates, and 2) 6-10 mg of recombinant protein. Each were covalently bound to 1 mL of AminoLink Plus Coupling resin (Thermo Scientific, Cat#20501) according to manufacturer specifications. Anti-sera were passed over the *E. coli* column to remove antibodies against bacterial proteins, then the flow-through was incubated with the antigen-coupled resin for 1 h at RT under rotation. This column was washed with 12 mL of AminoLink Wash Buffer and the antibodies were eluted with 500 µL 0.1 M glycine HCl, pH 2.5. The pH of eluted fractions was adjusted to neutral by adding 30 µL of 0.75 M Tris-HCL pH 9.0. Antibody titer was quantified by spectrophotometry (A280) and by visual comparison to BSA standards via sodium dodecyl sulfate-polyacrylamide gel electrophoresis (SDS/PAGE). The specificity of each antibody was validated by Western Blot, Immunoprecipitation coupled with LC-MS/MS, and by pre-adsorption with the injected antigen prior to immunostaining. Validation details and results are available in the Supplement.

### Western Blot

For each Western Blot, ~100 gemmules were grown in petri dishes with lake water containing 100 µg/mL ampicillin for 6-13 days at RT. Juveniles were scraped with a razor into 4X SDS-PAGE reducing loading buffer (1M Tris, pH 7.0, 20% SDS, 20% Glycerol, 0.02% bromophenol blue and 2.5% β-mercaptoethanol), vortexed and boiled at 95ºC for 3 min. Proteins were separated by SDS-PAGE on a 10%-12% gel, and transferred to a PVDF membrane (Millipore) at 350 mAmp for 30 min. Membranes were blocked for 1 h at RT in 5% nonfat milk in 1x PBST, pH 7.4 (0.05% Tween 20) and then incubated with affinity purified antibodies (1mg/mL stocks) against EmVcl (1:3,000), EmFAK (1:1,000) and EmITGB (1:1,500), in blocking solution for 1 h at RT and washed twice in 1x PBST pH 7.4. After 45 minutes of incubation with the secondary antibody (Alexa488® Goat Anti-Rabbit IgG Antibody; Life Technologies, 1:1000 dilution) at RT, membranes were washed in 1x PBST pH 7.4 and imaged using the Molecular Imager FX ProPlus (BioRad).

### Immunoprecipitation and Mass Spectrometry

Affinity purified antibodies were coupled to agarose A/G using the Pierce Crosslink CoIP Kit (Thermo Scientific Cat #26147). A control IP was performed using rabbit IgG (I5006, Sigma-Aldrich). For EmVcl, cell lysates were prepared by combining 1.1 mg of frozen adult tissues with 1.8mL Pierce Lysis buffer (Thermo Fisher Scientific) containing Complete Mini Protease Inhibitor Cocktail (Roche, EDTA-free), Aprotinin and Leupeptin (1 mM). Lysates for EmFAK and EmITGB1 IPs were prepared by scraping ~300 week-old sponges into 1 mL Triton Lysis Buffer (TLB; 20 mM HEPES, pH7.4, 150 mM NaCl, 1 mM ethylene glycol tetraacetic acid (EDTA), 10% Glycerol, 1% Triton X-100, 1 mM PMSF, 1 mM DTT, protease Inhibitor cocktail (ROCHE), 1mM Aprotinin, 1mM Leupeptin). Different tissue sources and lysis buffers were used as the technique was optimized over the course of the project. Samples were vortexed 15 seconds and returned to ice for 2 minutes; this was repeated 3 times. Samples were further then homogenized by hand (Argos Tech. A0001) for 30 seconds, and debris and gemmules were removed via centrifugation at 13,000 x*g* for 10 min at 4 ºC. 350 µL of the lysate was diluted with an additional 200 µL of lysis buffer and combined with the antibody-coupled agarose at 4ºC for 1.5 h. After collecting the flow-through (FT) and completing the washes recommended by the manufacturer, an extra wash using 1 M LiCl solution was performed to remove any non specific proteins. Finally, precipitates were eluted with Pierce Low pH Elution Buffer (Cat#21004, ThermoFisher) and neutralized with 1M Tris-HCl pH 9.0 (Cat#42020208-1, Bioworld). 20-25 µL aliquots of these precipitates were mixed with 5 µL 4x SDS-PAGE loading buffer containing freshly added 20% beta-mercaptoethanol (BME), boiled for 3 min, then loaded on a 12% gel for SDS-PAGE. One gel was used for Coomassie staining, and a replicate was analyzed by Western blot.

EmVcl precipitates were directly sent for further analysis by LC-MS/MS. EmFAK and EmITGB1 precipitates were excised from an SDS-PAGE gel to separate the precipitate from the co-eluted antibody. LC-MS/MS was performed by the Proteomics Core Facility, University of California, Davis for EmVcl and CU-Anschutz Proteomics Core Facility for EmFAK and EmITGB1. Results were analyzed using the software Scaffold (v3.1).

### Immunostaining

*E. muelleri juveniles* were grown from gemmules for 5-7 days on No. 1.5 uncoated dishes (MatTEK) or on glass coverslips. Tissues were fixed in 4% Formaldehyde in 95% cold EtOH for 30 min - 1 h at 4ºC. Juveniles were then washed three times with 1x PBS pH 7.6, and incubated in blocking buffer (3% BSA in 1x PBST pH 7.4) overnight at 4ºC. All antibody preparations were titrated to determine their optimal working concentration, from 1:250 - 1:5000. After incubation, samples were washed three times with 1x PBST and then incubated for 45 min with secondary antibody (Alexa488® Goat Anti-Rabbit IgG Antibody; Life Technologies, 1:500 dilution), plus Alexa Fluor568® Phalloidin (Life Technologies, 1:80) and Hoechst (33342, 1 µg/mL) at RT. Samples were washed once in 1x PBST and twice in 1x PBS pH 7.6 and preserved for imaging using anti-fade mounting medium (0.1M Propyl gallate, 1x PBS pH 7.6 and 90% glycerol). Confocal Images were acquired on an Olympus Fluoview FV3000 confocal laser scanning microscope using either a 20x/0.85 NA, 60x/1.4 NA or 100x/1.4 NA objectives, and processed using FIJI (Schindelin et al., 2012). Neither brightness nor contrast was adjusted in the antibody channel. Immunostaining results were validated by secondary-only control and by pre-incubating each antibody with its corresponding antigen for at least 1 h at 4ºC before staining (Supplement).

### Quantification of Focal Adhesion Abundance

Single gemmules (n=12) from *E. muelleri* were placed in 3 mL of lake water in No. 1.5 uncoated dishes (MatTEK) and allowed to attach (3 days). After attachment half were transferred to a rocking platform for 24 hours, while the others were left on a steady surface. All individuals were fixed and stained with anti-EmVcl and phalloidin, and the basopinacoderm imaged as described. Focal adhesion-like structures were counted in each of three image stacks per individual and their abundance in each treatment was analyzed using a one-way ANOVA with single factor of treatment in R studio (RStudio Team, 2015).

### Pharmacological inhibition of FAK

Five day-old-juveniles were treated with 5 µM FAK Inhibitor 14 (Sigma-Aldrich) for four hours at RT in the dark. Treatment was removed and tissues were immediately fixed, immunostained, and imaged as described. The effects of FAK inhibition on cell motility are described in the Supplement.

## Supporting information

Main Supplement

## Acknowledgements

This work was supported by NASA grant 16-EXO16_2-0041 (S.A.N.). The authors would like to thank Jeff Colgren for thoughtful discussions and critical input throughout the course of the project.

