## Supplementary material for "Diverse cell junctions with unique molecular composition in tissues of a sponge (Porifera)": Main Supplement

**Supplement Table of Contents**

Supplemental File 1: FASTA file containing DNA sequences for *E. muelleri* focal adhesion homologs
Supplemental File 2: FASTA file containing peptide sequences for *E. muelleri* focal adhesion homologs
Supplemental File 3: Annotated EmVcl immunoprecipitation results
Supplemental File 4: Annotated EmITGB1 immunoprecipitation results
Supplemental File 5: Annotated EmFAK immunoprecipitation results
Supplemental File 6: Video of migratory cells in the mesohyl

Supplemental Figure 1: Antibody validation by immunoprecipitation/LC-MS
Supplemental Figure 2: Antibody validation by immunostaining preadsorption assay
Supplemental Figure 3. EmFAK was rarely detected at Bacteria-Associated Stress Fibers
Supplemental Figure 4: Migratory cells form focal adhesion like structures in two-dimensional, but not three-dimensional environments
Supplemental Figure 5: EmITGB localizes to cell boundaries in the basopinacoderm
Supplemental Figure 6: Robust cell-cell junctions in the basopinacoderm of older sponges
Supplemental Figure 7: Alignment of Y822 region of *E. muelleri* and human vinculin
Supplemental Figure 8: FAK inhibitor 14 effects on cell-cell junction composition
Supplemental Figure 9: Focal adhesion proteins are not detected in the choanoderm

Supplementary Results: Antibody Validation (Western Blot and Immunoprecipitation)

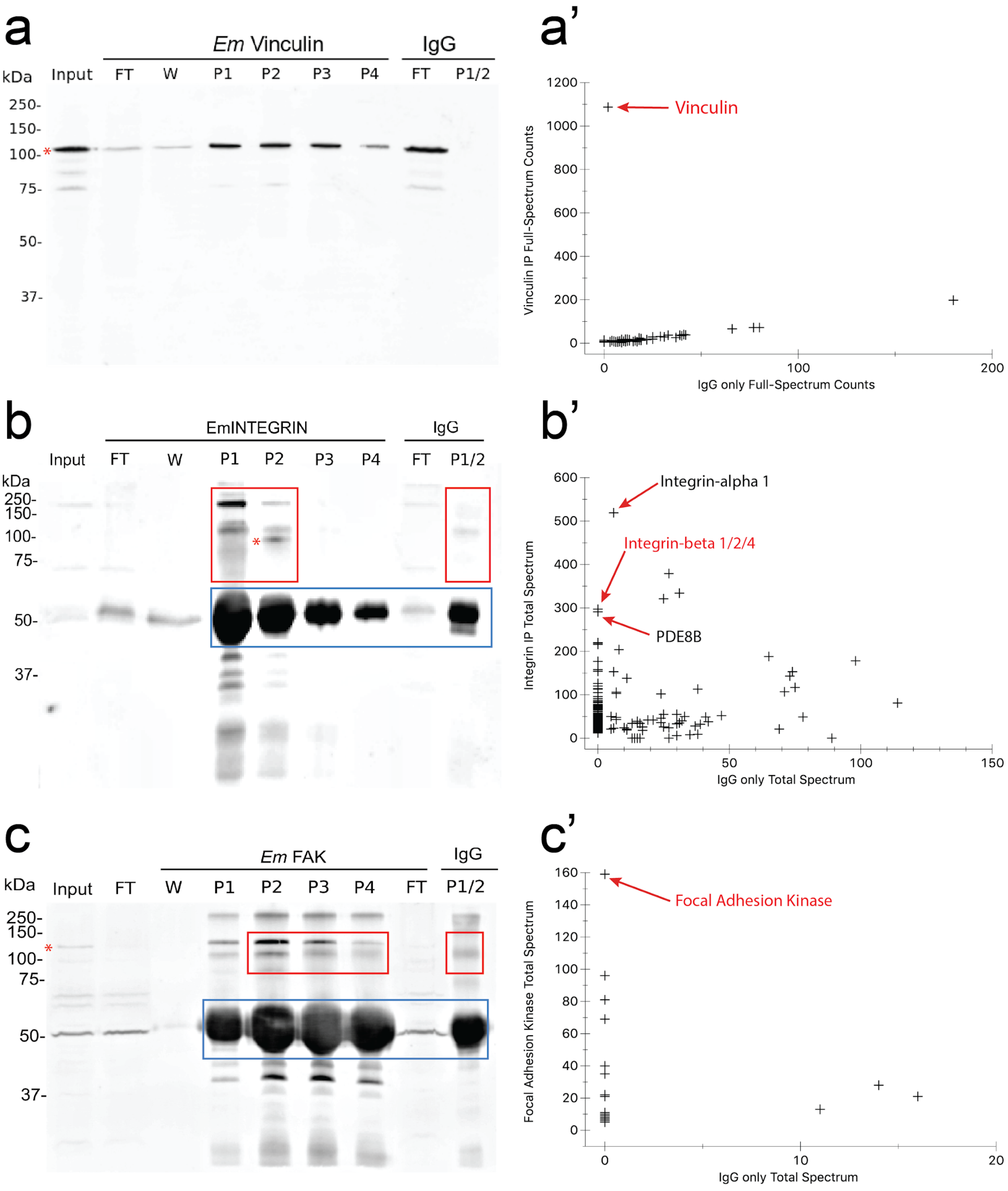


**Supplemental Figure 1: Antibody validation by immunoprecipitation/LC-MS.**
(a-c) Western blot analysis of immunoprecipitation samples. Bands corresponding to the predicted molecular weight of each target protein are indicated with a red asterisk. Anti-EmITGB and Anti-EmFAK lost activity upon cross-linking to the resin, so instead precipitates were co-eluted with the antibody and SDS-PAGE gel slices excluding the antibody fraction were

analyzed by LC-MS (red boxes = gel fraction analyzed by LC-MS; blue boxes = antibody heavy chain). Equivalent gel slices were sent from the IgG negative control sample. (a'-c') Scatter plots showing the abundance of *E. muelleri* proteins detected in each precipitate relative to the IgG control. The proteins with greatest abundance in each precipitate are indicated, with the target antigen highlighted in red. All samples were filtered to reflect only those hits within the 95% confidence interval, and represented by at least 5 unique peptides. See **Supplementary Results** section for further details. (input = whole-cell *E. muelleri* lysates, FT = lysate flow through/unbound fraction, W = 1M LiCl wash, P1-P4 = precipitate fractions 1-4).

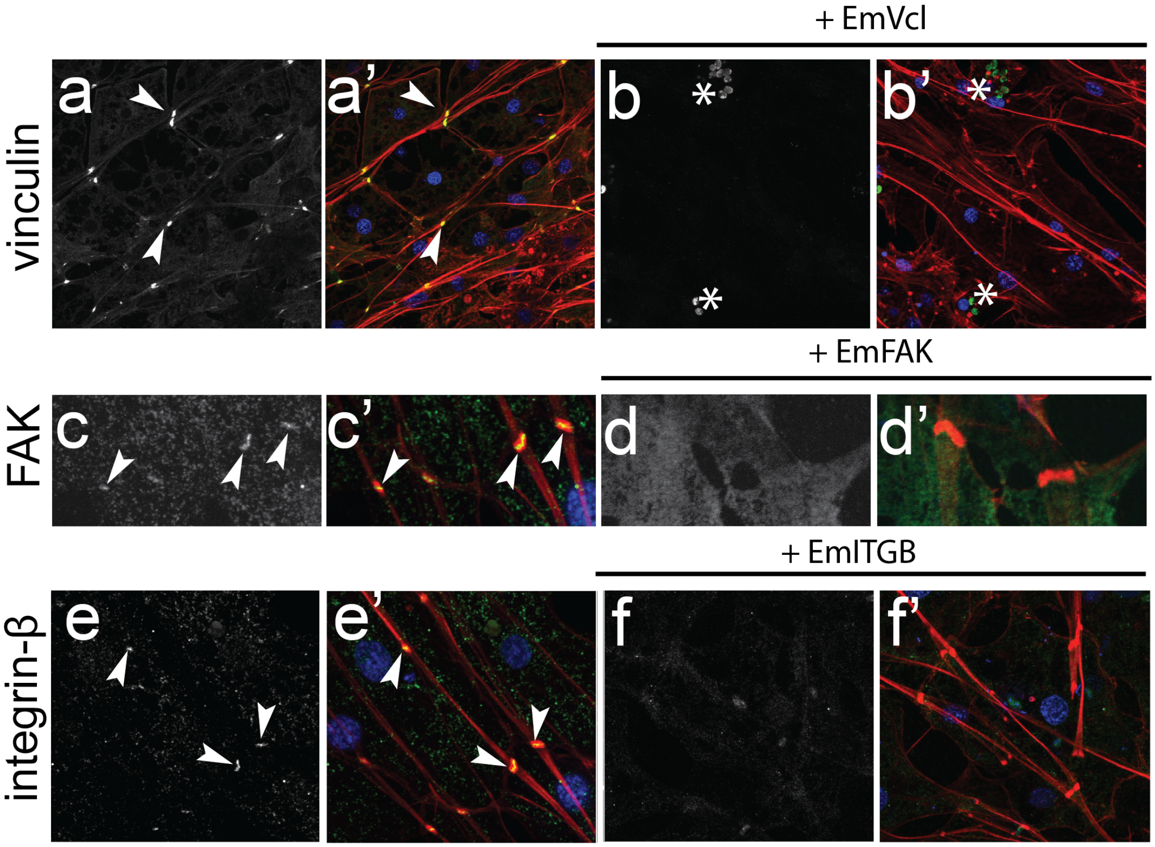


**Supplemental Figure 2: Antibody validation by immunostaining preadsorption assay.** Immunostained cell-cell junctions in the apical endopinacoderm using control (left panels) versus pre-adsorbed antibody aliquots. Preadsorption with 1-10 μg of each recombinant antigen completely abolished observed staining patterns, supporting the specificity of the immunostaining signal. [a-f = antibody only; a’-f’ = F-actin (red), DNA (blue) antibody (green)].

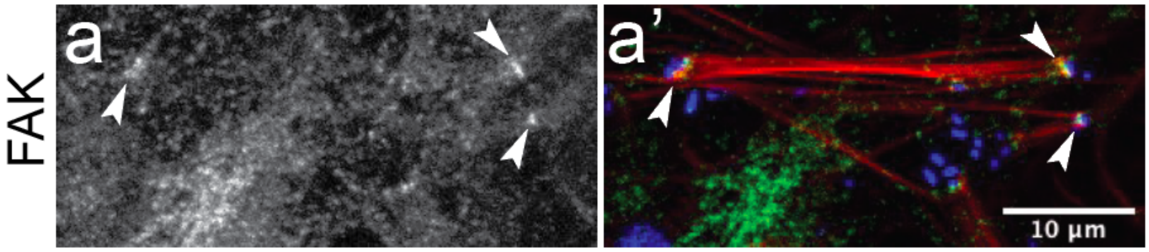


**Supplemental Figure 3: EmFAK was rarely detected at Bacteria-Associated Stress Fibers**
Whereas EmFAK was not routinely detected at focal-adhesion like structures in the basopinacoderm, it was sometimes present at the distal ends bacteria-associated stress fibers. It is unclear if this reflects a dynamic process or something about the maturation state of these junctions, or if the epitope recognized by anti-EmFAK is simply less accessible at these structures. [a = antibody only; a' = F-actin (red), DNA (blue) EmFAK (green)].

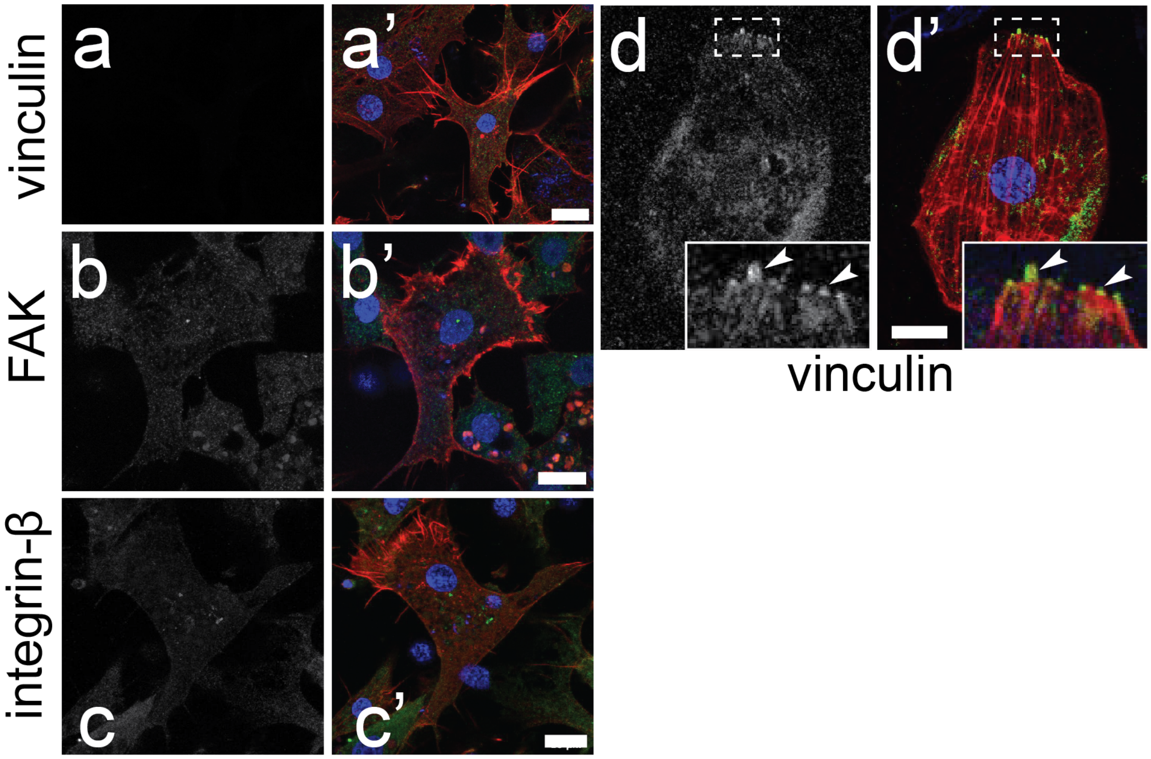


**Supplemental Figure 4: Migratory cells form focal adhesion like structures in two-dimensional, but not three-dimensional environments**
Migratory cells in the mesohyl where stained for (a) EmVcl, (b) EmFAK and (c) EmITGB. Focal adhesion-like structures were not detected in this three-dimensional environment. (d) Basopinacocytes sometimes detach from the leading edge of the spreading juvenile sponge and migrate on the coverslip. In this context, EmVcl was detected at focal adhesion-like structures. Arrowheads indicate EmVcl positive focal adhesion-like structures. [(a-d) antibody staining only; (a’-d’) antibody = green, DNA = blue, F-actin = red; scale bar = 10µm].

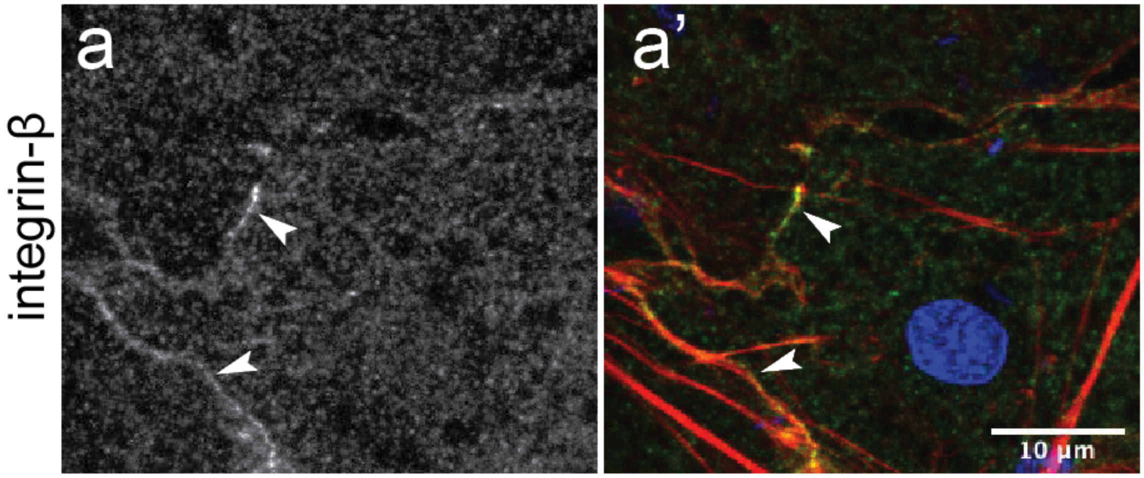


**Supplemental Figure 5: EmITGB detected at cell boundaries in the basopinacoderm**
EmITGB staining was patchy and of low intensity at cell boundaries in the basopinacoderm of early juveniles. [a = antibody only; a' = F-actin (red), DNA (blue), EmITGB (green].

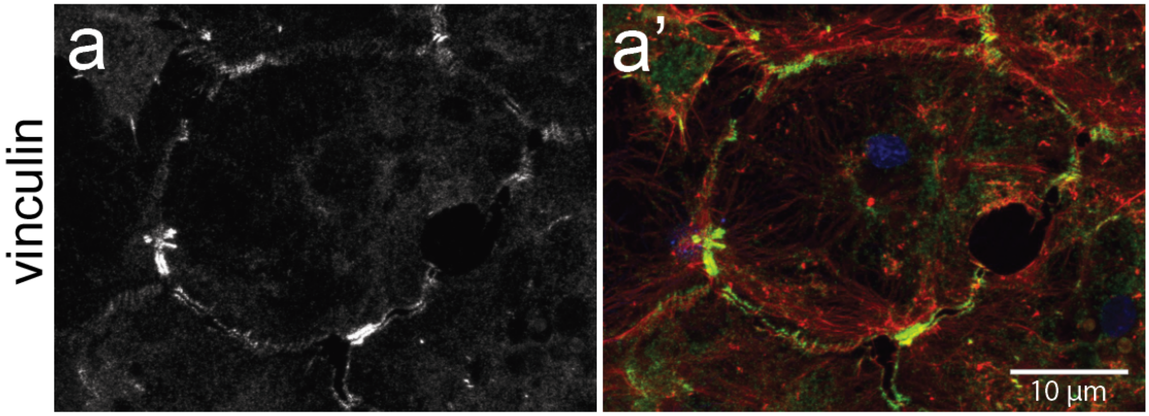


**Supplemental Figure 6: Robust cell-cell junctions in the basopinacoderm of 1 month old sponges.**
EmVcl was patchy and of low intensity in early juveniles, but robust in older (1 month) tissues, which formed cell-cell junctions similar to those in the apical endopinacoderm. [a = antibody only; a’ = F-actin (red), DNA (blue), EmVcl (green)].

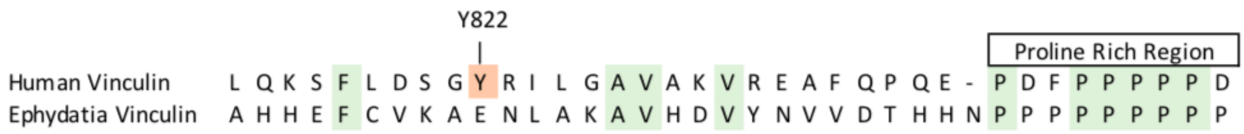


**Supplemental Figure 7: Alignment of Y822 region of *E. muelleri* and human vinculin**
Tyrosine 822 (Y822) of human vinculin is phosphorylated by Abelson kinase upon recruitment to the adherens junction. The homologous region of EmVcl (just before the proline rich region) contains a tyrosine residue but has low sequence conservation with human vinculin.

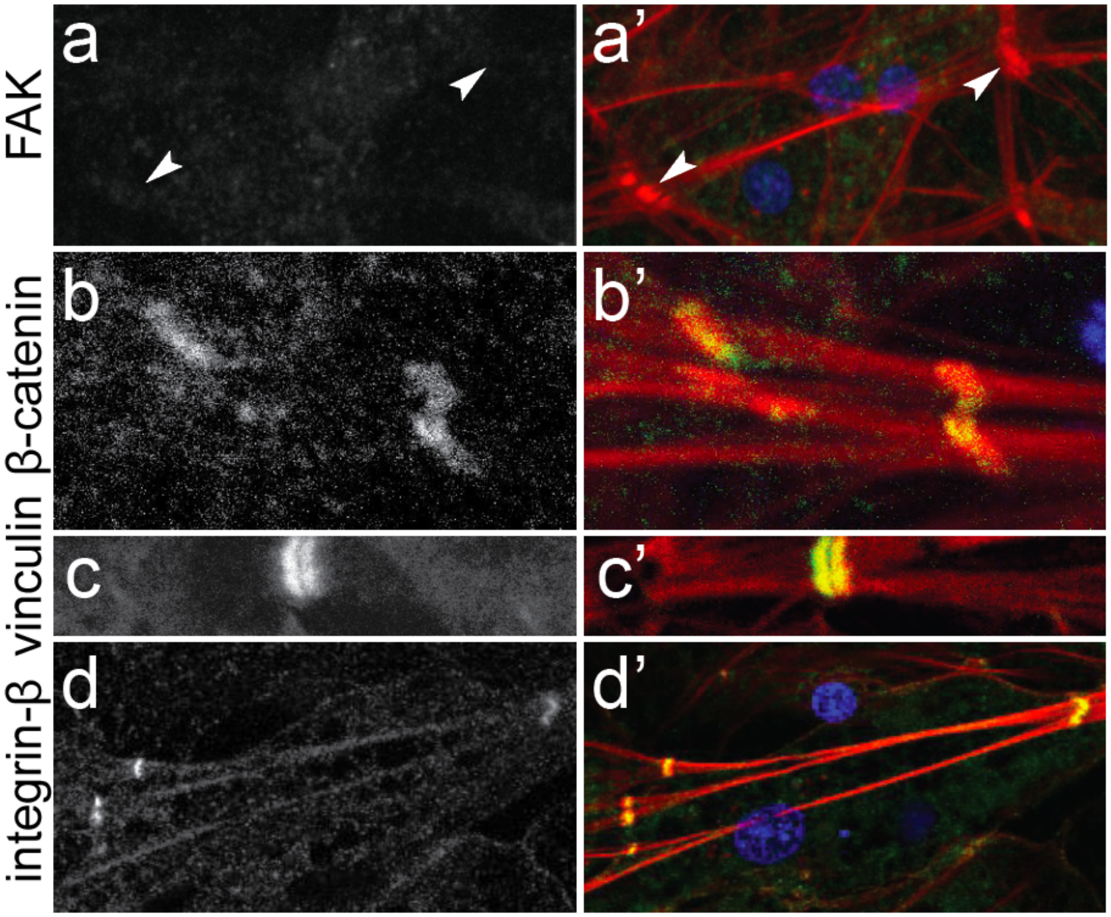


**Supplemental Figure 8: FAK inhibitor 14 effects on cell-cell junction composition**
(a) Treatment with 5 μM FAK inhibitor 14 (FAKi 14) abolished FAK staining at cell-cell junctions in the apical endopinacoderm, but had no effect on (b) Emβ-catenin, (c) EmVcl or (d) EmITG to staining. [a-d = antibody only; a’-d’ = F-actin (red), DNA (blue), antibody (green)].

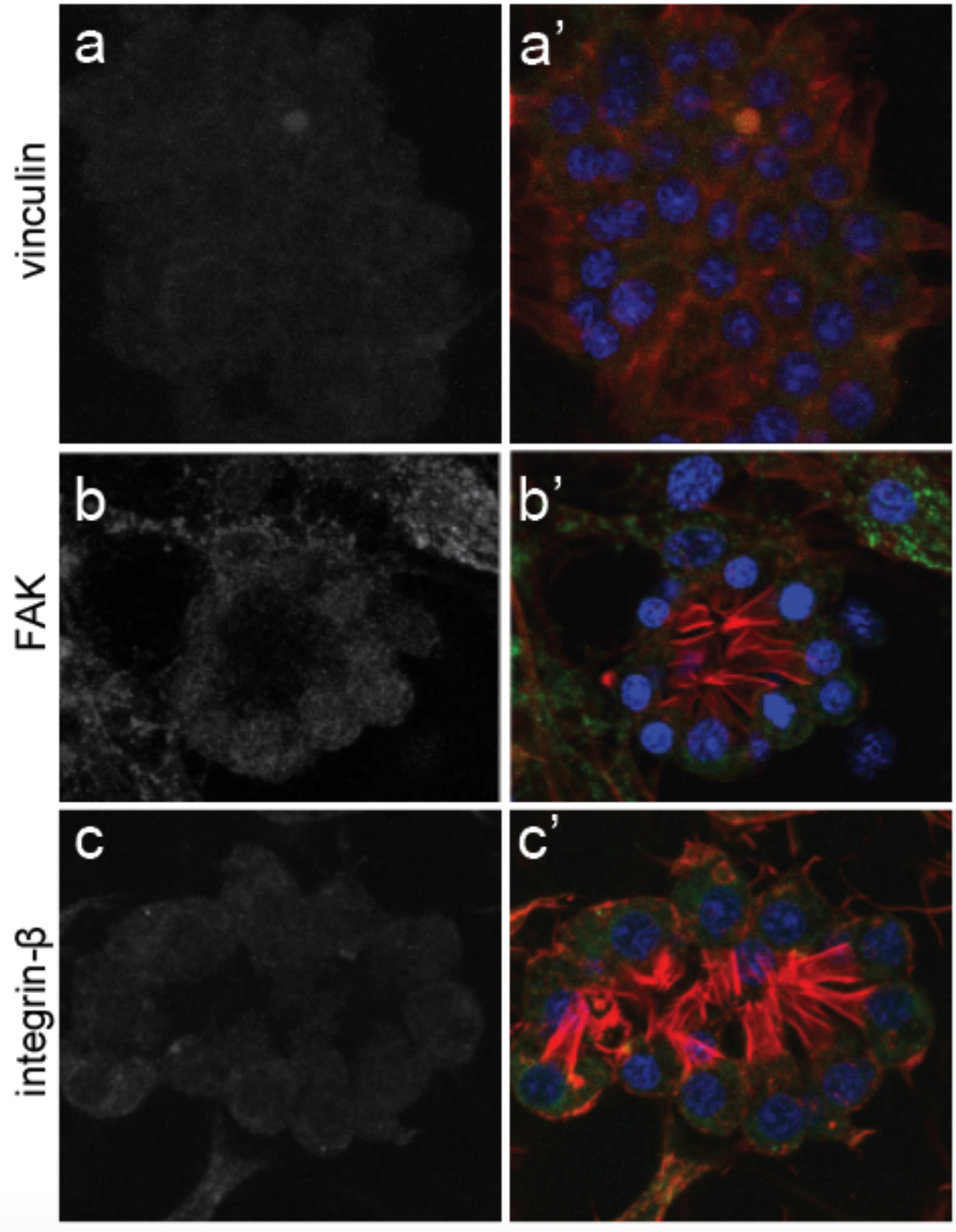


**Supplemental Figure 9: Focal adhesion proteins are not detected in the choanoderm**
No staining was detected for focal adhesion proteins in the choanoderm. [a-c = antibody only; a'-c' = F-actin (red), DNA (blue), antibody (green)].

**Supplemental Results: Antibody Validation (Western Blot and Immunoprecipitation)**

Anti-EmVcl recognized a specific band of the expected size by Western Blot of *E. muelleri* lysates used as input for immunoprecipitation. This band was depleted in the flow-through/unbound fraction and enriched in the precipitate elutions (Supplemental Figure 1a). LC-MS of the immunoprecipitate identified this protein as EmVcl, and there were few detected co-precipitates (Supplemental Figure 1a'; Supplemental File 3). Pre-adsorption of anti-EmVcl with 5 µg of the recombinant antigen completely abolished immunostaining signal (Supplemental Figure 2).

Anti-EmITGB1 had low affinity for multiple proteins in denatured *E. muelleri* lysates as detected by Western Blot, none of which matched the expected molecular weight of EmITGB (Supplemental Figure 1b). Immunoprecipitation with anti-EmITGB was hindered by cross-linking to the agarose bead resin, but was successful if the antibody was not cross-linked. Multiple bands were detected by Western Blot of the precipitate, one of which potentially corresponded to the expected size for EmITGB1 in precipitate, fraction 2. Analysis by LC-MS of the high-molecular weight fraction of EmITGB1 precipitates detected each of EmITGB1, EmITGB2 and EmITGB4. Of these, EmITGB1 was most abundant.

In contrast to the EmVcl IP, many other proteins were detected as co-precipitates of EmITGB1 (Supplemental File 4). The most abundant protein in the sample was EmITGA1 (it was also detected at very low levels in the IgG control). Integrin-beta is well known to heterodimerize with integrin-alpha, so it is probable that EmITGA1 is highly represented in the sample because it heterodimerizes with the multiple ITGB paralogs recognized by the antibody. Other than EmITGB, the most highly abundant protein in the precipitate was a phosphodiesterase (possibly PDE8). This protein was nearly equally abundant to EmITGB and has a predicted molecular weight of ~86 kDa. PDEs are known regulators of cell adhesion, and direct interactions with integrins have been characterized. Other known focal adhesion proteins detected in the sample include EmFAK, EmTalin2, EmITGA1, EmITGA2, and EmITGA3, strongly supporting conserved endogenous interactions between focal adhesion proteins in *E. muelleri*. Immunostaining signal was abolished upon preadsorption of anti-EmITGB with 1 µg of the injected antigen (Supplemental Figure 2).

Supplemental Figure 1c illustrates that anti-EmFAK recognizes multiples bands in denatured cell lysates by Western Blot, including a band of the expected size. This band was depleted in the flow through/unbound fraction, and enriched in precipitate. Like anti-EmITGB, anti-EmFAK activity was disrupted by cross-linking so immunoprecipitates were co-eluted with the uncrosslinked antibody. High molecular weight gel slices were used for LC-MS rather than the entire antibody-saturated precipitate. In the fraction sent for LC-MS, EmFAK was found to be highly enriched in the anti-EmFAK precipitate, and absent from the IgG negative control sample (Supplemental Figure 1c'; Supplemental File 5). Preadsorption of anti-EmFAK with 10 µg of the recombinant antigen fully abolished immunostaining signal (Supplemental Figure 2).

Collectively, these data strongly support that all three antibodies used in this study specifically bind to their expected targets under native conditions in *E. muelleri* tissues and lysates. Only EmVcl gives robust and specific signal under denaturing conditions (Western Blot). These data also support that EmITGB functions as part of a complex with EmITGA, EmFAK, EmTalin and possibly EmPDE8.
