## Supplemental File 1 for "Diverse cell junctions with unique molecular composition in tissues of a sponge (Porifera)"

>EmVcl_(comp69280_c0_seq1)
CTTTTTTTAAGTAAGACAGTTCAAAAAACTTATGAGATTTGCATAACTATGCCGGCCTTTCACACAAAGACGA
TTCATTATATCTTGGACCCTGTGGCTCAGCAGGTGTCCCAACTTGTGATCCTCCACGAAGATGCCCAGCAAGG
ACGCATCATGCCAGACATCTCGGCTCCTGTCACTGCGGTGTGTGCAGCTGTCCAGAACCTCATCGCTGTTGGC
CAGCAGACAGTGGGTCATAGTAAGGATGAGATCCTAAAGAAGGATCTGCCACTGACCTTGGACATGGTAGAGA
GTTCCTCCAAGATGCTGGTGGAGTCTGCCACGGGCTTGAAGGCTGACAGCCAGAGCAAGAAGCACCTGGAGCT
GCTGTTAAATGGAGCAAGAGGCATTCTCCAAGGGATCTCCAGCCTGCTCCTCACCTTCGATCAAGGAGAAGTG
AGAAAGATTGTGAAATCCTGCACTGGTGTGGCGGAGTACATCAAGGTGGCAGAGGTGGTACAGACCATGGACG
ATCTCGTCACCTTCACCAAGAACCTCTCTCCCGGCATCACCAGCATGACCAAGATGGTGGAGACACGTTACCA
GGACCTGACCAACCCTTCACACGCGAGCATACTTGCTGCCGAAAATGACCAGGTCAAGCAGGCCCTCCCTCTC
CTCCTCTCTTCAATGAAAGCTTTTGTCACCTTTCGTCGTGACAAGAAGAAAGGAGAAGCTGAAGCCCAGGAGA
ACCGGAACTACATAGTACAGGCCATGGGGGAATCCCTAGCTGAGATAATTCGTGTGCTCCAGCTGACCAGCCA
GGAGGAGGTCATGATGGCCTTGGCTGCGGAGACTGGATCCGCAAAGGGGACCATGGCTGGTGGGCTCCTAATG
GGGTCACTGGCTGCAAAGGTTCACTCGGCCAAGGAACTTGTCAGCCAAACAGGGACCAATGCCCAGACCAACA
AAGCTGGTATACAGGCTGTGGAGGCCGTGCTGGAGGAGGCAAGACGCATCGCAAAGACCTTGCCTGCCGACAG
GAAGGCTGTGATCGAGGGGCTCTGTGACGAGCTAGAGAGCCTCAAGAAGGAACTGGCCATGCTGCAGAGCAGT
GGGCAGGGGGACAGTGCCAGGGCTCATGCCATTGCCACTGCCCTCAACTCCAAGATGGATGAGCTGCAGATGA
AGCTGAGGGAAGCCATTACCAGGAAAGTGGCTGAAGACTTCATGGACCCTGTCGGACCCCTTAATGCCCTGAC
AGAGGCCAGCAGGGCTCCCCTCAATGCACCAGCACGTGCTGAGAACTATGTCAAGAGGGTCACTGACTTCCAG
GACCATTCCAAGAAGATGGCCGACACTGCTGTAGCTCTGGCCAAATCCGGCATTGTGACAGACAAGAATCTTG
CTGACAGTCTCCTGTTAACTGCTGGCAAGTTAAAAGCAGTGGCGCCACAAGTTGTCTATGCTGCAAAGATCGT
CTACGAGAACCCTGACAGCAAGGAAGCCAAAGAACATTACGACATGCTGAAGGAAGACTACCAGAAGCAAGTC
CAGAAGCTTACCAAACTAGTGGACAGCGGTCTGGACACCGTTGAGTTCCTAAAGGCCTCGGAGAGCATGTTGA
GGGACGAGCTGGAGGCTGCAAGGACTATCACCAAGGCTGGAACTGACCCTCAGTCTGCCTTCAAGCACATAGC
CACAGCGGCCAGGACGGCCAATCGAGTGGTTGCAGTCGTGCAAGGAGAATCAGAAAACAGCGAGGATCCTGCA
TTCAAAACCGAACTGGCTTCAAGTTCCCAGGCCATAACTGCTGCCATCGGTCCCATGGTAACCAGTGCAAAGA
CCTGCATTCAACAGGGGGGAAGTGCCACAGCTCACCATGAGTTCTGTGTGAAGGCTGAGAATCTGGCAAAAGC
AGTCCACGATGTGTACAATGTGGTGGACACACATCACAACCCTCCACCTCCACCTCCACCTCCACCAGTCGAA
GAAGCCCCAGAACGGCCACCTCTGCCGGCCGAGCCGGAAGTACCTCCACGCCCACCATCTCCTGAACTTGTGG
AGGTGGTGCCACTGCAATCCGTGGACCCCATTGGATATGCAGCACACAAGTTGGACAAAGATGCAAAGCAATG
GGAAGACAATGAAATGGTGACAACTGCAAGAAGGATGGCCAAGCTATTCATGCAGATGTCCAAGTTTGCCAGA
GGAGAGGAAGGTGAGGTGCACTCTAAAAAGGATTTCATCAACACTGCTCGCATGATCGCCAAGGAGAGTGAAG
ATGTCGTCAAGATGGCAAGGAAAGTGGCTGATGCATGCACTGATAAGAGGATGAAGAGAGCCATCCTTCAACT
TGTGGACAAGCTACCTACCATCTCCACCCAACTCAAGATCATAGCTGCGGTGAAAGCGACCAGGCAGGGCGGA
GATGATGCAGCCGCCGACCGAGAAGCCACGGAAATGCTAACTGATAACGCGCAGAACTTGATGCAGGCTGTCT
CCGAAGTCTTGTTCGCCACGGAAGCGGCCACCATTCGAGTCCCTCCGGATCAACGGGCCACACTGGGATTGCA
ATGGGTTAAGAAGGGAGCACGTACGCTCTAAAAAAAAACCAGTAAAAAGGCTATATTTAGCTGTGACGTTATT
CCATTTCAAACATTACGTAGTTCAGAATGCACAAATCGACTATAAATAAGAGCATGTTCTGGTGTTAACGATT
GAAAGAATGGCGGACGCTGTAAAAAAAAAA

Fwd primer red = green
Rvse primer region = red
Underline = CDS cloned for expression and antibody production

>EmFAK_(comp63096_c0_seq8)
TGCTTATTAATAACGTATTATTTGTGCTGTGGAGGGTCGTGTCTGTCTTTGTGGGCATTAGAACTGTGCAAGG
TTAACATGAATGGCGTCTGATGCAGTTATAAGAGTCTTCTTGGTTAATGGAGAGTCCAGGAGTGTCCGTGTGG
AAGAGAAGACGGATTCGATGGACGTCATTCGGTTTATCCTCCGTCGTCTTCACGTGAACACGGAGTACAGTGC
AAAGCTGTTCGGACTCCAACTGAAGCACACGTACAGTCATGAGTGCTACTGGCTGCAGCCAGGCTACACCATA
TTCGAGCTGCTGGATCAGTACTGCACCACAAAGCCTATGGAAGAGTGGAGGTTTTTTCTACGCATTCGAGTAC
TACCAAAGACAGCTCAGCACCTTTCGTCCCAAGATCCAGTTGCGTTTCAGTACTTCTATGACCAGGTGCTACA
TCTCTATTTGGAAGACGATACCGTACAGCTTGACAGTGAGACTGCTGTCAAACTAGGGTGCCTAGAGTTGAGA
CGCTTCTACAAAGACATGCCACAGATTGCTCTGAAGAAGAAAGAGAACTTCTCCATCCTTGAACGAGAGATTG

GTCTAGAAAAGTTCTTTAAAAGGTCGTTATTGGAGACCATTCCAAAGCGAAAGCTACGCAATCTTGTTGTCAG
TGCATTTGAGCAGTTGGAGACCTTGAACATGGATGGATGTATGTTCCAATTCTTCAATCTCCTTGCAAAACAA
TACCCATTGGATGTTGAGCAATTTTCCAACTGCTCTATTGGTGAGCGACAGGATGGACAGTCTAATGATCTGA
CATGTACCATCTTTGTTGGACCTAACTGTGATGTGGAGTATCAGTCAGAGGGCGGTACCAGGCGTTTCCTGGC
AGCCTTCAATGATATCCATGAAATATCATATGAGCCAGAGTTCGTGACAAGAGGGAAGGTGGTTCTGGAACTG
AGGCAGAATTCACAGTCCATAGTGATCCATACTGCTGTGCTGGTGAATGCTGTCCACCTGGCTACCCTGGTCA
ATGGCTACTGTATGGTGTACACCTCACTGCCCCACAGCAGGATGACAGGAGGGAGGAGAATCTCCAGTGCCAG
CAGACTGTCTGGTATGTACAACACGGTGGACGAGCTGAAGATGCGGGTGCCCAGCCCTTCAGGAGAACATCTC
GATGACTACGCAGAGGTGAAAGAACATCGCTCCTCTCTCTCTCAATTTGGTGCCACCATTCTGCCAAAGGACG
TGCAAGTAGGAGAGAGAATAGGCGAAGGGCAGTTTGGAGATGTGCACAAAGGTGTGCTGTTTCCTCAAACTAC
AGGAGAACAAGTAGTTGCTATCAAGACTTGCAAGCCTGAAGCAAGTGATGTTGAACGGGCCAAGTTCTTAGAA
GAAGCAGCCATCATGGCTAAATTCAACCATCGGCATATCATCAAGCTGTTTGGAGTGATGTCACACGACTCCA
CCACTTACATCATCATGGAGCTGGCTCTAATAGGACAGTTGCGTCGCTATCTGATGACGGAAGGGGCACACAT
CGCCTATCCAATACTGCTACAGTACATTTGTCAACTTTGCTCAGCTGTGGTCTATTTGGAGAGCAAAAACTTT
GTACATCGCGATATAGCAGCACGCAATGTGCTGTTGGCTACTCCAGAACTCATCAAGCTTGCTGACTTCGGCT
TATCAAAGCGTCTTGAGGATACTGACTACTATGTGGCATCTAAAGGCAAGCTGCCAATCAAATGGATGGCACC
AGAAAGCATAAACTTCAGGAAGTTCACTGGTCTAAGTGATGTGTGGATGTTTGGAGTGTGTTGCTGGGAGATC
CTGATGAAGGGAGTAAAACCATTTGTGGGTGTCAAGAACGACGAAGTCATCAACATGATTGAGATGGGACAAC
GCCTTCCCCTCCCCCCTGACTGCCCTGCCCCTCTCTTTGATCTGCTCAACCAGTGCTGGCAGTATGACGCTCA
AGACAGGCCAACTTTTGCAAAGTTGGAGCACATGCTGATGGCTATTGTTGAGCAAGAGAGATTGGAACAGCCA
CGCAAGACGAACAGTAGTGGCAGCTCAGCTAGACAGGACCCTTACGCTGTTATAAGGCAAGATGGACCCGAGA
AGCCTCCACGCAGGGACCAATCCAGCAGTAGTATTCACGGCAGAGGAGCTCAGGAGTCTTCAGTCAGATGGTC
TGGGTTTGTACCAGATGAACCCCCTCCTCCCCCTCCCCCCCGACTACGAGAAACTGATGATAGTCATCTTCGT
CTTAATGAAAGATTTCGTGGCACCTCAGGTAGTCCCAATGGAGACATTGGCTCTCCACCATTGGAACCTGCTC
CTTACCCGCCCAAACCACCGCGAACAGATAGTCCTGGTTCTCGTGATCGCATACGTCCTCCAGATCGTCCAAC
CCCAGCTCCCATCGTGCCTTACTCTGTCACCACTATAACACCAAGTGACCCCACCCCTCCCTCATTCCCTGTC
TCACCCACGTATCCTGTGCCTGAGCCTCGATTCAATGGCGCCATTCCTCCTCCTTCCATCCTGCGTCCCGCGG
CTTCCAGGTTGAGTGGAGACCACGAGAGGGTAGGGTACTATTCCGTGCCAGCAGAGGCAGGGGAGGGACCCAT
GGGGCGTGTATTGTCTGAAGATGGCACCTTTGTCGACAAGTATGGTGCGAACAAGCGTGGGGCGTCATCATCG
TCCAGCAGGCTCTCTACGAGCTCCTTGACATCCCAACCAGAGCCCAAGCAACCGGAACCAGCTGAAAGGGAGT
TGGATGAGTTTGATGATGAAAACGATGAGCTGTTGAAACAAACAACGGACGTGGTGAGGGCAGTGATGGAGAT
GAGCAACAAAGTGCCCATCTCCAGACCTGCCGATTATGTGGAGCTAGTCAAGAACGTTGGTAAAGCTCTGCGG
GAATTTCTGACCAAAGTGGAGATGGTTCAAAAGACCTTGCCCATAGAAAGCCACAATGAGATCATTATGGCAA
ACAAAGTGTTGTCTTCAGATGTCACCAGACTAGTGGATGCCATGAGAGATGCTCAGAAAAACTATCAGACTTT
CCTTGAACAAGAGTACCAGAAGCAAATGTTAAAGGCGGGACATATAATTGCAGTCAATGCAAAGCAGCTGCTG
GACACAGTGAACAGTGCAAGGCGCAAAGTATTGAGGACACCCAGATAGTCCAGATTTTACCCTTACTGCATTT
TGTGTTGCTGGTTTTTAAAATTATTGTCAGTGGTGTTTCAAGACAATAATAGTATGGTTTCATGTGTCGTTAC
AGTATAAGTGCTCTTTCCATGTACTGTTGTTGCAACTATAAAGCACACACACATGTCAAATCTATATCATTGA
TGGTCTCTATTAGCATAAACATTTCTGAAG

Fwd primer red = green
Rvse primer region = red
Underline = CDS cloned for expression and antibody production

>EmITGB1_(comp68476_c0_seq4)
CCCGCGCTTCCTTTTTTGGTCAAATTGATCGTTAAATTATTATTACCAATTCAGCTAGCTATCTGGATTGGGG
AGATGTTTATGACACTTTCTCTTGCAGTACTTGTTGCATTGTGCTACAGCGAAGCATTTGCGCAACCATGTAC
TGATCAGACGATGTGCGGAAAGTGCCTCCAAACAGCAGGCTGCGTTTGGTGCAACCTCACAACCTTCGATGGC
GCACGATGCTTTGGAAGGAACGTTAGCTCATCAATGGGTTGCAGCAGCATCGTTGATCCCAGGAGTGCTCCTA
CCACCACAACTGTCCCAGCTGCAACGGACTCAATATATACTACCACTCCAAACGTTGTACTGAGACCAGGTGA
TCCTTTGACTGTCAGTGTGAATGTTGTGTCACTACCCAATGCACCGTTGGACTTGTACATCTTGATGGACCTG
TCTGACTCCATGGCTGCTCCATTGGCAACTGTGAAAAGTATTTCACAGCTTATCGCTCAACAAGTGTCCAGTA
TCACCACCAATGTCAGGATTGGATTTGGAGCCTTCAATGACAAGCCAATCTACCCCTACTCACCTCAGACACC

AGCTGGCTGTCTTCCTGGTAGGGATGCTCCGGACTGCTCTGATAGGAGAGCAGGAACTCGTCAGTACAGTTTT
TTGCATCTAGCCAACTTCACTTCGAATTTTACTGTACCTAATGTGTTTGTCACCACCAACCTAGACCTCCCTG
AGTCTTCATTTGATTCATTGGTTCAGGTCCTTGCATGTGAAAAAGAGCTTGGATGGAGGAACCGCAGTGTTGA
GGGCCCAGAACGTGGGTTGCAAAGGCTTGTGTTGCTCATAACAGATAACCAGCCTCACCTTGCGGGAGATGGA
CGCCTGGCAAGCATATATCAACCCAATGATGGGAAATGTCACGTAAGGCCATATGCCTCTTCGGTGGGGTATT
TAGACACTGCTCCTGGTATACTGATATACAATGAAGATTCACTGCTCTATGACTACCCCAGTGTGGGTCTTGT
GGCTAGCCTTCTGAAGAAGTACGATGTCATCCCTATATTTGGCATTGTCCCAATTACATCAAGTACATTATTG
ATCAATAATACCTTCTTATCGTCTTATCAGGCTCTCCAGGACCTAATGACCAGTGTTGGAACAAAGGCCTTTG
CACGACCAATCTCGTCCTCTGCCTCTGATGTACTGGATGTGATCAAAACTGTGTATCAGGAGGTGATCCAGAA
CATTGCAATAACACTCCCACCTCAAAGTGATGTTGCAGTGTCTCTTTCTCAAATAACCTGTCCCGATGGCTCA
ATTCTTGTTGGTCAAACATGCACAAACGTGCCATTGTCCCGTACAACAACATTTAGTGTGACGCTTACACTTT
TGAACTGCAACACACCATCATCAAGCGCGCTCACATTCTCAGTGCCAGGATTTGGAACAACCACTATATCTGT
GGACAAGGTCTGTAGCTGCTCATGTGACAAGAATGTGACAGTGAACGCACAGCAATGTAACTTCCGAGGAAAT
TTCTCTTGTGGTGGATGCATGTGCATAGCAGGGTGGACTGGGCCAGCGTGTGAAAGGTCAAAATGCGGTCAAC
CTTGTGTGAATAATGGCACATGTGATAGTGCCACTGGCATGTGCCAGTGTACAGATTACACAGCAGGACCATT
CAGTAATGACAGCACAAGTGGAATCCTGCTGCCTCCATATACACCCAAATTCGCTGGGTCAACCTGTTCGTGC
AATAATTTCCAGAATTGTCCAACTAACAGCCAGAATTACATATGCAGTGGCAGGGGACAGTGTGCCTGTGGTA
GTTGTGCATGTGATGCAACACCGTACAGCTGGAAGTGGCAGGGCAAAGCCTGCGAATGTCCCGCTTCCAATTA
CAGTGACTGCTTTGACACAACATTTAAGAGTGGTCCACTTTGCAGTGGTAATGGTATGTGCTCTTGTGACAGT
TCTGGCAAAGGAATGTGTGTGTGCAGCAGTGGCTACACTGGAAAGTACTGTGAGACTAAAATAACGCCAACTT
GTGACACGATTGCTACATGCATAGCGGATGGCACTTGTGGCAGTTTAATGGCAGCAGACTCAGGTCAGCAAGT
TGTCTTGACCTGCCCCATTTCATCTGGAGAGTGTACATATAGCTACGATTTATCTCCTGATAACCAAGTGATC
AGAATGAACCAAGTGTGTTCATTTGCGGCATGGAAAATCATTGTGATAGTGATCTGTGGACTTCTGTTCCTGT
TTGTAGTCATATGTGCCATCATCAAAATTATTCTGGTGATTTTGGACTATGTAGAGGTCAGACGATGGGAGAA
AGAACTGAAGGAGGCTGACTTCTCAAAGAATCAAAACCCCCTCTACCAGAGTCCTGAAATGCAGTATACGAAT
GTGGCCTATGGAAAAGCCATGTGACAAGAGTTGTCTGAGTTTTGCAGTTGAACTTTGATTAGTGTACTAGGGA
GTATTTTGTGGTATGCATGTCGTTTTACTAACCCCCACAGCACTACGGTAATTACTAATTAGACGCAAGAGTG
TGAATTTGTATGTCAATTACAGTACAAGATTGTAACCAAAAAAAAAAA

Fwd primer red = green
Rvse primer region = red
Underline = CDS cloned for expression and antibody production

>EmITGB2_(comp69450_c0_seq1)
GTCACTTCCGATTGTCGGTTCATCGCCGGAGCGTTGATAAAGCAAGCGATCGTAAACACCGTTAACAGCACAG
TTCAGTGCAAATTTTTTGGCGTGCCTAGGCTGAGCCGCGAGAGAATGAAGCTCGGAGGGTACTGTTGCGTTTT
GTTATCGCTCGCTTCTATGAGCGGATCGGGCAACGCTCAACTCTGCACCGCTCAGACGTCATGTGCAGACTGT
ATAAACCTTTCCCCGTCATGCAAATGGTGCTCGACAGCAAACTACACCGGTTCCAGGTGCTTCTCGGGAACGC
CCGCAGTAAACTGTAGCAACGTTGAGAACCCCGCGGGGACTGTGACTGGAATAGACACCGCTACGTTAAGCTC
TCTTGTGCAAGTCTCTGCTCGTCAGGTCAACGTTACTGTAAGACCAGGCATTGGCACAAGCTTCAGTCTCAGT
GTGCAGCCGGCCAACAACTACCCTCTCGATGTCTACTTGCTGACTGACCTGTCCTACTCATTCCTTGATGACC
TCACAACCCTGCAAGCCCTGGGAGCCAGAATCGCAAGCTCTGTTCAAAACATATCTACCAACGCTCAAGTGGG
GTTTGGATCATTCGTGGACAAAAAGCTGGCTCCATTCATCAACATCCTGCCTGCACTGGTCAATGACCCATGT
GCGCCCCCTTATGGACCAGGTAAGTGTAACCCTCCATACAGCTACAAACACACTGTTAGTCTCACCACTGATG
GGGCATATTTCAGCTCACGCCTGCAACAACAGACGATTTCAGGCAATCAGGATGTTCCCGAGGGAGGTTGGGA
TGGCTTAATGCAGGCCATTGTTTGCAAGAAGCTGATTGGATGGAGGGACAATGCCAGACACCTGCTTGTCTTC
AGCACTGACGCCAATTCTCACCACGCAGGAGACGGGCTGTTAGGAGGCGTCGTCAGACCCAACCCCCATACGT
GCCTTATGAACAATTCTATCACGGCAGGCAATGTGGAGTACACCCTGAGCGAAACATATGACTACCCTTCCCT
TGGAGACATCAGAGAGCAACTTCGTCTCAACGACATCATTCCGATCTTTGCTGTTACGCCTGATGTGCAGAGT
ATCTACAATGCTGTGACAGCAGAGTTGGCCTCTGTTGGGGCTTCCACTGGCTCCCTGCAGAGTGACTCCGGGA
ACATCATTCAACTAATTCAGACGGCATATCAGACTGTCTCTCAGAGGATAGTGTTTGACCCTGTGCTGCCCAG
TGGCGTCACGATGACCTTCACTCCGCTAAATTGCCCGTTATTGGGGAGTGACAATGTGTGCAATGGTGTAAAG
CCCACCCAGGGAGCAGTGAACTTCACAGTCAGTGTCCAGTTGACGCAGGACTTCTGCAGAGCCAATCAGGGGA

ACACCATCTCTGTTCCCGTTCGAATCATCGGATTTGGGAGCTTCGTTGTCAACATCAGTCCTCTGTGTGGATG
TCCTTGTCAGCAGAGCCAGATTTCAAACAGTCCCTTTTGCACCTCAAATGGGACCCTCACATGCGGATTATGT
ACTTGCAATCCGGGAAGGTTTGGGAGCTCCTGTCAATGTGATGCAAATGGCGCCCAATCAGGGAATGCCACGT
CTTGCCCGACTGGACCCAATAACCTGCCGTGCTCTGGGCAAAGCAGGGGTAGCTGCATATGTGGAAAGTGTGC
CTGCAGTCAGTACCAGGACATTCGTCTTGGTCTCACCTCTACCTACTACGGCTCGGCTTGCGAGTGTGACAAT
TCGTTGTGCGACACGTCCAATGGACAGCTCTGTGGGGGCAGCTCCCAGGGGGTCTGTCAGTGTGGGGGTTGCC
AGTGTGCCAATGGTTTCTATGGCACGGCCTGCCAGTGCTCAAACTCTCTGTGCGTGGACCCCACTGACACAAT
CACTACACGGACTTGCAATGGGAGGGGAGTGTGTAGCTGCAATCAGTGCACAGCCTGCAAGCCCCCTTACACT
GGCCAGTACTGCCAGAGCTGCCAGGCCACGGACAAGACCACGTGCGCCAGTCAACTGTGCCCACCAAACCTTG
ACTGCGCTAAGTGTGCCCTCCTCAACCAAACCATGTGTCCCTCGTGCCCGACCACCTACTTCGTCAATGCCAC
AACCCTAAGTACCATTTCAGGTGCTGCCACCACAGAGTGTCAGTACACAGACTCTGATGGTTGCCAGGATACC
TACTTTGTTGTGCAGGATACAAAGGGCAATGTCACAGCCCTATATGTGAGAACAGACAAAGCATGTCCTCAGC
CATTAGCTCCCGCCCAACTCGCCACCATAATCGTAGTGCCTCTGGTCGTTATCGCCATAATTGCAATCCTTAT
TCTTCTGACACTGCTGCTTATCTTCTGGCTATTGAACCGTGCAGAAGTGCGTAAGTTTGAAAAGGAGTTGGCC
CGAGCAAAGTATACTAAGAACCAAAATCCCCTGTACGTGCCTGCAAACCAACAGACGAAAAATCCCATCTACG
AGGGCGAAAAGGCTCAGTAGCTTCCTGCCGGAGCAGTTCAGTTGAATATTACCCAGTCTTCATATAGTACAGC
ACAAAATGTGTGAACACTTGTTAAATTTTTTGTGTATATTGTAGTGTTGTATCATTCACAATGAGATTAAAAT
TTAGCAGCTAAAAAAAAAAAA

>EmITGB3_(comp69866_c0_seq1)
AATAGGAAGAAATTTTTGTGGGACAGGATCGAGGAGTCAGATCGTTTGGACTTGGCGAGCAACCTGAACTTGA
ACTACTAGGCGTGTGTCACGCCGCGTGTTGAGAATCGCATTCAGTCTCCAGGCACATGCACATGCAGCATCAC
AGAAGAAATGCGTGTTGGATCGTCATCCTGATCTGGCTCCCATTTGCTTATGGACAAGGAGCATGTTCAGCTT
ACACACGCTGTTCAGACTGCCTCCTGGAAAATCCATCTTGTGGTTGGTGTAATGACCCGAGTGTATTCCAGAG
AGAAGTGGGTGTGGCCTACCAGCTCTCCAGCCTGAACCCAATCTCACTGTGCAGGAACCTCACTGAGCTCTCC
AACAGTCTCAACTGTCCCGCCCAAAGCATTCTCTTCCCAAAGAGCTCAAATGTGACCACTTATCAGCCATCCT
CACCAGTGCAACCATCAAGTGTAGTAGTCTCACTGAGACCAGGTGATTCCTTTCAAATACCCCTCACCGTAAC
TCCTCCCCAGTCCTTACCCATCGACCTGTACATACTGATGGACCTTACGAAGAGTCTGGAACCCTATGTGAAT
GGTCTGAAGACCACTGCCACCAAACTAATTACCACAATGCAAGGTTTGACAAGCAAGTTTCGCATTGCGTTTG
GATCCTATGTTGACAAACGATTGGCACCTTTTAGTGATAGGGAAAGTCTGGATAACCCCTGTGAAGGGGTGAC
TGCAGCTGGAGTCTGCAACGCTGTGTACGATTTCCACCACACCCTTAACTTCACTGACAATGCATCACTCTTC
ATGGAAACTCTGAATGCTTCTAATGTGTCAGCCAACCTGGACACTCCTGATGCCCTGTTGGATGCCCTCCTGC
AGATCGCTCTATGTGAGGACCAGGTAGGATGGTCACCAGCTGGCCAGTCCAGAAGGATCGTCTTCGTAATGAC
CACAGGAGACTACCACTACGCTCTGGATGGAACGCTTGCTGGCCTGGTCAACCCGCCATCACTCACCTGTCGA
CTGTCCCCATCAGGGGTCTACCAGGACAGTGAACTGTCCGACTACCCTTCTGCTGCCGTCATAAGCCAGGTGT
TGAACGAGAAGAGGATCATCCCCATATTCGCTCACATAGACACGTTTGCCACTTCCTACGTCGCATTGGCTAA
TCGTATCAAGAGCGCTTTCCTGGGGAAGCTCTTGGGAAACGAGAACAATTTGCCGCAGGTCTTGAGCAACACC
TACACAACACTTTCAAGCACCGTCATCCCCGTGGTTACGGGTACAAACAATGAGCGTTACCTTTCCATCAGCG
TTTCGCCGCAAAATAACTGTGCACCAGGATGGCTCCAGACTAGCACCACAAACACATGTGCGAACATCACAGT
GAACACAACGGTTCGCTACATAGCAACGTTGACGGTCGCCAAGGAGTTCTGCGCCCAACCCAGCAACAGCAGG
ACAGTGGCTGCCAACATACAGTTCATCGGGTTTGGTGACTTGCGGCTGAACATCTCCGTGATGTGCCAACCTT
GCCAGAGCTGTCTTTCAACGGACTACACAAGCAGCTCGTGCAGTAGCTCGGGTGCATTGGAGTGTTCCACCTG
TGTGTGCTCGCCTAACAAGAATGGACCGACATGCAACTGCAACACAGATCCACAAGCCATCAGTCTGTGCAGA
CCTGATACTACCACTGAGATGTGCAATGGACGAGGCAGTTGTGTGTGTGGCAAGTGTGTCTGTGATTCTGTGG
GCGGGGTACAATATGGCGGCCAGTTCTGCCAGTGTGATAGGAACAAGTGTCCCATGGGGTACAACAGCAAAGG
CCAGCTAGCCATCTGCTCGGGCAATGGAGACTGCTTCTGTGACAGCTGCTCTTGTAACCAAGGTTACACAGGA
TACTCCTGTGGTTGCCCCACCTCCCAGCTACAATGTGTCGAGCCGGGAGCGAAGAGTGTGTGTTCGAATGCAG
GTCAGTGTCTCTGTGGCATTTGCATATGCAGCAATGCCACAGCTCGCATAGGAACCTACTGCGAGGAGTGCAA
GACCTGCACTGGAGCTTGCAGCAACATACTGAGCTGTGTGGAGTGCCACATCACTGGAACGTGCGGTACCCGA
TGTGCCAACATCACCTACGTACAGAACCGTACCTCAGTGCCTGGCTATGATGGAACTATGGTGTTCCGTACCT
GCTCCATTACCTCCCAGAGCTGTGAGGTCATCTACGAGCTGGATCGCTATGTCACTGGATTGGAGGGAAATGT
TTATGTTGTCCTCGACACCACACAGAAGAACTATGCCACACTGCGTGGCAATGGTGACTGTAATCCTAGCCAG
GTGATTTGGCCCATCCCGGTGGGCATTGTGCTAGGGATCATTGTCGTTGGTGTCATTGCCCTCATCCTGTGGA
AAGCCTGCAGCCAGCTGGGTGAGTTCCTTGAATACAAGCAGTGGGAAAAGAGCCTTAGGGAAGAGACGAATAG

GAGTGGAAGCAACCCATTGTTTGTGGACCCTACATCCAGTTACTCTAATCCTCGCTACAACGCCAGATCCTAA
CAAAATCATTGTTCAGAAAATCATTTGTGTGTCAGGGAGGGATGCATTCATTTCCCATGTGTGTGGTGATTGA
TTGCATCACAGCTAAACAGTAGCAACTTCATTGTAGGCGTTATCAGCCATCACTAGCTCAATTTTTTTTTTTT
TTTGTATTCACACTGCAATTATAAATGATGACACAAAAAGATTAAACGCTGAGATGCAAGAAATTATCAGTTG
TGCAAATATTATGATGTAACACATGTCTGTTGTCAAGCAACTGCTGTATTGGATGGCTTCAGGACCCAAACAG
TCTATGAAAGGCATTGTGGAGAGAGCATACCTGTCAGTGTCGCTGATATCATATAAA

>EmITGB4_(comp35829_c0_seq1)
TATTAATCACGTGTGCTTCGAAAAGCTAAACAGGTGTAAATGGACCGGTGGAGACAGCATCGTACTGTTATCC
TTCAAACAGTATGGTTCGTTGGACTTTTGATGATGAAAGTTGGTTGGACTGCATCAATGTGCAGCACTGCACA
GAATTGTGCCACCTGTGTGTCCAGTGGAATCAACTGTGTGTGGTGTAGCCAGAGGACTGCCAACATCACCCCG
TCATGCATGGACAGATCAGTGGCTAACGTTTCGTGCAACTTGAGCTATGTCGAAGATCCTCAGGTGTTTGTAG
CAAACCTTGCAGAGGAAAATTTATCTGAGAGTGTGCTGATATCACCCCAGTCAGTTTATCTTAATCTGCGAAC
AGGACAACAGGCTGTCTTTAATGTGAGCGTGAAAACTAGCAGAACATACCCTTTGGACTTCTATTTCATGATG
GATCTGACAGGATCACTTAAATATGATGTTGAACAAGTGCAAGCTAGTGGAACAGACATTGCAAATGTACTGA
AGAACATCTCGCAGAATTATAAAGTTGGATTTGGCTCATTTGTGGCTAAACCTGTCCCTCCATTTGTTGTTGC
CATACCATACCGACAACCGGATGGCACTTGCTACAATAAAGAGGGGTCGTGTATTGAGCCTTATGCCTATCGC
CATATTCTCCAGATGACAAATGTTACAGAAACATTTCTGAACATTTTGAACACTAAGTTGAATTTGTCATCTG
CTGCGGAAAATCCACAAAGTGGAACAGATGCCTTAGCACAAGCCATTCTCTGCAAGAATATTGTTGGTTGGAG
AGATGAAGCCTTTCGGATGCTCATGCTCATTTCTGACAATGCTGTTCACTTTGCGGGTGATGGAAAAGCGGGT
GGAGTAGTGGTACCTTTTGATGGAAAATGTCACTTGGAGTGGAATAATGCCACAGGAACATATGATTACTTGC
CCAAGTATAGCGCACTATATGATTTCCCTTCTGTTGCGCAGTTGAAATCGCTAATTGCAGACACTGGTGTATC
TGTAATATTTGGAATTGCTGGTGCTAATACAAATAATACAACTCCGGGAACTTTTTTCTTCCCAGATGTGTAC
AAGGCTATAGCCAAGGTAATGGATATTCCGGAGAGTTCTGTGGCTATCCTTTCTAAAAACTCGACCAACATTC
TTCAAGTCATCAATGACCAATATCTGAAAGCCATAGGGCTTATTAAATTTTCAGTCCCTGCAGTGAATGGTGT
TAACATTACAGTTCAGCCCATAATGGGATGTAACAGTTCATTGCCTAGTGGCTGTAATGATGTCAGTCTGGAG
AAAGAAGTGGTCTTCCGTGTCACTGTTGGTCTTACTCACTGTCCATCAACTCCGCAAAACAATATCGTTGTTC
CACTTAGAATTCCGGCATTGGCACAACAAATTACCATTGAAATTAACCCCAATTGTCAATGTGCCTGTGAGTC
TAAACCGGCGGCAAATAGTTTATTGTGCAATAATCAGACTCTAGTGTGTGGTCTTTGTCACTGTGAGGCTGGC
AGATATGGAGAATTGTGCCAGTGTCGTGGAGCGACTGCTTGTCCTGTTGGATTGCAAGGACTGACCTGTTCAG
GGTCTGCAGGGATTTGTAAACCTGATAACTGTTATAAGTGTCAGTGTTTGGGCAGTTATTTTGGCGATGCTTG
TGAGTGTGACCGTTTGAAGTGCCTTACATCAAGTTCAGGTATCTGCTCAGGTGAAAGTAATGGTCTGTGTTCA
TGTTCTGGTAGCAACGTTGCATGTAGCTGTAAGCGAGCGCCTCTCTCAAATATCACTTACATGGGATCAGCCT
GCAATTGTGACCCTGATGATTGTGTCAACCGAGAAACCAATGCCACCTGCAGTGATCCCGCCATAGGAAGTAC
ACTTTGTCATTGTACCGGAAGCCCTTGTAGTTGTTCATGCAGTTGCCCAGCCAATACTGTTCCACCATTGTGT
GAGCCGCAAGCACTTGTGAATTCTCGATGCGTTGCTGCAAAGCAGTGTGCTGAGTGTGGTTCAACCAAAGCAT
TGTCTGAGTGTACAGAATGTGTTTTTATTAACAGTGACAGTCAAGCATACAAGTGTGGCGTGATTGTATCAGG
TACCTGTACTGACAGCCACACCTACTATGTGGACACTCAAAAGAGAGTGTATCTCAAGCGTAATACTGTTACT
TGTGACCCGGGACCGGGTCCAATAATCATCGTATTCAGTGTACTGGGAGCTATTATTGCACTTGGGCTAATAT
TTCTCATCATTGCAAAGATCATCCTGATATGCTTGGATCAGGTCGAGTACAAGAAATTTACATCACAATTGGA
GGGAGCAGACTGGGCACCGCGAAATAATCCATTATACATGTCACCAGAGCAGAACTATACGAATGTTTTATAC
AGAAAGCGCTCATATCGTGGAAGCAAGTGAAACTGCTTACTACCTCCTCGAGGCGCTTACACTCCACATATAC
TTTCTCACCTGTAGTCACATGTGACAATTATTTCTTTACTCATTTCTATTTGAGTGCATTTATATTTTTACCT
TGTTGCGTCAGTAGAATGGCATGAGGAACATTTTTGTGTAATTTATTTGCTTATCCATACAGTATTACAGTTG
TCTCATAGTTACAATTGAAAATAAGAGACATTCAGCGTTTAGGCAACATGCAA

>EmITGB5_(comp67946_c0_seq1)
GGCGAGTTTGCATGAATTTTGTTTGATGTGTTTGTACTACGGTGTGCACACACAAAGGGTCGTACATCGAGCG
ATGATAACTCGCGATCAGCTGCAGTGGGTTCTGTTAGTGCTGGCATGTCTATGCGACCTCGACAGTGCCCAGC
AGCTGTGCAGTTCGCAGACAAACTGCAGCCTATGCATTCAGGCATCGCCCTCTTGCCAATGGTGCTCCGATCC
GAGCTACGCGGGCTCGAGGTGTTTCTCCTTTGATACCCCAAATATCAACTGCAGCAAATCCTTCGTAGAGAAC
CCCATCGGAACGAAGACTGCGCAGACGATGGACGTACTGGGATCTACTGTTCAGATATCACCGGGCGTGGTCA
ACATCACCGTCAGACCTGGGTCTGTCACTAACTTCACCCTGAGCATACGGCCAGCTAGGAACTACCCTCTGGA
CCTGTACATCCTCACAGATCTCTCCTATTCCTTCAGTAACGATCTGAGCACACTGAAGACGCTTGGTACCAAC

ATCGCAAACACACTGTACAATATCACTACAGACTATCGCATAGGATTCGGGTCCTTTGTTGACAAAAAGGTGT
CCCCGTACGTTGACGTTACTCCATCAAGTTTGACGAATACAAATGCTCCATACAGTTTCAAGCACAGTGTCAG
CTTAACCAGTAACATTACTCTCTTTAACAACCGACTTGAAGCTCAAGTTATCTCTTATAATCAAGATGCACCA
GAGGGTGGCTTCGATGGTTTCATGCAGATCCTCCTATGTAAAAAGCTGATTGGTTGGCGTGACAATGCTCGCC
ATTTGCTCCTCTACGATACTGATGCAGACTCTCACCAAGCTGGTGATGGAAAGCTAGGTGGTGTTGTCAAGCC
CAACCCTCACACCTGCCTTATGAATGACACCTATGGTCTGCAGGACGTGGATTACCAAGCCTATGGTCTATAT
GACTACCCTTCGCTTGGCGACATCAGAGAGCAGCTGCGCATCAATAATGTCATTCCCATCTTAGCCGTTACAT
CTGACGAACTTGCACGTTACACGGCATACACTACAGAGCTACAGTCAGTAGGGGCTACAGTCGGGACATTGGC
CGCTGATTCAAGCAACATTTTAAACCTGATTTCTTCGGCCTACAAGAGTGTCACTCAGAAGATTGTATTTGAT
CCAGTTCTGCCAGCAGGTATTTCATCACTGAAGTTTATTCCAATCAACTGTCCACAACTCGAAAGTGACGGGA
TCACATGCTCCGGTGTCCAAATCGAGCAGACCGTCAATTTCACAGTGCAAGTGCAGCTGGCATCGTGCGCCAA
CAATCAGACCATGCAGATTCCCGTTCGGGTCGTTGGCTTCGGCACATTTACAATCAACGTTCAACCCATCTGC
AACTGCGGATGCGAAAGCTCTGGAACAGCTACAAATAGCACCAGCTGCACCAATGGTAATGGTATCCTCTCAT
GCGGTGTGTGCAAGTGCAACCCTGGAAGATTTGGAACTCTGTGCCAATGCAACAGCCAAGGCGTCGGCCAAGG
GTCCAGCACAAGCTGTCCGACTGGTCCAAACCAGATGCAGTGCTCAGGACAGAGTCGAGGGACGTGTGTGTGT
GGCAAGTGTGAGTGTGCCACTTTCCAGGACTCACGGTTTGGGACCACTTCCACTTACTTTGGCTCAGCATGTG
AGTGTGATAACTCTCGCTGTGACACGACCAATGGCCAGCTGTGCGGAGGGACCAACCAAGGGGTGTGTCAGTG
TGGAGGGTGCCAGTGTCTGGGATCGTACTATGGAAGTGCCTGCCAGTGTTCCAACTCTCTGTGTGTGGATCCC
AATGACAGCCAGGGGAGGGTGTGCAATGGCAGGGGGACCTGCTCCTGCAACCAGTGCTCGGGTTGCCAAGCGC
CCTTCACTGGAGTCTACTGCCAGAGCTGTCAGGCCTCCACCCCGGGAGCCTGCGCCAGCTTCCTCTGTCAGCC
CAACCTGGTGTGCGCCCAGTGTGCCCTGGGACAGGTCAACAATACTCTCTGCTCCTCCTGTCCCTCCCTATTC
CTCCTCAATACCACAGACATCAATAGCATCCAAGGCTCGATAACACAGTGCCAGTTTACAGACAGCAAAGGTT
GCACTTACACATACTACACAGTGGAAGATGCCAATGTAGTATTGCAGTCTGTATACGTTGATTCAACTCCAGT
GTGCCCAACGGTTTTGAGTCCAGCCAGTATTGCAGCCATTGTGATCACCCCTCTGGTAGTCATCGCTATCATT
GGTGTTGCTATCCTCTTAACTGTCATGCTGATCTTCTACCTTCTGAACAGAGCGGAACTGCGTAGGTTTGAGA
AGGAGGTGTCCAAGGCGTCCTTCGCTAAGAATTTAAATCCTCTGTACATTGCTGCCAGTACTGACATCATGAA
CCCAATATTTGATGGTGGAGAGGTCAAAGGCACAGCCATGTAGCGCTGTCTGAATGAGATGCTTTAATGCATA
GCCCTGCAATCTGTGTAGTAGTGGACTTGATATAGAAGTACATATGCATGCCTAGTGCTGTAAGCTGTTAATT
TGTCTGTGAATGTCAATTATGTTCTAAATAGTTATAATGATTA

>EmITGB6_(comp56683_c0_seq2)
GCTAAGCAGTGCCAGGAACTTATAGAATACACAGCTTACCCTATAGGGCCAATGGCGACATTTATCGTATCGT
CAGTGCTTCTACTATGCATCGCTTCAGCAACGGTCTGGGGACTGAGGCAAACTTCTTGCGACACCAGAACGAC
CTGCGGCGACTGCATTGCGTCATCTCCACAATGCGTGTGGTGCTCAGACCAGAATATTACGGGATCAAGATGC
TTTGCTCAAGGTTCGGGCCAAAACTGTTCCCAGAGTGGCCTGCAAAATCCAAAAGGTGGTATTTTGAATAAAT
CCCAGGAAACTCTGAGCCCAACAAATCAAATATCTCCACAACGGATAAGCGTCAGTGTCAGACCAGGAGAGGA
CATTGCTTTTGGCCCTCTGTCAGTGAAACCCGCCAGGAACTACCCCTTGGACATATACCTACTGATGGACCTC
TCATACTCCATGCTGGACAATCTGCAGAATCTGAAGATGTTAGGGGCTCAAATAGCGAGCAAGATCGTTGACG
TCACTACAAACTACAATTTGGGGTTCGGATCATTTATAGACAAGAAATTATCTCCTTACATCAATATCCTGCC
CAGCCTCCTTAAAGACCCCTGTGCACCACCATACGGAGTTGGAGGCTGTGTTCCCACATACAGTTTCAAACAC
GCCATCTCCCTAACCAGCAACAACACTGAATTCAATACAAAGATACAAGAACAAAACATCTCCGCCAATGTGG
ACCCTCCAGAGGGCGGCTTTGATGGTATACTGCAGGCAGCTGTTTGCCAACAGCTACTCAAGTGGAGACATCC
AGCCAGACACCTCCTTGTCTTTATCACAGATGGACCGTACCATCAAGCCGGAGATGGAAAACTTGGAGGTGTG
GTCACTCCCCATCCAGGAACATGTCTGATGGACCAGACTATCACAGATAGAGCAGTGGAATACGAAAAAGCAG
TCATATATGACTATCCATCACTTGGCCAGCTAAGAGACAAGTTGGAGCAGTATGACATACTGCCCATATTCGC
AGTGACTTCTGAAGTACAAAAACTGTACAGCGATGTAGCATCTCAGATGCAGGACATTGGAGCACAGGTCGCA
ACACTCGCCAGAGATTCCAGCAACGTCGTGGACCTGATCAGTGAAACTTACAGCAAAGTAGCACAGCAGATTC
AATTTCAGGCCGATCTGGTGCCTGGAGTCTCCATCAATGTGGTGCCTCGCAACTGTACTAAAATTCAGAATGG
CATATGCTCTGGCATTCAGATTGAACAGGAGGTGCAATTTGATGTCCACGTGTCATTGGATGGCGCTTGCACG
CCTGAATTGCAGAGCGGACCAAAAGAGGTTAATGTACGTGTCTTGGGGTTTGGCAGCTTCACCGTGGAGATAA
ACGCAATCTGCCGCTGCCCATGTGAGGATAAACCTGTGAAAAATAGCCAATTGTGCTCCAGTGGGAACGGTAC
ACAAGTGTGTGGCCTGTGCGTGTGCAATACAGGAAGATTTGGGGACCAGTGCCAATGTGATGGCAAGAGCGCT
AGCACAACGAACAGCTCCACTTGCGACTTGGGTTTCAACAACCTTCCATGCTCAGGAAACACGCGTGGTAACT
GCGTGTGTGGAAAGTGTCAATGCTCAGAGTTCAAGGACACGCGAGGGAACACTGGGAGATACTACGGCAACAA

ATGCGAGTGCGACGACCTCAGCTGCGAGTCGTCAAACGGTGCCCTGTGTGGGGGTGTTTCCCAGGGGGCGTGC
CAATGTGGGGTGTGCAAGTGCAAACCTGGATACAGTGGGTCGGCGTGCCAGTGTTCAGACCTCTTCTGCATTA
ACCCCTTGGACCCAAAGTCACAGATATGCAGCGGGCAAGGAAAATGCAGCTGCAATACCTGTGTCAACTGCAA
GGAGCCCTTTACAGGCCGCTACTGCCACTCGTGCATGTCCACACTCAACCAATGCACCAACTACTACTGCAAA
CCCAACCAACCGTGTGCCATGTGCGCTGTGGGCGTGGCCAAGGGACCCCAGTGTGATAAATGCACAAATTTCA
CCAGCGTGGAGACCTTCGATTCATACCCCATCGATTCCCTGGCCAACTGCCAGTTTACAGATGACAATGACTG
CATCTACAAGTTCTACATCAACACTACTCTCCGGGTTGTCGTGGCAACAACACCAAGTTGTAATGTACTTCCG
CCATGGTTCCTGGCTGCTGTCATCGCAGGCCCTTTGGTAGGACTGGCCATATTGGGATTGATTATACTAGCCA
TCGTTGCAATAATTATGCACATACTGAATGCAGTTGAGCTGAAGCGATTTGAGAAGGAGTTAAAGAGCGCAAA
ATCCACCAAGAATGACAATCCATTGTTTATTCGCGCAAACACTGAGTATGTGAACCCCATATATGGAAAATAG
TCCTATCTAGCGAGGATAAGCCATTTCAGCACGCGGCCACAACAAACTCTAGCCACTGTACTTCTGCACTGCA
ACTTCAAGCTGTCTTCTCAACTCTTCCTCCATCACTGTATTCTGAAATCAAGCATATTACAATTGTCACATTC
ACTGTATTTATATTTCAACCTTTCACTACAAAAACAAAAGCACACGCAAAAAAGTGTGACATTGAAAATTAAA
AAA

>EmITGB7_(comp61500_c0_seq1)
TTACAATGGATAAAAAGCGTTTAATGGTGATTAGCACCGTTATTCTGTTCGCGCGCTCTGTTTTATTGGTTAA
AGGGCTGAACTGTCAAACAGCTAAAAGTTGCAGCGAATGTGTGCAACGTGGAGTTGACTGCATTTGGTGCACT
CTGCCTAATGTTACATATCACTGTGTGCAGAGAAACACTGCTGAGGCTCATTCATGTGGAATGTATGTAGAAG
ACAAAGTCAGCAATTTCAGAAGTAAAGAAAAGAATGATCTTCTAAATAAAACGCTGATTTCTCCACAATCAGC
TTATATTCAAGTGCGCGTTGGAGATTCAGTTGCATTCAATGTGAGTGTGATGACAAGCAGAACCTACCCAGTT
GATTTCTATATTTTAATGGATTTGTCTTCATCACTGAGAGATGATGTCGAAACATTAAAAAACACTACCCATA
AGATAGTTGCCACACTTCTGAACATATCGAGCAATTATGCAGTGGGCTTTGGATCGTTTGTGGACAAACCTGT
CCCTCCATTTGTACCAAATATTCCCTACAGCACTCCCACCCCAAAAGGACCAGTATGCTTCAATCATCAAGCC
CAATGTGCAGAGCCCTACAGTTATCGACACATTTTAACTTTGACAAATGATTCCCAAAAATTGCAGTACATTG
TGAACACTCAGCTGAACATATCACATAGTTCTGATAACCCTGAGAGTGCCCTGGATGGTGCTGGTCAAGTTCT
AGCATGCAAAAAGCTAATAGGATGGAGAAATGAAGCGTTTCATATGATGATGATTATCACAGATGCAAACTAC
CATCGTGCAGGGGATGGCAAATTGGGTGGTGTAATTGTGCCATTTGATGGAAGGTGCCACATGGACAGAATTG
CAGCTGAACTTTACCAGTATAATCAGAGCACTTTTTATGATTATCCTTCAATTCCACAACTAAAAAATCTATT
TACAGATGTTGGTGTTACCCCAATATTTGCTGTGACCAAAACTGCTCAAAATTACTACAAGGACTTGGTGAAA
CAACTTGGGACTGGAGTTGTAGGGGAACTGGCCAACGATTCCAGCAATCTGGCCCAAATTATTGCAGAGAAAT
ATCTGAAAGCAATTGGCCACATTCAATTTTCAGTTCCTGAGATTGACGGTTTGACAGTAACAGTGCAGGCTGT
ATCTGGATGTGAAACGAAATTACCATCAGGTTGTGCAGATGTAAAGCTGGAGCAACAGGTCACCTTCAGAGTG
ACAGTGGCACTGGACAAGTGCACTGTCAATATCATGAGGCAGCTTCAATCTCAGCAAAGCTTTGACCTCATCC
TCAAAATTCCTGTTCTTGCTCAAGAGTTCATTATTCACCTGACACCAATTTGCAAGTGCAGTTGTGAGATGGA
GAACGAAATGAACAGTGCAACATGTCATGCTGGTGGAAGTCTTGCATGTGGTCTCTGCTCTTGTAACATGGAA
CAGAAGAGATTTGGAACATTTTGTGAATGTCAAGGAAATCAAGCATGCCCCATCGGTCTAGCAAATTTGAATT
GCTCTGGAGCTGATCATGGCACTTGTTTAGGCAACTGCTTCGAGTGCCAGTGTAAGGAAAAGTATTTTGGACC
AGCATGTCAATGCAACGCTGAATATTGTCCAGCAGTCAAAGGACAAGTGTGTTCAAATCATGGTACCTGTCAA
TGTCCTCATACTGTTTGTGAATGCAATACAGCACCATTATCTCAGCTGAAGTACTCTGGCACAGCATGCAGTT
GTGACCCTGATTTTTGTGTCAATCCAAAGACTAATGCAGTGTGCAGCAGATCAAATATAACTGAAGGAAAAAC
TCTATGTCATTGCTCTGAAAATAAAGCGCAATGTAAATGTGGTTGCACGTGCCCTGCAGGCACTGCTCTACCA
TTTTGTCAGGATGAAACTGAAGTGAGTGATGTATGCATGAAACAGGAAAAATGTGCCCTCTGCGTGCTGCGCA
TGGGAGGAGGATCAAAATGTGACGGGTGCAAAGCAGAAATATTGTCCGACATAGCAACAGTACAACTTCAAAA
ATCGAGGTGTCCACCCTTACTGTTTGATGGCTGTTATTATGACTACTTTGTTGATCCATTTGGAACAATTTCA
GTGTCAATGGTTCCAGACAGTTGCCCTCCTGCTTCAGTGAACCCATGGTATATCGCCATTGGTGTTTTGGGTG
GTGTGATCATCACTGGAATCATAAGTCTGGTTCTTATCAAGCTGATATTGATGATTATGGATCGTGTCGAATA
TAAAAAGTTTGCAAAACACTTGGCTGAAGCTAACTGGGCACAGAATGACAATCCATTGTATGTGTCCCCAACT
CGACATTATGACAACGTAGCATATGAACGAAACAGGTCAAGACCTGGTAATTAAGTGAATGCAACAACCTATT
GAATTACTTGAGCTGTATAATATTTTAGTGAAATGCACATCAATGTTGACATATTTGATCATTATTTACAG

>EmITGA1_(comp66904_c0_seq1)
CGAGGACGGATCACGTGCGGCTTATAGGGTACTTGGTGTTTCCGTTGAGTTGTGTCATTGTCGCTGCCGCACT
GGTAACTCCACACAGCGATGATCCAGCTTTTAGCGGGTGCCTTCCTTATTTCTGTTTATGGGGTCATCGCTCA

ACGCCTTCCGTTGGACACGACAGTTCCGATAACACGAATCGCTCCGCCCTTGAGGAGAGGCGATGCTAATCCC
GATAACTTCGGTTTTGCAGTAGCCCTCTATCAATTGGACCCAGCGGGCACGAACCTTTCGTTGTGGAGAGTGG
TGGTTGGAGCACCCAACGGGTCCTACCCCGGTGGGCTCTCACTAACTAATCCCAGTTGTTCAACTCCCACAAT
TAATAACACTGGACTGGTCTACCTGTGTTCGATACAGCCGGGAAAGGACACTTGTGATGGCGCGGTGGGCAAC
GGGACTGTCGATAACGGCAGACTCTTCGATTGTCAAGCACCTAGTAATTCTCCGAACTATCAGCAGACGGGGG
CATCTTTGTACAGTTCTGGCGGTTACCTCATCGCCTGTGCACCTGGATTTAGTACCGGATCTTACAGCCAACC
TGTGCACAGGGGAACATGCTACGTGTCCTTCAACAACTCTCTTCGTACATTTGCCCAAATCTTCCCTTGCCGG
TCAGCACCGGTGACCACGTTTAGCTATTATGACGAAACCTACTGCTATGCAGGACTGAGCGTGGCTTTGAAGA
ACAAGATAGCAATGTTCGGAAATCCTCTAGCGTTCACAACTGGAATGGCATCTTATCTCCAGTTGGATACTCC
ACCTGTTAACTTCTCGATGATAGCACCAAAGAACAGGACCGTATCAACGGGTGGTTTGGTGTATGGCCCAACG
ACCATGAATATCAATGGCAGCATCGTATACAAAACCAATACGCTTAAAGGCTACAGTGTTGGAATCGGTCGCA
TCACTCGATCAAACGTATCTGACTACCTGGTTGCTACACCAAGGTGGAGCGTCGACAACTATGTGGGAACTGT
GGAAGTCTTTGCGTTTCCGAACACTGGAGCGCCACTTGCAGGCCCTTCTTATCCTGTTGGGAATGCCTTCTCT
ATTGGCGTGGGGTCGAATTCCTTCTTCAATACTGATGTCGGTGTGTACAATTCCTTGATGAATGTCGCAGCAT
ATGGTACACAGTCTGGTGAACAGTTTGGAGCTTCTATGGACACTGCTGACCTCGACCAAGACGGCTACGATGA
ACTGATTGTTGGAGCTCCATTTTATACTGACTACACAAAATCCAGCTACATGTATGAAGTTGGGAGGGTCTAC
GTTTTTCAGAACACAAAGGGAAACCTCAGTGCCACTCCCATCATCCTCTCGAGCCCCAATCCTTTCACCGGGG
GACGCTTCGGCCATGTTGTCTTGTCAGTCGGGGACATCAACGGTGATGGAACTGAAGACTTTGCTGTTGGGGC
TCCATATGAGTCCTGCACCTCAAGTGATGGTACCTCCTCCACTGGAAGTGTGTACCTGTATACTGGAGACAGA
ACTACTTTTGTGTCACAGACACCAATTCAAAAGATCACAGCTTGTGACATTCGTCAGAGGTCATTAACAGCTC
TGAATAACGCCACCCTTCGTAGCTTTGGCTACTCTTTGGCAAGCAAGGCAGACTTGGATGGAAATGCTTACAA
TGATTTGGCTGTTGGTGCGTTTGAATCAAGGGCAGTGTTTGTTTTGAGGACATTTTCAGTGGCCAATGTCACA
GCCACACTGACTAATGTGGGTGGTGGTGTGTCTGCGACAAAGGCAGGATGCAACAGTTCACATGCTTGTGGCA
TTGTGATGCTATGTGCCACGTACAATTGCAGTAGCAGGCCAGGCAGTTGTAGTCAAAGTCTGAGCATGACATT
TGACATTTCTGAGTCGACTATTAAGGCTTTCTTCAACACGACAACAATTGCACGCACGTCTACTGTTGTTGTC
ACTGCAAATATGTTAACCCCAAATTGTATCAACGTAACGTTCTATTCGATGCTTGGTCAAGTTGACCTCTCTC
CATTTAAGTTCACGGCCATAGTCAGAGATCTTTCTCGCGACCTTGTCTCTGATAGTGGCGCTGCCCTTGCTGA
TTTTAAGGCAATTCCAGTTCTGAACGGTGGCAGTGCTAGCATAAATGTTGGCTTTGCAAGGGCTTGTACGAAT
CCAAACGCTTGTGCGAGCCAGCTGGCACTTAGCCAGACTTCTGCTGTTGTGTTCAAGGATGCGCAAGCAACTG
TTGAGAAAGGTCTCATAGTGAATGAGACAGCAAGTATCACGTTCTCTCTCACTGTGAACAATTCTGGGGATGA
AGTGTTTGGAATCAATCTGGTTCTATCTGCCCCTTCCTTTGTGACTGGCATGATAATCACAAAGTCGTCTGGT
GTACCAATTACTTGCACGAAGACAAATACGACTGGCTTTGTCTGTCCTACTGTCGTTGATGGATTCCTGGCTC
AGGGCAATGCGAGTGTATTGGGAGTTGCCCTGACCATTGACACCACTTCCATCTCATTCGATGCATCAAACAT
GACGTTTAACGTCTCCTCTGACGACATCGAGACAAACACGCTGGACAACGTGGTCAAAATTCTGTTGCAGCCG
ACTGCGTCAGCAGACTTGAGCATCTTGTCCGCATCCTTCAGTCCTTCTGGTACCACTTACACTACCCCAAAAA
GCATCACCAACCCAACCAACTTGGGATCGATTGGGTTGTTGACCACTTTTGAAGTTACTTTCCAGAGGAGTGG
ACCTACTTATATCCCCAGTCTGACTCTGACCATCTCTCTACCTCTGGGGGGCTCCGATTGGAGTAGTTACTAC
CTGTATCCTGCGTCCGTGGCCAGCAGTGTATCCACCATAGCCGTCAGCTGCGCCGCGGGTGTGTTGAATCCTT
ACAACCTCCAAACCACTAAGAAGCGAAGTTTGAGTGCAGACATGATCAAAGATCTGCAGAGGGGTACCAGACA
AGCACAGAATGCATTAGTGCTCCTTGACTGCAGTCAATCATCAGCAGGTTGCAAGAACCTGGTCTGTAACATC
ACTAACATCACATCAACCCCCAGCTTCAGCGTCAGAGTTTCACTCTATGTGAATGACAAATACTTTTCGGCCA
GAGGTGATAACAGTAATTTCTCTGTAACAGCAGCAGCAACAGTCACTATTCCAACTCATGCAATTTTCACTGG
GATCGTTTCAAAGTCTTTCAAAACGGCAGTACTCAACATCAGTGCAGCACAAGCGCCACAGCCGAAGCCGCTG
AATCTTGTTGTAATCATCGTACCAATTGTTGCTGTTGTTGTGATCATTGTGATTGCAGTAATAGTTCTATATG
CATGTGGCTTTTTCAAGCGTAAGAAGAGGGAAGATGAGGACGCTGTGGAAGGAATTGATGGTGCTGTGGCCAC
TACAGCTGTCACCAAGAAGGACCCCTTGGAAGATTCTACAGTGAAGATGTAGACCTTGCTACACGTTCAAAAT
AATACTCACGTTCATTGCTAGGAAAATGTTGAACCATTAAAATATGCAGTCATATCTTTTACTTAGTTGTTAG
TGTATGTGTGTACATAGAGGGAGTGTGTAATTGTTTGTACGGTGTGGTATTTCCCTTCCCTAATCTTTTTGAA
ACACCATGCATTAAACGAGTTATCGTTAAAAAAAAAAA

>EmINTA2_(comp69364_c0_seq1)
GTCCTAACCCCGGAAGGATACAATAATAAAAGGAGTGAAACACAAAGCTTTAACAAGAAGAGTTGCAGACAAT
GATGTTAAGCGCAACGGGGTCTAGGCAAGCGACGACTACTTCGATCGCCCTACTGCTTCTGACAGCAGCAGTC
TGGAGCGTCCAGGCCCAGAACATCGATACAAAGCAGCCTTACATCAGATCTTCGCCCGATCAAACGAATGTGG

ACTACTTTGGATACAGCATCGTTCTTCACCAGACAATCGCCGGCAATCCTAGCTCTACAATGCTTATCGTCGG
AGCACCGAACGGCACAGCTCCGGGATCTCCCGTTAGATACACCGGTCTGATATACTCCTGCCCGTTGAATTCG
AGTATTACTTGCTCGGGACTAATGGGCAGCACGACCGGGACCGATAGAAGACTCTTCGACACAGACCCAAATT
CTGGCTCACCAACTCAGGTGGAAGAAAAGAGTGGACAGTTTTTGGGTTCTGTTCTTGTCAGCAAAGGTGACAA
GTTTATGGCTTGTGGGCACAGGTACTTCAACTGGGGGTCCAATGGGGGCTACCGCAGCTCTTTTGGACGTTGC
TTCATCGCTGGCAGAAGTCTAAGAAATTTTGCAGAATTTCAGCCCTGCGATGGAGTTGGAGTTAGACAGCCAT
ACTCTATTGATGTCTGTCAAGCTGGATTCAGTGGAGCCATAGCTAATGTCTCCAGATCAGGGCCTGGTGCTAT
TGCAGTAGGGGCTCCGGGAACATACACTTGGAGAGGCATTGTTATCAGAAATAACCCAACTACCAATGCCATC
CAGTTCACAGCCTACAATCCTTCTGCTACGATTCTCTATGGTTACTTTGGCTATTCAATGACCAGTGGTTACA
TCCTGTCTAAAACACAAGAGGATTATCTGGTCTCTTCACCTGATCTGAGCAATCTAGGAGCAGTGTCACTTGT
CAGAAACTTGGACACAGTTGATGTGGTCTCTGAGCCATTGCAGGGTCTGCAGATATCAGAATTATATGGTTTC
TCAGTTATCACGGCGGACCTTACGGGAGATGGGTACGATGAGGTTCTAGTTGGAGCCCCCCTTTATTCTCCTG
TGCAGAACCCAGAGGCTGGCAGGGTGTATGTCTACAGGAACATTGCAGGGACTCTACAATTTGTCACACAGCT
CATTGGTGATGGGATAAGCTATGGAAGGTTTGGCCATGCCATGGTCAACTTGGGCGACATTAATTCTGATGGC
TTTGCAGATGTGGCCATTAGCGCACCCTTCTCCAATGATGGAGGGAAGGTATACATCTACAATGGACAGAACA
TCAACACCATCAACACTGTGCCAGCTCAGATCATTGTAGGCAGATCGCTGCTGACAACGGCCAACCTTTCCTC
CCTCATTGGCTTTGGAGCATCGCTGGCAAGCCAAGTGGACATTGATGGCAACACCTACAATGACCTGGCCATC
GGTTCGTATCAAAGCCAGCAGGTCTTTGTTTTGAGAACGAGACCTATAGCACAGATGGCTGTGTCTCTCACTG
CAAGCAGCCTTCTGGTTCAGGTTTATAACGGTTACTTCCCGCTCTGCACTCTTAACAGTGTGAACTATACATG
TTTCAACGTCAGTGCCTGTGTTACCTACACAGGTGTTGGTGTTGCCAACCAACTCAATCTGAATGTCACTGTC
TTTGGCGACACAACCAACCAGCTGTTGCTTTTGACACCTCGAGTGTTCTTTGGAACGAACCAGCAGGCATCCA
GTGTAGTAACCACGGTGACAGCAACAAAGAATGTGCAAACCTGTTCCACGCTGAATGTGTACATCAAGAACAA
CATTGCAGACATCCTGAGCAGTTTTGTGTTCAACATGTCCGTATCAGTCCAGGACTTCAACCCGGCACCTTCA
AATGGGGCCACCTCCTCCCAAGACCTCTCCCTCTTCCCCATCCTGTCACAGAGCGGAGCCAACACTGTGCAGG
TGCAGACCAATAAAGGAGGCTGTGGTGCATGTATACCAGTTCCAGACTTGTCAGTCGAATACATCAACACAAC
TTACGATCAGAAGACCAACAGCTCGGACAACAGCTTCATTGCACAGGAGACCACAGGGATCAGCATATGGCTA
AGGGTGACCAATCGTAAGGACAACGCCTTTGCAACTGTTGTCTCTTTCAGCGTCCCCAAGACGCAACTCACAT
TTATCCGCTGTGGTCCTGACCTGAGTTTCCTGTCATCAGTCAAAGATATTTCTGCTACTGTATCTCTCTGCAC
GTGTCAGATTGCTAATCCGTTACTAAATGGAGAACACCGTGATGTTGTTATTCGTCTTGAACCAGCACCTACC
ATCGATGGATCTCAACTCAGCTATACGATCAACTTCAATGCTTCCAGCCAAAATGCTGAATACTCCAACACCA
CTTCTGACAACACGATATCCCTTCCGCTGGCAATCAAAACAGTCAGTGCTCTCTCAATAGACTCTGTGGGAAT
CGTCAAGCCAGAACAAATCATCTTCACCTCTTCTTCCGTCAACACCTCTGTTGCCCTGACGACTTCACTAGGA
CCTTACGTATTGACCACGTTCACAGTGCGCAACGGTGGTCCCTCGACCATTCCATTGGTGGGTCTGGACATCT
ACTGGCCTTTGGACAGCACCACAAAGGGCCTCTACTATCTCATTCCCACTTCAATTAAGGCTCTGTCGTCAGT
CTTCGTGACCCAGTGTGACTCAACCTACACCATGCTGCTACAGAACGATGTCAGCAATATCATTCCCACGGGC
AACAAGCGATCCACAGACGGTTTGAGAAGGACCAGGGCTGCCGTTGCAAGCGCACCCTTGCCTGGTGGCACCA
CTACGGTTGACTGCTTTTCCCAGCCACAGTCCTGTGTTCGAATAAGGTGCTCTATTTCACAGCTAAGCCAGGA
CCTGGTCACTATTGTAGTTAACTCTACTGTCGATTCTCGATTCTTTGCACAGGGACAAGGGCTAACAGCGCAA
TACAACTTCATTCCACGTGCCGCAGTTTCAATATCAGGGGCTGGAAGCAGTTATATTGTCGATATGGGCAGCA
ACAAGAATGCATCCGCTGATATTGTCATCATCTACCCTACTCAAGGGAGTAGCAGAGATCTTCCGTGGTGGGT
GTACGTGGTGATCATAGTCCCAGGTCTTCTGTTCCTGCTGATCACTACTGTCCTTGTGTTGGCGATCATCTAC
TATTGTAGCAAGCGAAGACAGGCCATCAAGAACAAGTACCAGGATAAGCAGGCATTGACAGGGATGCAGGGTC
CCACGGATGGCCAGTGAACACAGGGGGGGGGAGGAGTCAAACACTCACTATTCAAACATCTCAGCTTAGTATA
ATAGTCTATATCTTCTTTTCTGCAGTGGCTCTTAGAGGATAGAAAAGAATAATTATATGTTGTACATGTGTGT
TTTTTTTTTTGGG

>EmITGA3_(comp59085_c0_seq4)
CGGATTATTAATTCGGCAATAAAGAAATGAGGTGGATGCAAGTATGGAGTACGGTGTTATTTTTTGTTGGATG
GATAGTTTGCCTTCAAAGTGAGAGGCTGGACACTCTGGCCCCCATCATACGCAAGTCACCTGCAGCTTCGGTC
AATAGCGATGACTTCTTCAGCTATGCGATCGCGTTGCACCAAATAGACGTGCCGACCACCGGAAACTTCAAAG
AAAGCCTCGATGTCTCCAGGATAATTGTAGGTGCCCCAAAGGGTACATTCCCAGGAGGACTGAACTACACTCA
TCTTGGTGAACCACCAGTGAACACCACTGGTCTTGTCTACCTCTGCCCCATCTTGAACAGCAGCTGTGAAGGG
CTTCTGGGAAATGGAATGAGCTGGGATCGGAAGCTATTTGATGAAGATCCTAATGTCCGTGCAAATGAATTGT
TGGGGATTCCTTCCACTGCAACACTTGAAGACAAGGAGCGTCAGTTCATGGGAGCCTCAATGGATAGCACTGG

AGACATGTTTGTTGTGTGTGCTCCACTTTGGGTGAATACCTTCAGACATACCCAATCTGACCCAGATTACAGA
CCACAGGGACGCTGCTACTATTCACCGCGCAATCTCACCGACTTTCATGTCATTCAACCTTGCAATGGTGGGA
CTGTCACTAGCAATGAGGGAGACAGCCAGTGCACGGCTGGGATTGCTGTGACCACGCTGAATGACACTTTCAT
TCTTGGTGCTCCAGGCCAGTCCTCCGGCTCTGGTGCCCTCTACTACACTCCACAGTTGCCTCAATACTCTAGC
CCGCCACATGGTGGAGCGACCATACTGACATTCACTGAGTCTTCGTTCTTCTCGATTGGTCCTACCATCCCGC
TGTCCACCTACCAAGGATTCACTTTGGCAACAGGAAATATCCTTGACAAGGTCAAGAAAAATGTGGTGACTTC
ATATCGTCAGTTTGTCGGGTCTTCTTATTATGAACTGTTGAAGTTCTATAACGCAAGTGATCCGTATGTGTCC
ATCATGACACTACCGATGGGAGAGAGTCCCACAGAGAACTTTGGCTATTCTCTTGTAGCAGCTGATCTTAATG
GAGATGGCTGGGATGAGATCATTGCCGGTGCACCAATGTACAGCACATCATCGATGCTGGAAATCGGACGAAT
CTACATCTTTTCCAATTATGACGGCACCTTCATGAGCAATGCAACTGTCATTGTTACGGGCACGATCTCCCTA
GGTCGTTTCGGGCATGCCATTGTAAACCTTGGGGACATCAATGGAGACAATTGTGATGATCTGGCCGTCAGTG
CGCCCTATGCCTCTCAGAATGGTACCAGCTCCAGCTCTGGTGTAGTGTACATCTTCTTGGGCTCCAATGCCAA
CCTACTCAATACCGTGCCATTTCAGACGTTGGATGCAGCTAATGTCATGAGAGCAAACCAGCTGCCTGAACTG
AAGAGCTTTGGCTTCTCTTTGGCTAGTGGTGTGGACGTGGACGGCAACCTGTACAATGATCTTGTCATTGGTG
CTTTGTTCAGTCAAACTGTGGTCCTCTATAGGACGCTCTCCATTGCACTGATCAACGTGACTCTCAACGCACC
TGACGATGTCAGCATTGTCATGCAGAACTGCTCTGGCTACGCCTGTTTCGTGGTAGGCATTTGTGCTTCTTAC
TCGGGACGAGGACTTGCTGGACCTCTAGGTCTCAATATGGAAGTGAGGGAGGTTGTCAGCGGCGTTCAGCCAA
AGCGACTCTTTTTTGGCGCTCCCGTTGGCAAGATGGATGCCTACGTGAGCCAGTTTCAATTCCTGAGCAGTGG
GGGTGTCTATTGCCTCAGCGTTGTCAGCTACATAGAGAGTTCATCGGCTGGAACAACCAGCCTTCCGTTTGAA
GTTCAGGTTCTCTTTTCTCCAAATTCCCGCAATGTGACCTCGAGTGGGCCACGCCCCCTCCTTGATCTGAGGC
AGTACCCTGAGATAAACGTGACCGGCAACAACACTGTTCAAGTAGCCATCAGAAAGAACTGTAACACAGCCAC
TATGTGTCTCCCTGACCTGGACCTTAGCTTAAATCAGATTGTGTACAGCTCTGGATCAGGCTCCACCCCCAGC
TTGGTAGCAGACGTGACAAAGTACATCAATATGTCTGTGAAGTGTGCCAGCGACAAGGATGATGCTTTTGGCA
GCACATTGACTGCCACCTTCCCCTCCTATTTAAAGCTCACTGGTCAAAATGGGCTCCCCTCGTGTGCCACCTG
CTGGATTTGCTTGTTGCTAGGCAACAGCACCACCTCCTGCACGTTTGAAACATTTCTCCTGCGTGCAAACGAT
TCTATACAAGTTGAGCTGAGGTGGTCTGTTGAAAGTGCCACACTGCTGGGCAATGAGAAGTTCAACATATCTG
TTAACGTCTTAGTTCCAAACGACATTCGTACCAACAACAATGAAGTCACTGTTCCAATTGCAGCCACAGCTAT
GGCAGAACTGTCTTTGGAATTAGCTGTGGACCAAACCTCACTGCAATACTCTCTTGCTGCAAATTATTCAGCC
ACAGACACTTACAGTTTGAATGACTTTGGGACCACACCTTCCAAATTGACGATGTCGTTTCTAAACAAAGGTC
CCTCCACCATTCAGAACCTTCTCCTTGACATCTACTACCCCTACCAATCAGAAGACACTGGAAAACTGTTTTA
TCTCTATGCTGGACCAGCGAGTAATGAAACATTCAATACGGGCATACACGTGAGCTGCAAGGGTGATAACCCA
AGGGGCATCCCCTCTGTCCCGGCAAGAACGCGCAGGTCGGCCGAACGGTACACGTCGTTGGCCCAGTGGTGGC
TGGACCAGGCCTCGGGGCTCTTGCCCAAGAGGAGGAGCGTCCCTCCCTCGGCTGTCGTGTCTGGCACTGTGGA
CTGCTCCGATTTAACCCAAAGAAGTCATTACTGTGGTCACGTGAACTGCACCATAAAGTACCTCCTCGTGCCT
GCAGACAGGACCAGCAAGAACCTCTTCATCAACCTACCTCTCTACGTCGATGACAGATACTTTGCAAGTAAGA
AAGGAAACTACAGCCTTGTAATAGGTGCCCAGGTAACCATCCTCAACAGCTATATACAAGACACTGCTTCCGT
ATCAGGAAAAACTTTCTTGACCTCGCTTACATTGAGTCCCAGTCCAGTCGTTGAGACAACCACGCCCAACGCT
CCAATCCCTTGGTACTTATACGTGATTCCTGCTATTGTTGGCTTGGTGTTCATCATCATCATCGGATGCATAT
TGTACTTCTGTGGGTTCTTGAGGCGGAAGAGGCTCATTCCACCGGAAGCAGATCAACCACAGTCTCCACGCGG
TGTACCTCAGTCCGCACCTGGCACAGCAGGACAACCTACTCCCAGCACACCAGGACAGCCCACTGCACAGCCA
TCACAACCGGAAGGACCCAAAACTGGAGAAGCATCTTCACCGGAAGATGACTTGCCCGAAAAAATTGATGATG
ATGCATACGAGTCAGAAATCTGATCAGTCCAGAACAGACCAACAAACAAAATATAAGCTCCCCATTGTTAGAA
CCGAACCATTTTACAAGTGAAGTGGCCCTATTACTTTTTTTTTTTTTACTTTTGCCATCTATTTCATTGCATT
TTATAGCATTTTGTACCGTACTAATCGAAATAGCCGAGAAAAAAAA

>EmITGA4_(comp62999_c0_seq1)
CTGTAATACTATTATTATAGACGATTGCGTTGATTGATTTCTAATTAGTGGATTTTCGTTTGCAACTGTTAGC
GCTGGGGTGCGCAGCTGTTGATCGTTTCTCTGGCCATTGTTATTGCTAATCTGTCTTGGGGAGAAAAATGAAA
AGTACAGCGCTGGAGGGTCCTGTCCTATTCCTTCTGCTCGTGGTGGCGGCGGCCGCATACAACGTCCACCAGA
AAGCTCCAGCCAAGGCGGTGAAGGCTGTCGGTGGCTTGGGCGAACTGTTTGGATTCAGTGTTGCCTTGCATCA
GTTTACCAACGGCTCTACCGTGAATCTCGTGGGTGCGCCGAAGCTTTTTGCTAATAGCAGTGGTGTGAGAGAG
GGCGGTGTGTACGTGTGTCCGACCGCCACTGGGAGCAGCTGCTACCTGGAACCTCTCTTCAGCAGCAGACTGG
ATGCCCGCAATGATCCTGGTCAGTTACTTGCTGATCCCGCAATTTACCCTGAGAATAATTTCGATGGACAGCT
GCTAGGATACACACTACGGAGCTTTGATGATCACGTGATGGCATGTGCGCCCCTCTTCATTGGACAGAGGAGC

GGGGTGAAGGCCAGGTACACAGGTCGCTGTGTCCGTCTTCCACATGACTTCCAGCCTTCAGATGTTCCAGACC
AGATCACACTGGGCATATTTCAAGACCCGCTGGAGGGTTTTCTGATGGGTCTGGGCGTGGCACTTCTGAGTGA
GGACGCAGTCAATTATAATTTTGCCGTTGGATTTGGAAAATCAAGCAACGCCCCAGGTGGAGTTTCATACAGC
ATAAGTGTGGCAAAGAGCACAAGCACACTGTCTGTACCAGGTGGCAAATTCTATGTCACCCAGATCTACGACG
ATCTCCTAGGGTACCACGTTGAGGCTGGTCACGTCACCTCTCCCACTTCCTACGTTGCCATCATCGGAGCACC
TCGTGGCAGCAATCATTTTGGAAATCTATTCGTAACTGACTTGTCACAAGCAAACCCTCTTACTCTGTTTACA
GCCAAAGGAGTGCAGTCAGATGGCCACTTTGGATTCTCCTTTGCAGTTTGTGATTGGATGACTGATGGTTATG
ACAGTCTTGTGGTGGGCTGTCCACTGTGTGACAATGATGTGGGGAGAGTGTACGTGTACTTGCACAGCGGCAA
TCCAGCCAATCCGTACCCTACTGTACAGGAAGTCACGCCTCCTACCATCGTAGCTGGTCGTTTTGGCCTCTCC
GTAATCAATACTGGTGACTTGGACAAAGATGGCTACGATGACGTTGCCATAGCAGCCCCTCACGATGCAGGGG
GCGTGGTCTACATATACCGAGGCTGCGAGTCAGGTCTATGTGACAGTCCACAAGTGATCAGACCAGCAGCAGT
TACTCAGACTTCCTTATTTGGGTACAAGCTCTCTGCAAAAGTGGACGTGGACAACAACAGCTATCCAGACTTA
TCTGTGACGGACATGTCTGGCACTGTGTACACATTCAGGACCAATCCTCTTGTTGTGGTTGACACAACGTTTG
AAGGACTCGCATCCACCCTGAACATTAACACAGACATATGTAGTGTTTCAAGCTTTTCAGCTGCATGCTTCAA
CTTCAGTGTGTGCTTCGGGTACAGACCTCTTGCCGGAGGTGAAGACATTGGCACATTTGCATTCAGCTACACC
CTATCAGTGGATCAATCATTTGGTCGTGTGGTCCAGAAGTCCACACTTCCAACGTCCAATCAAATCACTCTAA
GCAGAAACACTTCTTACTGCACAGCTCAATACTACTACTTCCTGAAGCCTGATTCCAGTGACACCAATTCTCC
CATTACGGTACAGCTGACACTCCAGGACGTAACAAATCCCCTGGCTCTTTCATCTAGTCCCCAGTCAACTAAG
TTTGGGGGCTCCCTCACTCCCCTCCTCGATCGTTTCGTAGCTGGAAGCCAACCGCGTAACATTGTGAATCAGT
CGATCTCCCTCATCACCAGCTGTGGCAGTGGAGGAATTTGTGTGCCAGAATACGTGATCCGCACCAAGAAAGA
AGGGTCTGAGAAGGTTCTAATCTCCAGTGAGCCTTTTTTTGTGGTCATTGCTATAGAGAACTTGGCCAACCAG
AGTGGTATAGCACCCCAGCTGTTCATTGATGTACCTTCGGGTGTTGGAATCTATGGAGGGAGCTCCAACGTGA
GCACCATTGCCGGTGTGCAGCAGTGCGTCCCCCTGACCACCGCACAGTTCAAATGCTCTCTCAGCCTCATCCA
ACCTTCTTCCTTCCTCAACCTCACCATCCCTTTCATCCTGGACTCCGCCATCTTGGGAGTCAATCTCCTCTCT
GGCGAGTCTGTCCTCCCCACCCTCACCATCAACTTCACTATTGGGCAGAACAATTCTCGCGCAGGCGCCATGT
CTTCTTTGCCTCTCATCCTCGACGCACAGGCGCAGTACACAGTGACCTCGAGCACCAATGAAGCCGAATATGT
AGCCTATTCCCTGACATCATCATCTCTGTCGGGACAAGGTCCTGCCCTCAAGTACACAGTGAAGGTCACAAAT
AGTCTCGTGGCAGGTACCACCATTCCCAACACAACACTATACATCTACTGGCCCACATATTTCAAGTCAGTCA
AGGGCCAAGTGCCACTCCTGGTGGTTACTCAAGACACTTCTCCGAGTGCTCCTTGCAAACACTACAACACTGT
TCCCCAAGAGGCCCTCCGTTTCAGCAACGCGTCCCTCCCTCAACAACACAACAGCACCACATACAATCTGCCG
AGCCAGTACACCACGGACAGCAAGAGCTTCACGTACGGGGTGATAGTCTGTGACATCAAAAATCTGGCGCCAC
AGTCGTCAGTCGATGTCATCATCCTGAGCCAGCTGTGGAGCACATCAGTACTGACTTCTCTTATGCTGAGTAT
TGAGGCAGCTGTTGTCAGTGGTACATCGCTGCCAAATTTCCTTGGAGGAAAGCCATCAGACAGCGTCACTGTC
CTGCTTAGTCCTGGAGTGAAGGAGTTTCCTGTGGCATGCTTCCCTATCTGGATCATCATTGTGGCTGTCATTG
GAGGTTTCTTGTGCCTTGCGCTCCTTGGCACATTGATTGCCTGTATCTTCCTGATCTATCGTTTTGTTAGAAA
GAGTGCCTCTTACGACCCTAATGCAGACAACCCTGACAATGATGATGATTTTGTGTACAAGCAGCGAACAATA
CCTCCCATTGCAGGTTTGGCAGAAAGCAGACCTACGTCTTTCCAGTATCTGTCAATGACGGGAGACCATGCTC
CAAGCCAAGACCAAGAGACCATTAGGAAGGAGAAGGAGAAAGAAATGGAAGAGGAGTTGCAACTACAAGCAAA
GCTGACACACACCATGACTCGGCTAGTCAAGCAGCCACAATCCAAGGAGCCAAGCGAGGCTCCAAGCGATGAC
CCATTTGAGACAACACTGTGATCAGCTGACATTTTCATTTTATCTTTTCATATTTGTTTTTCTTCTTTTTTTT
TCCCTCACAATATAACATATTGATTGTAAAAAAAAAA

>EmITGA5_(comp68292_c0_seq1)
GATGCTTAGAAATTGTATTTATAGATCTCAATTGCATTCCCAGCGCTCGTTCGTATAGCCTGCTATAAGATAC
TGCTATCCTGCCGCGTGGATAATCGCAAACAACATGACTCCTTTCCTTGAACCGCTGGCGAAAACTGTTCACG
TCCTTTGCATTGTGATGGCGTGTTGCCACGCCCAGAGCATCGATACCAAAGAACCTATCATTCGGACGTCCCC
TGATCTCACGAGCACAGACTACTTTGGCTACAGCGCGGTGCTACACCAAACGCCTGTAGCTACGGTCGTACTC
ATAGGAGCTCCAAACGGAACGGCTCCGGGCTCAGCAGTGAATAATACCGGCTTGGTGTACGTCTGTCCAGTGA
CTAATCCAGGCACGTGCGCAGGTCTGACAACATTTACTAGATCCAGTCTTGCCACAGACACACTGCTTTATGA
CAGAAGCAACAACAGTGCGAGCGAACAGAAAAGCGGACAGTTTCTAGGAGGGACGATCATCAGCAAAAGAGGA
TTGGTTGTGATCTGTGGACACCGGTACTTCAAGCCACAGTACACCCCCAATGGAAGGTGCTTTGTATCTAACA
GCAGCCTGATAGGTTTTCAACAGTATGCGCCATGTTCCTCAAGTGGTGACCGGTGTCAGACTGGTGCTAGTGC
ATCCATTGGTAATGCATCAGGAGTAGCATTTCCTCTCCTAGGATCTCCAGGACACAATGGTTGGAGTGGCACT
GCATCCAGGGTCACTGTGGCATCTGGCGCATTAAGAACCACAACTCGTCTGTCAGGAACAGCAGAATTAGGTT

ACCAAGGATACACTGTGGCCAGTGGCCATATTCTTCAGAAAACAACTGAAGATTTCCTGGTCTCTGTACCGAG
GCTCAATAACATGGGTGTGGTCAACTTAGTGATGAACTCTGCCACTGTGACTGTTGTCTCGCAGCCACTTCAG
GGAACACAGTTGTCTGAGTACTATGGGTTCTCCATCATCACAGCAGACCTTACTGGTGATGGATATGACGAAG
TTCTGGTGGGAGCACCTTACTACTCTCCTGTTCAGAACCCTGAAGCTGGAAGAGTCTATGTGTATAGGAACAA
TGCAGGAAGCCTTCAGTTTGTCAAGCAACTCTGTGGGAGCGCTGAGAACTATGGAAGGTTTGGGCACGCCATG
ACCAACCTTGGAGACATTAATGGAGATGGACTGGATGATGTAGCCATAAGCGCTCCTTTTGCCAGTGGAGGAG
GAAAGGTGTTCATCTATAATGGAGTTGTCACCACGACCATTAGCTCCACCCATTCCCAGGTTATTCAAGGAAA
TGCACTGCAATCGACAATAAATCTGTTCAATCTTACCAGTTTTGGGACTTCTCTGTCCAGTGGTGTGGACATT
GATAAAAACACTTACAATGATCTGGCTATTGGAGCTTATAATAGTGGGCAAGTGTTTATTTTAAGAACACGCC
CCACTGCATTGGTTGCGGTGTCACTAACAGCAAACCCTACTCTTGTTCAGCTGACCAATGGACTGTACCCGTC
TTGTAATCTGACTGGGACAACTTATGCATGTTTCAACCTCACTGCCTGTCTCACCTACACTGGAAATGGAGTC
AGTAACCTATTAAATCTGACAGTGACCATTATTGGAGACACATCAAACCAGGCCCTGGGCCTTGCATCAAGAT
TATTTATTGGAACCAACTCTAGTATCTCAACCTTTGTAACCACAGTTGGCACTACTAAAAATATCCAGTCATG
TCTATCATTACCAGTATACATCAAGAATGAGATCCTTGACAAACTGAATGGATTTACTATGCAAATGAATGTA
TCAGTACAGGACTTTGCACCACCATCTCAGAATGGGAATGGGACTCTCACCAATCTGACTGGCTACCCTGTAC
TCTCTGTTCAGGGCACCAGCTCTGTGCAGGTGACAAACATCGACAGAGGGAACTGTAGTTCTGGCACCTGTAT
TCCTATTCCCAATCTGGCCATCCAATTTTTGAACATCTCATATGAAGTTTCCAATGGTACTGGACAGTCACTA
GTGGCCCAGCAAACTACAAACTTGAACCTTTGGTTTAACATATCCAATAGTGGACAAAACGCCTTTGCAACTG
TTCTCACATTCCTAGTTCCCAAATCTCAACTTTACTTCATTCGCTGTGATCCAAATCTGGCATATCTGTCAAC
AATCCAGGACTACTCCACTGCACTGTTCCTCTGCACCTGTCAGGTTGCCAATCCTCTGCAAGGTGGCAATTAT
AGTGTAATAGCAGTTCGATTGGAGCCAACTGCCAATATCGACGTTACACAAGGAGTCATTCCACTCATGTTCA
ATGTCTCCAGCCAGAATCCTGAGAACCAGACAACTGTGCAAGACAACTCTGTGTCAGTGCAAATGAACATTAC
AGCTGCCAGTGGACTGTCTGTTGATCCGATTGGAATTGCAAGACCAGAACAGATTATCTTGAGTACAACAACA
AACACCACATCAGTTAATCCATTGGGACCTTCCATTTTGACCACCTTCACAGTGAGAAATGCTGGGCCTTCAA
CAATACCATTAGTTCAGCTGAACATATATTGGCCTCTTAACAGCTCTGAGACTGGAAGCTATTACTACCTTGT
TCCTACCTCATTGCAAGCATTGACCACAACGTTCTTCACTCAATGTGACACAACATATCTGAATCTGATTGCA
TCAAGTGTCAATGCTACTCAAGCTCCATCATCCTCCGGTGGTGGAAATAAAAGAAGGGCAGCTAGTTTGTCAA
CATCAACAGCACAAGACATTAATAAAACTATTGACTGTCAGCTGACCCCTTCTTCATGTGTACGGATGCAGTG
CAATATTTCCCAGCTCATCCAAAGTCAGGTGCAGATAACCATCAATGCTGCATTGGATCTACGCTATTATACA
GCTGAAAATATAAAGCTCACCTTCATTCCTTATGTGAACGTTTCCATAGAAGGAAATGGAGCTAACTACATTG
TTCAGACCTCCTCAATGAAAAGTGCAACTGCAAACATTAAGGTGCTTAAACGCACCACCGCAAGCAGCGATTC
CCAAACTGGAAGACCTGCATGGGTTGGATGGGTGATTGGAGGAATCTGTGTCCTGCTAATTCTGATCATTGTT
GGGCTCATCGTGATTGTGACTGTTGTCTTCATCAAAAAACGGAAGGCCATGAAGGATAAATTTGGGGATGAAC
TGAAATGGAAGACAGTTCAAGTTACTGACAACCATACTGATAGTGTTCGTCAGCTGATACCAGACAATTAAGA
CTGCTGCATGTGACAATATGCAAGTTTTTCTTTCCAATATTGTAACAATTGCAACTTATTAGATGAACAGGTG
TGGAGAAAATGTGTCACTTATTCAGTACAGTTGATTTCCTCATTTCTTACATTCTGTAATGATTAATGTGTCA
TGCACCAATAAGTAGTAGCATGCAAGAACACATGTGTGTTGTTATCAGCATTTTTTCATGTATGCATAGTACT
AGCAAAGTCCATTACTATACCTAAGTACTAGCTCTACT

>EmITGA6_(comp60005_c0_seq1)
CATCACGATCCGATTTTTGTTGCTGAGAAGGAATTGCACTGCACAGACGAGCGAGAGGTTGCCATGGCAAAGG
CTTCGATCTACAGTAGTGCGATATACCTGGTACTGACGACGGTTTGCACGTCCGTGACCTCGCCTCAGACCAT
CGATAGCAAGCAGCCCATCATTCGGACGTCTCCAGACCGCACAAACACCGACTACTTTGGATACAGCTTGGCT
CTACACCAAACGCCTACGAGTACAGTTATCATTATTGGTGCTCCAAACGGAACGGCTCCGGGATCGGCAGCGA
ACAATACTGGGCTGGTATATGTGTGTACGCTAACTCCCGGCACGTGCACTGGTCTGCCTGCATTCACGAGGTC
AAACTTGTCAACCGACAAGTTACTGTATGACACGAGTGGAAACAGTGGTGAACAGAAAGCGGGGCAGTTTTTA
GGAGGAACAATTGTCAGCAAGAGAGGTCTAGTTGTGGTCTGTGGACATAGGTACTTCAAACCACAGTACAACC
CCACTGGAAGGTGCTTTGTATCTAACAG

>EmTalin1_(comp28662_c0_seq1)
AAGAAATTGCGCATGCGCATAGCCACCAAATATCGTTTGAGAAAGAAAGTTTGTTTTTCGCTTGAGGACCTTT
ATTCAAGGCGGCATAAGAGATGGCGACCCAAACGGTGTCGCTAAAGATTAACATCACCAAAACAAACAACATC
AAGACGATGCAGTTTGAGGAATCAATGATGGTCTTTGATGCATGTCGTCTGATTCGTGAAAGAGTACCTGATG
CAGTGCAAGGCCAACCCACTGAGTGTGGGCTGTTCAAACCAGACGAGGACCCGACCAAGGGGAGGTGGCTAGA

GATGGGGAGGACCTTGGAGTACTACCATCTCAAGAGTGGTGACATGTTGGAGTATCGTAAGAAGATTCGACCA
CTCAGAGTTCGAACGTTGGATGGCTCGATTAAGACGGTTCTGGTCGATGACAGCAATACTGTAGCGGAACTCA
CCAAAACTGTCTGCTCCAGGATAGGCCTGGCAAACCACGAGGAGTTCTCGTTCACTGTTGATGAGGAGACGAG
TGAAATGACTCTGAGACGGCAACACACCCTGGCCAGGGACCAGAAGAAGTTGGATAAACTGAAGAAGGAGCTT
CATACAGATGATGAGTTGAACTGGCTGAACTCAGACAAGTCCCTGAGAGAACAGGGTATCAGTGAGACCGCAG
TCCTCACCTTGAGGAAGAGGTTCTTCTTCTCCGATCAGAATGTGGACAGGAACGACCCAGTGCAGCTCAACCT
CATCTACGTGCAGTCCAAGAATGCCATCATTGATGGCACTCACCCTTGCACCAGAGAAGAAGCAGTGCAGTTT
GCCGCTCTGCAGTGCCAGATCCAGTATGGGAACCACAACGAAGCGAAACACAAGCCAGGTTTTCTCAATCTGG
ACGAATTCCTCCCAGAAGAGTATGTCAAGTTCAAAGGAATCGAGAAGCTCATTTGGACGGACCACCGCAAGCT
GCACAACCTCACAGAGCTGAATGCCAAGTTCCGTTACATCCAGTTGTGTCGCTCCTTGCGCACATACGGAGTG
ACCTTCTTCTTGGTGAAGGAGAAGCTGAAAGGACGCAACAAGCTGGTCCCCCGACTCCTGGGAATCACCCGGG
AAAGCATCATGAGGGTGGACGAGACAACGAAGGAGGTACTCAAGACCTGGCCTCTGACCACTGTGCGTCGGTG
GGCCGCCTCCCCCAACTCCTTCACACTGGACTTTGGGGACTATTCGGAGTCATTCTACTCTGTGCAGACGACA
GAGGGGGAGACCATATCGCAGCTGATTGCTGGCTACATCGACATCATAATGCAGAAAAAGCAACCCGTTCAGA
ACGACATGGCAGACGACGACGATACAGCAGTGGTGGTGGATGATGAGGTCCTGCCCAATACAGCTGTAGCATT
CCGATACACTGGAAGCAATCAAGGCACAGGGGAGTTGGACAACACAGCTAACATGGTCATGCCCCAGCAAGCC
ATGACTGATGAAGGCGCCATGTTCCATGCCTCTGGCTCCGCCCAGTTTGCAGAAGAGATTGGTGCCCCTGGGA
AAAAACACCAGCCACATCAAGACAGCCAAACCGCACAGTTGCTTGCAGCCCAGCAAGGCCTGCTTGCCAACAT
TGGATCAGCTCAGCAGGCCATAGGAGCCATCAACAAAGACCTTCTGTCTCAAGCACAACTGCCACAGCTGGGC
TCAGACTCTGCTTCCATCAAATGGAAACAGACCACCTTGGATGTGTCCAGGCAGAACGTCAGCTCTGCAGTGG
CGGCCATGTTGGCCTCCACAGCTTCCATCATCACCCTGACCCAAGGGGATCCTATGGACACCAACTACACTGC
TGTGGGGTCAGCAGTGACCACCATCTCCACTAACCTGACAGAGATGGCTAAGGCAGTGCGCCTGCTGGCAGCA
CTCAGTGCCTCACAGCTCGAGGGAGATGACCTGCTCAAGGCAGCCAGAGCTTTGGCAGCTGCAACTGCCGCAC
TCCTGAACGCAGCACAACCTGAAAACATGGAGAATCGACAGCAGCTGTTGATGACAAGTGGTGACATGGCCAT
GAGTGGCAGCCAGCTGCTGGGTCTGGTAGGGGAACAGGAAGTGGATCAAGGGACACAGGATGCACTGGTTGCC
ATGGCAAAGGCCGTTGCCACGGCCACCGCTGCACTTGTCACCAATGCAAAGAATGTTGCAGCAAAGTGTGACG
ACCAGGCTCTCCAGAACCAGGTGATTGTGGCAGCCAAGCAGACTGCCCTGGCCACCCAAGGACTCATAGCCTG
CACCAAGGTACTAGCTCCATGCATCAACAGTCCCCTGTGTCAGGAGCAGCTGATTGAAGCGTGCAAGCTGGTG
GCAGCAGCAGTAGAGAAGATAGTACTGGCAGCTCAGGCGGCGTGCAAGGATGGTGATGCCCTTCGTGACCTGG
GGGCAGCAGCTACAGCAGTGACCACAGCACTCAACGACCTCATCCAGCAGATCAAGGAAGGAGTGCGGATGGA
GGCGGGGCAGTATGACGAGGCCTGTGAGGCCATCTTGGCTGCCACTGACAGGCTCTTCAGTTCCATGGGCAAT
GCCCAGGAGATGGTGAAGCAAGCCAAGCTCTTGGCTGAAGCCACGTCAGCTCTTGTCAATGCCATCAAGCTGG
AGTCAGAGAATGAGAATGATCCAGACGCGAGGAGGAGACTTCTCGATGCAGCAAGAGCCCTTGCTGATGCCAC
CTCCAAGATGGTGGAGGCCGCAAAGGGTGCGGCCCGTAACCCTGGCAACGAGCAGGCCCAGGAAGCTCTGCGC
AAGGCAGCAGAGTACCTTAGGGCTGTGACCAACGCAGCAGCCTCCAATGCCCTCAAGAAGAAGGCCATCAGGA
AGCTCGAGATAGCGGCCAAGCAGACAGCTGCTGTGTCCACACAACTGATAGCAGCTGCACAGGGGGCTGGTGC
GTCCAATCGCAATGAAGCGTCCCAGAGCCAGCTGATCAGCCATTGCAAGGCAGTGGCTGAACAGATCTCCCAG
CTTATCCAGTCTGTACGTGCTAGTGTGGCCAACCCTGACAGCCCTAGTGCCCAGCTGGGGCTGATCAATGCCT
CAATGAACATGATTCCTCCTGCGGGCAAGATGGTGGCAGCTGCCAAGGCAGCAGTGCCCACAGTGGGGGACCA
AGCTGCAGCCTTGCAACTTGGGAACTTTGCCAAGGCCACTGCCTCTGCTCTGGCTGATCTGAGGACTGCAACA
TCCAAGGCCTCTGAGATGTGTGGATCGTTGGAGATTGACAGTGCCATTGACACAGTACGCAGTCTATCCCAGG
AGATGGGAGAAGCCAAGATGGAGGCCCAGACAGGACAGCTATTGCCCCTCCCTGGTGAGACGGTGGAGAGCTG
TGCACTTGAGCTAGCTGCTACCTCCAAGACGGTGGGCTCGTCCATGGCACAGCTACTCACTGCCGCTAGTCAA
GGTAACGAGAACTACACTGGTATGGCTGCCAGAGACACAGCCAGTGCTCTGCGTATCCTTGGCAACGCTGTTA
GGGGTGTGGCCGCCGGAACGAAGAACCGTCAGACTCAGGAGTACATCCTGACGACAGCCCAACAGGTGATGGA
CCAGAGCTGTGCCCTGCTGGTGGAGGCAAAGGCAGCTGTGGAGGATCCCAATGCCCCCAACAAGCAACAGAGG
CTTGCTCAAGCTGCCAAGGCCGTCTCCCAGGCCCTCAACCAGGTGGTCAACTGCCTGCCTGGACAGATTGAGT
TTGACCAAGCCATCAAGGCTATTGCTCAGGCCAGCCTAACTCTGCAGGCAGAGAAGTTCCCTGATGCATCTGG
TGCAAGCTACCAGACGCTGCAGAGTAACCTCAGTTCAGCAGCAGCTGCTCTGAATGCCACAGGAAGTGAAGTG
GTGGCAGCAGCCAGGGCTACTCCAGAACAGCAGGCCATCGCCACAGTGAAATTTGCACACTGCTACGAAGAGC
TCCTTAAGGCTGGCCTAACCCTGGCCGGAGCATCAAAGGACAAGGAGAGTCAGAATGAGATGCTGGGATACCT
TCGTAACATCAGTGTGTCCTCTTCCAAATTGTTGCTTGCGGCCAAGGCACTCTCTGCTGACCCCAATGCGCCC
AACGCCATGAACCAGCTGGCAGCAGCTGCAAGGACTGTGACAGACGCCATCAACTCCCTGCTCAACCTCTGTT
CCTCCTCGGGCCCAGGCCAGAAGGAGTGTGACAATGCTCTCAGGAATATTGAGGCCGTGGCGCCGGTTCTGGA

CAATCCCAACGAGCCAGTCAGCGAGCTATCTTATTTTGACTGCCTTGACATGGTTATTGAGAAGTCCAAGATG
CTTGGAGAAGCAGGTACTCTGATCACCTCACACGCCAAGAAGGGTAGCATCGAGGAGTTTGGGAAGGCTGTTG
AGAGCACTGCTTCGGCAGTGTGTGTGTTAACAGAGGCAGCCGCCCAAGCTGCCTATCTCGTGGGCATCTCTGA
CCCCAGTAGCACGGCAGCCATTCCTGGGCTGGTGGACCAGAACCAGTTTGCAAGGTGCAACCAAGCCATTGCT
ACGGCATGCCAGACCCTGCTCAGCACGTCAAGCACTCAGCAACAGGTGCTAGCGTCTGCCACTGTCATTGCCA
AACACACCAGCTTGCTTTGCAACGCTTGCAAGCAGGCGTCAAGCAAGACAAGCAACCCTGTTGCCAAGAAGCA
CTTTGTTCAAGCAGCCAAGGAAGTAGCCAACAGTACAGCAAACTTGGTCAAGAACATAAAGGCCTTGGCTGCT
GATCTCTCTGAGGAGAACAGGCAAGCATGTGCCTCTACCACACGACCACTCCTGGAGGCTGTTGAAGCTCTCA
CCACCTTTGCCTCGTCTCCTCAGTTTGCATCTACACCAGCCAGGATCAGTGAGCAAGCACGTGTTGCCCAGCT
GCCCATTGTACAGTCTGGTAAGAACGTGATCAAGTCTTCAAGCAGTCTTCTCACCTCAGCCAAGAGTTTGGCC
ATTAACCCTCAAGATCCTCCCATGTGGCAGCTGCTTGCGGCACACACAAAGGCCGTGACAGACTCCATCAAAG
CACTCATTCTAGCTATCAGAGACAAATGTCCTGGCCAGAAGGAATGTGATTCTGCTATTGATGGCCTCAATGC
TACCATCAACCAGCTGGACCAAGCCATCCTGTCAGCCATGAACCAGCAGCTACACCCCAATGCTTCCAGCAGC
CTGCAAGGGTTCCAGGAACAGCTGCTCCAGGCTGTCGGGGACATTGGGGAGCACGTCAAACCCATCGCCACGG
CAGCCAAGGGGGAGGCCGAGAAGCTAGGGCATCAGGTGACCGCCATGTGCAACGTGTTCCCTAGTCTGGCAGG
GGCAGCCATTGGAGCTGCCTCAAAGACCACCAGCTCGCAGCTGCAGATAAGCCTGCTCGAGCAAACCAAGACC
GTGACTGAGTCAGCCTTGCAGCTGGTGTACGCAGCCAAGGAGGCCGGAGGGAACACCAAGTCCACAGCTGTGC
ATGGAAAGGTGGATGAGGCAGCCATTCTTGTGCAGACAGCCGTGAGTGAACTGACCCAGACACTGGAGAAGGC
TGGAAGCGAGACCGGCATCATTACTGCTATGGTGGATGAGATCAAGAAAGCCATGGCCCGTGTGCAGGAGTCA
CCTGGAGAAGTCTCCAAGACCTTTGCAGACTATCAGACGGACACATTCACATATTGCAAAGCCATCACAAAGA
ATGCTCAGGAAATGGTGGTGAAAGCTTCTAGCGTGTCTCAGGAGCTTCCCACATTCAGTCGGGAACTCACCAA
CGCCTACTCCCAACTCGTAGACACCACTCAGTGCGCCTTGGCAACAATCGATTCACAGAATATTGCATCTCGC
CTGAGCCAGAACGTGCGAGCCCTTGGGGAAGCATGTATTGAATTGGTGTTTGCTGGAGGCACCCTTCAGACCA
GCCCTGACGACCAAGCTGCACGAAGAGAACTCACTGATAATGCCAAGAGTGTGACAGAGAAGGTGTCATATGT
TCTGGCAACTATACAAGCTGGTGCAGTTGGGACACAGGCCTGCAACAGCGCCATAGCAACCATCATGGGCTTG
GTTGGAGACCTGGACACAACCACCATGTTCTGCACAGCAGGTGCTCTCCATTCAGAAGACAAGCTTGGCACAT
TTGCGGAGCATCGTGTGAACATCCTGGAGACGGCCAAGGTGCTGGTGGACGACACCAAGAAGCTGGTCAGCAG
TGCGGCTGGTACTCAAGAGCAGTTGGCTGAAGCAGCCATACAAGCTGTGAAGACCATCACTGCTGAAGCAGAG
CACGTGAAGCTTGGGGCTGCGTCCCTTGCCACTGAAGACATGGAGGCACAGTTGCTGTTGTTACAAGCTGCCA
AGGATGTAGCCAACGCTCTGAGCGATCTGATTGGCGCAACACGTTCTGCAGCAGGCAAGAGTGTCCAAGATGC
AGCTATGGAGCAGCTCAAGTCATCAGCTAAGGTGATGGTGGCCAAGGTGTCAAATCTGTTGAAGACTGTGAAG
AATGTTGAAGACGAAGCTGCAAAGGGAGTGAGGAGTTTGGAGAATGCCATTGAAGCAATCGCATCAGATCTGC
AGGAGTTTGAGTCTTCCAATCCTCCCAAGAGTCAGGCGACAGCAGAAGATCTCATTCGCAGTACAAAAGGCAT
CACATTGGCCTCTGCTAAGGCTGTATCCGCTGGGAACTCGTGTCGACAGTTGGACATTTCGGCTTGTGCCAAC
CTGGCACGAAAGGCCGTCACTGAACTGCTCGAGACATGCAAGTCTGCTGCATACAAGGCAGAGAATGGAGAGC
TGAAAGCTAAGACCCTCATGACTGGGCGGGAGTGCGCAACATCATTTAAAGCTCTGCTGGAACTGGTACACCA
GATTGTACTCAAACCTACCTACGAGAAGAAGCAGAGCTTGCCAACCTTCTCTAAGGAAGTGGCCACTTGGGTG
GGCGATGTTGTCCAGGTTGCTGAGCAACTGAAAGGATCCGATTGGGTTGACCTGAGGGATCCCAACGTTATTG
CTGAGAACGAACTGCTCCAAGCAGCTGCCTCCATTGAAGCAGCTGCTAAGAAGCTGTCTGAATTGCAGCCACG
CAGGGAAGTGCGCGCTGATGAGTCCCTGACCTTTGAAGAGCAGATATTGGAAGCTGCCAAGAACATAGCCTCA
GCCACCAGTGCCTTGGTCAAGTCTGCCTCAGCTGCCCAGAGGGAGTTGGTGGCCCAAGGCAAGCTGAGCTCCA
AGCCTCAGTCAGAGGACAGTCAATGGTCTGAGGGCCTGGTCTCTGCTGCCAAGTTGGTTGCCGCAGCAACGAG
CAATCTGTGTGAGGCAGCTAATATGATGGTGCAAGGACACGCCCAAGAGGACAAGCTGATAGCAGCTGCCAAG
TCAGTGGCCGCTTCCACTGCACAGCTGCTCATTGCCTGTCAGGTCAAGGCTGATGCCCGCAGTGAAAACAACA
GACGTCTGCAGATGGCTGGTCAAGCTGTGAAGAAAGCAACTGAGACTCTAGTGGCAGCTGCGCAGCAAGCAGC
TGTGGAAGGTGGCAGAGCAGATGGGGCAAGTGGTGGAGCAGCATCCATCCAGGTGAACCTCGGAGGCAAAGTG
ATGAACAGATTCCGGCAAGAGCTGGAGATAGCAGAGCAGATAGCAGCAAAGGAGAGAGAATTGGAGCAAGCCA
GGCTTCAGCTGACCAAGATCAGGAAAGGACAAAACTAATGAACTTTCATTTAATCCACATGAACACTTTTGCT
TTTAGTGTAAGAAGATATGTTGTGTAGTTCAGAACAGATGGCTTATATAGCCCGTGTGTTGTGTGCCTTGCTC
ACAATTTGTACTTTAGTGAGTACCAGAGTAACTGCTTCCATGTTGTGTACGTGTGATATGAATTTTGATGTCG
GTAAAAAAAAAAAAAA

>EmTalin2_N-terminus_(comp57177_c0_seq1)

CGGATCTAGCGTGTCCTAGTCATAGTTCTATCAAAATGGCTACAGCTAAAGTTGCGATTCATATTCACTTGGT
TGATACAAACGACGTTCGAAGCGTGATGTTCGACGAGACGATGCTTGTCTGTTATGCATGCAGTTTCGTTCGT
GAAAAATTTCTCCCTTCCAATAGTAAGGATGGTTCAGCAAGCGAATATGGGCTGTTCAAACCGGATGAAGGCA
GTCCACAATGGCTAAATGTGGGAAGTATGATCAAAGACTATGGACTTCGCGACGGGGATACACTGCAGTACAA
GAGAAAGGTCCTTCCGTACTCAGTGACGTTGTCAGACGGACGTACGTTGAAATTCAAGTTGGATAACAGTCGA
ACAGTGGCTGAAAACGTCAAGACAGTGTGCAGCGAAGCTGGACTACCCAATGACTATGAATATTCATTTGAGA
CAACAAACTTGGCAGTCAAATCGTCAAAAAAGCAAGATACGAAGGCTAAGAAACGGTCCTCGTCTTCTGCAAA
GAACGTATGGCTGAAAGCGAACAAAACTTTCGCCGCGCAGGGCGTGGATGAGAACTGCATACTTACACTTAAA
AAAAGGTTTCGCATCGTTGATGCACCCATATCACTTGGAGACTCTACCGCTTTAAACGCTGTTTATGCGCAGT
GTAAAGATGACATCACCAGTGGTTTGCACCCATGTACAGAGGACGAAGCGATACAACTTGCAGCTCTGCAGTG
CTACGTAAGGTTCGGGAAGAACAGACCCGTGTCGATTAAAATAGCCGAATTCTTGCCACTCGATTACGTGGCG
AGAAAGGACGTAGAGCAATCTGTACTTTTGGCGCACTCCAAACTGAATGAGATGACAGAGGCAGAATGCAAGT
TTTTTTACATCCAGTTGTGCCGCTCTCTTCACACGTACGGAGCAACGTACTTTCTCGTCAACGAGAAGCGAAA
GGCCAAGAAAAAGTTGGCTCCAACGCTACTGGGCTTCAAAGCATCGGAAATTATCAGGGTAGACAAGGTGACA
AAGCAGGTTGTTACCTCGTGGCCTATTGAAAACGTTCGGCGTTGGATGGGCACAGAAGACTTCTTCAAGATAG
ACTTTGGCACAGCTTCGCAGCTGATCTATACAGTGCAGACTACGGAGGGTCTAGAAATATCCGAACACCTCTC
AGAACTAACTGACATAATTGAAAAAATTAAAAAAAACAACGCTGGGAACAAGAAAGATGGCGATCGAGAAATG
TGCAAAAAGAATGACACTGAGTCGCAGCTACAAACAGCAGATACCATTTGTAGTGCAGAATACGACCTGGATA
GTGGTCCCTTTGATGACGCCCAATTTTCCAATGGGTGCCTGTCTGCTGAGTACCCTGACTGTCCTTCACCAAG
GATGGACTGTCCTTCACCAAGGATCATGGACTGCCCTTCACCAAGGATCACGGACTGCCCTTCACCAATGCCA
CCTGAGGTTGGAAATCTTTTGCCAGTTACAGCTCCGGGTGAACTGCAAAATAGTAGTAGCACACCTCAATGCA
TCTCGCCATCTCCTGAAGAGCAGAGAATGATGTTGCCACCTCTTCAAATCACAACAGAGCCAACATTGGGGCC
AACTGCACATTCTGATTTGCCAAACACTGCACTAGCCGGGTACTTGACACTTCCTTTGCCACATAAAAAATCA
ACTGCAAAAGCAGGAAATGAATCAGACCCTGCAGAGGAAATACAGAGACTGCAGATGCTTATCCTGAAACTTC
AAATGGAGCTTCAGATGGCTATGGACCAAGTCAATTATGCAGAGCAGAGAGCAGAGCTTGCTGAAAAAAAAGC
TGCTCTGGCTGAAGAGAGAGCAAAAAGGGCAGAACATTCCTTTCGACTTTCATTCCAAATGACAAATCAATGA
AGTCGTCCAAGGATAGACGATTACTTACTTTGGTGCTGATCATAATATTATGTGTAGTAGAAGTACATTGTTC
CATAAAATGCATCCATTGCATAGTACATAATCCTGAATAATTTATATAGTCTTGCTATGAAATCCAAGTGAGA
AAATAGAGTATTATGCCA

>EmPaxillin_(comp26110_c0_seq1)
MELDDLLEDLQQALPPEAQAYSSSNTAKWERGSAEMSVSTPSHPQQFDEAQYSEVRRGPIYTPKSPPNSVSPP
PTKPPRTAPAASEGLSELDSLLAMLGDTQAANQEKPPHDTGLSRPTISAFVDELTQLENQVSNKTSASKFTTG
GIPMAAPVTASSATKELDDLMANLSKFEPSTVSVQDDKAPKGAQPKTRSALESLNSMLGSLEEDMSKRHGVST
MAKGTCAACNKIILGKVVNALNMQWHPEHFTCASCDAELGQVTYYESNGRPYCEKDYNELFAPRCAYCNGPIL
EKVMRALDRTWHPEHFFCTLCGKHFGTDGFHEKDGKAFCRECYYEKFAPRCKRCEKAIMEGFITALNAQWHPD
CFTCKVCNVSFPRGNYFDHEGEPHCEIHYHAARGTLCASCQKPVVGKCVSAMGKKFHPEHFTCAFCLKLLNKG
TFKEHRSNPYCQACYIKLFG*
