## Supplemental File 2 for "Diverse cell junctions with unique molecular composition in tissues of a sponge (Porifera)"

```
>EmVcl_(m.284417)
LFLSKTVQKTYEICITMPAFHTKTIHYILDPVAQQVSQLVILHEDAQQGRIMPDISAPVTAVCAAVQNLIAVG
QQTVGHSKDEILKKDLPLTLDMVESSSKMLVESATGLKADSQSKKHLELLLNGARGILQGISSLLLTFDQGEV
RKIVKSCTGVAEYIKVAEVVQTMDDLVTFTKNLSPGITSMTKMVETRYQDLTNPSHASILAAENDQVKQALPL
LLSSMKAFVTFRRDKKKGEAEAQENRNYIVQAMGESLAEIIRVLQLTSQEEVMMALAAETGSAKGTMAGGLLM
GSLAAKVHSAKELVSQTGTNAQTNKAGIQAVEAVLEEARRIAKTLPADRKAVIEGLCDELESLKKELAMLQSS
GQGDSARAHAIATALNSKMDELQMKLREAITRKVAEDFMDPVGPLNALTEASRAPLNAPARAENYVKRVTDFQ
DHSKKMADTAVALAKSGIVTDKNLADSLLLTAGKLKAVAPQVVYAAKIVYENPDSKEAKEHYDMLKEDYQKQV
QKLTKLVDSGLDTVEFLKASESMLRDELEAARTITKAGTDPQSAFKHIATAARTANRVVAVVQGESENSEDPA
FKTELASSSQAITAAIGPMVTSAKTCIQQGGSATAHHEFCVKAENLAKAVHDVYNVVDTHHNPPPPPPPPPVE
EAPERPPLPAEPEVPPRPPSPELVEVVPLQSVDPIGYAAHKLDKDAKQWEDNEMVTTARRMAKLFMQMSKFAR
GEEGEVHSKKDFINTARMIAKESEDVVKMARKVADACTDKRMKRAILQLVDKLPTISTQLKIIAAVKATRQGG
DDAAADREATEMLTDNAQNLMQAVSEVLFATEAATIRVPPDQRATLGLQWVKKGARTL*

>EmFAK_(m.70476)
MASDAVIRVFLVNGESRSVRVEEKTDSMDVIRFILRRLHVNTEYSAKLFGLQLKHTYSHECYWLQPGYTIFEL
LDQYCTTKPMEEWRFFLRIRVLPKTAQHLSSQDPVAFQYFYDQVLHLYLEDDTVQLDSETAVKLGCLELRRFY
KDMPQIALKKKENFSILEREIGLEKFFKRSLLETIPKRKLRNLVVSAFEQLETLNMDGCMFQFFNLLAKQYPL
DVEQFSNCSIGERQDGQSNDLTCTIFVGPNCDVEYQSEGGTRRFLAAFNDIHEISYEPEFVTRGKVVLELRQN
SQSIVIHTAVLVNAVHLATLVNGYCMVYTSLPHSRMTGGRRISSASRLSGMYNTVDELKMRVPSPSGEHLDDY
AEVKEHRSSLSQFGATILPKDVQVGERIGEGQFGDVHKGVLFPQTTGEQVVAIKTCKPEASDVERAKFLEEAA
IMAKFNHRHIIKLFGVMSHDSTTYIIMELALIGQLRRYLMTEGAHIAYPILLQYICQLCSAVVYLESKNFVHR
DIAARNVLLATPELIKLADFGLSKRLEDTDYYVASKGKLPIKWMAPESINFRKFTGLSDVWMFGVCCWEILMK
GVKPFVGVKNDEVINMIEMGQRLPLPPDCPAPLFDLLNQCWQYDAQDRPTFAKLEHMLMAIVEQERLEQPRKT
NSSGSSARQDPYAVIRQDGPEKPPRRDQSSSSIHGRGAQESSVRWSGFVPDEPPPPPPPRLRETDDSHLRLNE
RFRGTSGSPNGDIGSPPLEPAPYPPKPPRTDSPGSRDRIRPPDRPTPAPIVPYSVTTITPSDPTPPSFPVSPT
YPVPEPRFNGAIPPPSILRPAASRLSGDHERVGYYSVPAEAGEGPMGRVLSEDGTFVDKYGANKRGASSSSSR
LSTSSLTSQPEPKQPEPAERELDEFDDENDELLKQTTDVVRAVMEMSNKVPISRPADYVELVKNVGKALREFL
TKVEMVQKTLPIESHNEIIMANKVLSSDVTRLVDAMRDAQKNYQTFLEQEYQKQMLKAGHIIAVNAKQLLDTV
NSARRKVLRTPR*

>EmITGB1_(m.254559)
PALPFLVKLIVKLLLPIQLAIWIGEMFMTLSLAVLVALCYSEAFAQPCTDQTMCGKCLQTAGCVWCNLTTFDG
ARCFGRNVSSSMGCSSIVDPRSAPTTTTVPAATDSIYTTTPNVVLRPGDPLTVSVNVVSLPNAPLDLYILMDL
SDSMAAPLATVKSISQLIAQQVSSITTNVRIGFGAFNDKPIYPYSPQTPAGCLPGRDAPDCSDRRAGTRQYSF
LHLANFTSNFTVPNVFVTTNLDLPESSFDSLVQVLACEKELGWRNRSVEGPERGLQRLVLLITDNQPHLAGDG
RLASIYQPNDGKCHVRPYASSVGYLDTAPGILIYNEDSLLYDYPSVGLVASLLKKYDVIPIFGIVPITSSTLL
INNTFLSSYQALQDLMTSVGTKAFARPISSSASDVLDVIKTVYQEVIQNIAITLPPQSDVAVSLSQITCPDGS
ILVGQTCTNVPLSRTTTFSVTLTLLNCNTPSSSALTFSVPGFGTTTISVDKVCSCSCDKNVTVNAQQCNFRGN
FSCGGCMCIAGWTGPACERSKCGQPCVNNGTCDSATGMCQCTDYTAGPFSNDSTSGILLPPYTPKFAGSTCSC
NNFQNCPTNSQNYICSGRGQCACGSCACDATPYSWKWQGKACECPASNYSDCFDTTFKSGPLCSGNGMCSCDS
SGKGMCVCSSGYTGKYCETKITPTCDTIATCIADGTCGSLMAADSGQQVVLTCPISSGECTYSYDLSPDNQVI
RMNQVCSFAAWKIIVIVICGLLFLFVVICAIIKIILVILDYVEVRRWEKELKEADFSKNQNPLYQSPEMQYTN
VAYGKAM*

>EmITGB2_(m.284559)
VTSDCRFIAGALIKQAIVNTVNSTVQCKFFGVPRLSRERMKLGGYCCVLLSLASMSGSGNAQLCTAQTSCADC
INLSPSCKWCSTANYTGSRCFSGTPAVNCSNVENPAGTVTGIDTATLSSLVQVSARQVNVTVRPGIGTSFSLS
VQPANNYPLDVYLLTDLSYSFLDDLTTLQALGARIASSVQNISTNAQVGFGSFVDKKLAPFINILPALVNDPC
APPYGPGKCNPPYSYKHTVSLTTDGAYFSSRLQQQTISGNQDVPEGGWDGLMQAIVCKKLIGWRDNARHLLVF
STDANSHHAGDGLLGGVVRPNPHTCLMNNSITAGNVEYTLSETYDYPSLGDIREQLRLNDIIPIFAVTPDVQS
IYNAVTAELASVGASTGSLQSDSGNIIQLIQTAYQTVSQRIVFDPVLPSGVTMTFTPLNCPLLGSDNVCNGVK
PTQGAVNFTVSVQLTQDFCRANQGNTISVPVRIIGFGSFVVNISPLCGCPCQQSQISNSPFCTSNGTLTCGLC
TCNPGRFGSSCQCDANGAQSGNATSCPTGPNNLPCSGQSRGSCICGKCACSQYQDIRLGLTSTYYGSACECDN
```

SLCDTSNGQLCGGSSQGVCQCGGCQCANGFYGTACQCSNSLCVDPTDTITTRTCNGRGVCSCNQCTACKPPYT
GQYCQSCQATDKTTCASQLCPPNLDCAKCALLNQTMCPSCPTTYFVNATTLSTISGAATTECQYTDSDGCQDT
YFVVQDTKGNVTALYVRTDKACPQPLAPAQLATIIVVPLVVIAIIAILILLTLLLIFWLLNRAEVRKFEKELA
RAKYTKNQNPLYVPANQQTKNPIYEGEKAQ*

>EmITGB3_(m.284880)
MHMQHHRRNACWIVILIWLPFAYGQGACSAYTRCSDCLLENPSCGWCNDPSVFQREVGVAYQLSSLNPISLCR
NLTELSNSLNCPAQSILFPKSSNVTTYQPSSPVQPSSVVVSLRPGDSFQIPLTVTPPQSLPIDLYILMDLTKS
LEPYVNGLKTTATKLITTMQGLTSKFRIAFGSYVDKRLAPFSDRESLDNPCEGVTAAGVCNAVYDFHHTLNFT
DNASLFMETLNASNVSANLDTPDALLDALLQIALCEDQVGWSPAGQSRRIVFVMTTGDYHYALDGTLAGLVNP
PSLTCRLSPSGVYQDSELSDYPSAAVISQVLNEKRIIPIFAHIDTFATSYVALANRIKSAFLGKLLGNENNLP
QVLSNTYTTLSSTVIPVVTGTNNERYLSISVSPQNNCAPGWLQTSTTNTCANITVNTTVRYIATLTVAKEFCA
QPSNSRTVAANIQFIGFGDLRLNISVMCQPCQSCLSTDYTSSSCSSSGALECSTCVCSPNKNGPTCNCNTDPQ
AISLCRPDTTTEMCNGRGSCVCGKCVCDSVGGVQYGGQFCQCDRNKCPMGYNSKGQLAICSGNGDCFCDSCSC
NQGYTGYSCGCPTSQLQCVEPGAKSVCSNAGQCLCGICICSNATARIGTYCEECKTCTGACSNILSCVECHIT
GTCGTRCANITYVQNRTSVPGYDGTMVFRTCSITSQSCEVIYELDRYVTGLEGNVYVVLDTTQKNYATLRGNG
DCNPSQVIWPIPVGIVLGIIVVGVIALILWKACSQLGEFLEYKQWEKSLREETNRSGSNPLFVDPTSSYSNPR
YNARS*

>EmITGB4_(m.9557)
MDRWRQHRTVILQTVWFVGLLMMKVGWTASMCSTAQNCATCVSSGINCVWCSQRTANITPSCMDRSVANVSCN
LSYVEDPQVFVANLAEENLSESVLISPQSVYLNLRTGQQAVFNVSVKTSRTYPLDFYFMMDLTGSLKYDVEQV
QASGTDIANVLKNISQNYKVGFGSFVAKPVPPFVVAIPYRQPDGTCYNKEGSCIEPYAYRHILQMTNVTETFL
NILNTKLNLSSAAENPQSGTDALAQAILCKNIVGWRDEAFRMLMLISDNAVHFAGDGKAGGVVVPFDGKCHLE
WNNATGTYDYLPKYSALYDFPSVAQLKSLIADTGVSVIFGIAGANTNNTTPGTFFFPDVYKAIAKVMDIPESS
VAILSKNSTNILQVINDQYLKAIGLIKFSVPAVNGVNITVQPIMGCNSSLPSGCNDVSLEKEVVFRVTVGLTH
CPSTPQNNIVVPLRIPALAQQITIEINPNCQCACESKPAANSLLCNNQTLVCGLCHCEAGRYGELCQCRGATA
CPVGLQGLTCSGSAGICKPDNCYKCQCLGSYFGDACECDRLKCLTSSSGICSGESNGLCSCSGSNVACSCKRA
PLSNITYMGSACNCDPDDCVNRETNATCSDPAIGSTLCHCTGSPCSCSCSCPANTVPPLCEPQALVNSRCVAA
KQCAECGSTKALSECTECVFINSDSQAYKCGVIVSGTCTDSHTYYVDTQKRVYLKRNTVTCDPGPGPIIIVFS
VLGAIIALGLIFLIIAKIILICLDQVEYKKFTSQLEGADWAPRNNPLYMSPEQNYTNVLYRKRSYRGSK*

>EmITGB5_(m.219092)
ASLHEFCLMCLYYGVHTQRVVHRAMITRDQLQWVLLVLACLCDLDSAQQLCSSQTNCSLCIQASPSCQWCSDP
SYAGSRCFSFDTPNINCSKSFVENPIGTKTAQTMDVLGSTVQISPGVVNITVRPGSVTNFTLSIRPARNYPLD
LYILTDLSYSFSNDLSTLKTLGTNIANTLYNITTDYRIGFGSFVDKKVSPYVDVTPSSLTNTNAPYSFKHSVS
LTSNITLFNNRLEAQVISYNQDAPEGGFDGFMQILLCKKLIGWRDNARHLLLYDTDADSHQAGDGKLGGVVKP
NPHTCLMNDTYGLQDVDYQAYGLYDYPSLGDIREQLRINNVIPILAVTSDELARYTAYTTELQSVGATVGTLA
ADSSNILNLISSAYKSVTQKIVFDPVLPAGISSLKFIPINCPQLESDGITCSGVQIEQTVNFTVQVQLASCAN
NQTMQIPVRVVGFGTFTINVQPICNCGCESSGTATNSTSCTNGNGILSCGVCKCNPGRFGTLCQCNSQGVGQG
SSTSCPTGPNQMQCSGQSRGTCVCGKCECATFQDSRFGTTSTYFGSACECDNSRCDTTNGQLCGGTNQGVCQC
GGCQCLGSYYGSACQCSNSLCVDPNDSQGRVCNGRGTCSCNQCSGCQAPFTGVYCQSCQASTPGACASFLCQP
NLVCAQCALGQVNNTLCSSCPSLFLLNTTDINSIQGSITQCQFTDSKGCTYTYYTVEDANVVLQSVYVDSTPV
CPTVLSPASIAAIVITPLVVIAIIGVAILLTVMLIFYLLNRAELRRFEKEVSKASFAKNLNPLYIAASTDIMN
PIFDGGEVKGTAM*

>EmITGB6_(m.33363)
AKQCQELIEYTAYPIGPMATFIVSSVLLLCIASATVWGLRQTSCDTRTTCGDCIASSPQCVWCSDQNITGSRC
FAQGSGQNCSQSGLQNPKGGILNKSQETLSPTNQISPQRISVSVRPGEDIAFGPLSVKPARNYPLDIYLLMDL
SYSMLDNLQNLKMLGAQIASKIVDVTTNYNLGFGSFIDKKLSPYINILPSLLKDPCAPPYGVGGCVPTYSFKH
AISLTSNNTEFNTKIQEQNISANVDPPEGGFDGILQAAVCQQLLKWRHPARHLLVFITDGPYHQAGDGKLGGV
VTPHPGTCLMDQTITDRAVEYEKAVIYDYPSLGQLRDKLEQYDILPIFAVTSEVQKLYSDVASQMQDIGAQVA
TLARDSSNVVDLISETYSKVAQQIQFQADLVPGVSINVVPRNCTKIQNGICSGIQIEQEVQFDVHVSLDGACT
PELQSGPKEVNVRVLGFGSFTVEINAICRCPCEDKPVKNSQLCSSGNGTQVCGLCVCNTGRFGDQCQCDGKSA

STTNSSTCDLGFNNLPCSGNTRGNCVCGKCQCSEFKDTRGNTGRYYGNKCECDDLSCESSNGALCGGVSQGAC
QCGVCKCKPGYSGSACQCSDLFCINPLDPKSQICSGQGKCSCNTCVNCKEPFTGRYCHSCMSTLNQCTNYYCK
PNQPCAMCAVGVAKGPQCDKCTNFTSVETFDSYPIDSLANCQFTDDNDCIYKFYINTTLRVVVATTPSCNVLP
PWFLAAVIAGPLVGLAILGLIILAIVAIIMHILNAVELKRFEKELKSAKSTKNDNPLFIRANTEYVNPIYGK*

>EmITGB7_(m.56163)
TMDKKRLMVISTVILFARSVLLVKGLNCQTAKSCSECVQRGVDCIWCTLPNVTYHCVQRNTAEAHSCGMYVED
KVSNFRSKEKNDLLNKTLISPQSAYIQVRVGDSVAFNVSVMTSRTYPVDFYILMDLSSSLRDDVETLKNTTHK
IVATLLNISSNYAVGFGSFVDKPVPPFVPNIPYSTPTPKGPVCFNHQAQCAEPYSYRHILTLTNDSQKLQYIV
NTQLNISHSSDNPESALDGAGQVLACKKLIGWRNEAFHMMMIITDANYHRAGDGKLGGVIVPFDGRCHMDRIA
AELYQYNQSTFYDYPSIPQLKNLFTDVGVTPIFAVTKTAQNYYKDLVKQLGTGVVGELANDSSNLAQIIAEKY
LKAIGHIQFSVPEIDGLTVTVQAVSGCETKLPSGCADVKLEQQVTFRVTVALDKCTVNIMRQLQSQQSFDLIL
KIPVLAQEFIIHLTPICKCSCEMENEMNSATCHAGGSLACGLCSCNMEQKRFGTFCECQGNQACPIGLANLNC
SGADHGTCLGNCFECQCKEKYFGPACQCNAEYCPAVKGQVCSNHGTCQCPHTVCECNTAPLSQLKYSGTACSC
DPDFCVNPKTNAVCSRSNITEGKTLCHCSENKAQCKCGCTCPAGTALPFCQDETEVSDVCMKQEKCALCVLRM
GGGSKCDGCKAEILSDIATVQLQKSRCPPLLFDGCYYDYFVDPFGTISVSMVPDSCPPASVNPWYIAIGVLGG
VIITGIISLVLIKLILMIMDRVEYKKFAKHLAEANWAQNDNPLYVSPTRHYDNVAYERNRSRPGN*

>EmITGA1_(m.165272)
MIQLLAGAFLISVYGVIAQRLPLDTTVPITRIAPPLRRGDANPDNFGFAVALYQLDPAGTNLSLWRVVVGAPN
GSYPGGLSLTNPSCSTPTINNTGLVYLCSIQPGKDTCDGAVGNGTVDNGRLFDCQAPSNSPNYQQTGASLYSS
GGYLIACAPGFSTGSYSQPVHRGTCYVSFNNSLRTFAQIFPCRSAPVTTFSYYDETYCYAGLSVALKNKIAMF
GNPLAFTTGMASYLQLDTPPVNFSMIAPKNRTVSTGGLVYGPTTMNINGSIVYKTNTLKGYSVGIGRITRSNV
SDYLVATPRWSVDNYVGTVEVFAFPNTGAPLAGPSYPVGNAFSIGVGSNSFFNTDVGVYNSLMNVAAYGTQSG
EQFGASMDTADLDQDGYDELIVGAPFYTDYTKSSYMYEVGRVYVFQNTKGNLSATPIILSSPNPFTGGRFGHV
VLSVGDINGDGTEDFAVGAPYESCTSSDGTSSTGSVYLYTGDRTTFVSQTPIQKITACDIRQRSLTALNNATL
RSFGYSLASKADLDGNAYNDLAVGAFESRAVFVLRTFSVANVTATLTNVGGGVSATKAGCNSSHACGIVMLCA
TYNCSSRPGSCSQSLSMTFDISESTIKAFFNTTTIARTSTVVVTANMLTPNCINVTFYSMLGQVDLSPFKFTA
IVRDLSRDLVSDSGAALADFKAIPVLNGGSASINVGFARACTNPNACASQLALSQTSAVVFKDAQATVEKGLI
VNETASITFSLTVNNSGDEVFGINLVLSAPSFVTGMIITKSSGVPITCTKTNTTGFVCPTVVDGFLAQGNASV
LGVALTIDTTSISFDASNMTFNVSSDDIETNTLDNVVKILLQPTASADLSILSASFSPSGTTYTTPKSITNPT
NLGSIGLLTTFEVTFQRSGPTYIPSLTLTISLPLGGSDWSSYYLYPASVASSVSTIAVSCAAGVLNPYNLQTT
KKRSLSADMIKDLQRGTRQAQNALVLLDCSQSSAGCKNLVCNITNITSTPSFSVRVSLYVNDKYFSARGDNSN
FSVTAAATVTIPTHAIFTGIVSKSFKTAVLNISAAQAPQPKPLNLVVIIVPIVAVVVIIVIAVIVLYACGFFK
RKKREDEDAVEGIDGAVATTAVTKKDPLEDSTVKM*

>EmITGA2_(m.284498)
MMLSATGSRQATTTSIALLLLTAAVWSVQAQNIDTKQPYIRSSPDQTNVDYFGYSIVLHQTIAGNPSSTMLIV
GAPNGTAPGSPVRYTGLIYSCPLNSSITCSGLMGSTTGTDRRLFDTDPNSGSPTQVEEKSGQFLGSVLVSKGD
KFMACGHRYFNWGSNGGYRSSFGRCFIAGRSLRNFAEFQPCDGVGVRQPYSIDVCQAGFSGAIANVSRSGPGA
IAVGAPGTYTWRGIVIRNNPTTNAIQFTAYNPSATILYGYFGYSMTSGYILSKTQEDYLVSSPDLSNLGAVSL
VRNLDTVDVVSEPLQGLQISELYGFSVITADLTGDGYDEVLVGAPLYSPVQNPEAGRVYVYRNIAGTLQFVTQ
LIGDGISYGRFGHAMVNLGDINSDGFADVAISAPFSNDGGKVYIYNGQNINTINTVPAQIIVGRSLLTTANLS
SLIGFGASLASQVDIDGNTYNDLAIGSYQSQQVFVLRTRPIAQMAVSLTASSLLVQVYNGYFPLCTLNSVNYT
CFNVSACVTYTGVGVANQLNLNVTVFGDTTNQLLLLTPRVFFGTNQQASSVVTTVTATKNVQTCSTLNVYIKN
NIADILSSFVFNMSVSVQDFNPAPSNGATSSQDLSLFPILSQSGANTVQVQTNKGGCGACIPVPDLSVEYINT
TYDQKTNSSDNSFIAQETTGISIWLRVTNRKDNAFATVVSFSVPKTQLTFIRCGPDLSFLSSVKDISATVSLC
TCQIANPLLNGEHRDVVIRLEPAPTIDGSQLSYTINFNASSQNAEYSNTTSDNTISLPLAIKTVSALSIDSVG
IVKPEQIIFTSSSVNTSVALTTSLGPYVLTTFTVRNGGPSTIPLVGLDIYWPLDSTTKGLYYLIPTSIKALSS
VFVTQCDSTYTMLLQNDVSNIIPTGNKRSTDGLRRTRAAVASAPLPGGTTTVDCFSQPQSCVRIRCSISQLSQ
DLVTIVVNSTVDSRFFAQGQGLTAQYNFIPRAAVSISGAGSSYIVDMGSNKNASADIVIIYPTQGSSRDLPWW
VYVVIIVPGLLFLLITTVLVLAIIYYCSKRRQAIKNKYQDKQALTGMQGPTDGQ*

>EmITGA3_(m.41332)

MRWMQVWSTVLFFVGWIVCLQSERLDTLAPIIRKSPAASVNSDDFFSYAIALHQIDVPTTGNFKESLDVSRII
VGAPKGTFPGGLNYTHLGEPPVNTTGLVYLCPILNSSCEGLLGNGMSWDRKLFDEDPNVRANELLGIPSTATL
EDKERQFMGASMDSTGDMFVVCAPLWVNTFRHTQSDPDYRPQGRCYYSPRNLTDFHVIQPCNGGTVTSNEGDS
QCTAGIAVTTLNDTFILGAPGQSSGSGALYYTPQLPQYSSPPHGGATILTFTESSFFSIGPTIPLSTYQGFTL
ATGNILDKVKKNVVTSYRQFVGSSYYELLKFYNASDPYVSIMTLPMGESPTENFGYSLVAADLNGDGWDEIIA
GAPMYSTSSMLEIGRIYIFSNYDGTFMSNATVIVTGTISLGRFGHAIVNLGDINGDNCDDLAVSAPYASQNGT
SSSSGVVYIFLGSNANLLNTVPFQTLDAANVMRANQLPELKSFGFSLASGVDVDGNLYNDLVIGALFSQTVVL
YRTLSIALINVTLNAPDDVSIVMQNCSGYACFVVGICASYSGRGLAGPLGLNMEVREVVSGVQPKRLFFGAPV
GKMDAYVSQFQFLSSGGVYCLSVVSYIESSSAGTTSLPFEVQVLFSPNSRNVTSSGPRPLLDLRQYPEINVTG
NNTVQVAIRKNCNTATMCLPDLDLSLNQIVYSSGSGSTPSLVADVTKYINMSVKCASDKDDAFGSTLTATFPS
YLKLTGQNGLPSCATCWICLLLGNSTTSCTFETFLLRANDSIQVELRWSVESATLLGNEKFNISVNVLVPNDI
RTNNNEVTVPIAATAMAELSLELAVDQTSLQYSLAANYSATDTYSLNDFGTTPSKLTMSFLNKGPSTIQNLLL
DIYYPYQSEDTGKLFYLYAGPASNETFNTGIHVSCKGDNPRGIPSVPARTRRSAERYTSLAQWWLDQASGLLP
KRRSVPPSAVVSGTVDCSDLTQRSHYCGHVNCTIKYLLVPADRTSKNLFINLPLYVDDRYFASKKGNYSLVIG
AQVTILNSYIQDTASVSGKTFLTSLTLSPSPVVETTTPNAPIPWYLYVIPAIVGLVFIIIIGCILYFCGFLRR
KRLIPPEADQPQSPRGVPQSAPGTAGQPTPSTPGQPTAQPSQPEGPKTGEASSPEDDLPEKIDDDAYESEI*

>EmITGA4_(m.69874)
MKSTALEGPVLFLLLVVAAAAYNVHQKAPAKAVKAVGGLGELFGFSVALHQFTNGSTVNLVGAPKLFANSSGV
REGGVYVCPTATGSSCYLEPLFSSRLDARNDPGQLLADPAIYPENNFDGQLLGYTLRSFDDHVMACAPLFIGQ
RSGVKARYTGRCVRLPHDFQPSDVPDQITLGIFQDPLEGFLMGLGVALLSEDAVNYNFAVGFGKSSNAPGGVS
YSISVAKSTSTLSVPGGKFYVTQIYDDLLGYHVEAGHVTSPTSYVAIIGAPRGSNHFGNLFVTDLSQANPLTL
FTAKGVQSDGHFGFSFAVCDWMTDGYDSLVVGCPLCDNDVGRVYVYLHSGNPANPYPTVQEVTPPTIVAGRFG
LSVINTGDLDKDGYDDVAIAAPHDAGGVVYIYRGCESGLCDSPQVIRPAAVTQTSLFGYKLSAKVDVDNNSYP
DLSVTDMSGTVYTFRTNPLVVVDTTFEGLASTLNINTDICSVSSFSAACFNFSVCFGYRPLAGGEDIGTFAFS
YTLSVDQSFGRVVQKSTLPTSNQITLSRNTSYCTAQYYYFLKPDSSDTNSPITVQLTLQDVTNPLALSSSPQS
TKFGGSLTPLLDRFVAGSQPRNIVNQSISLITSCGSGGICVPEYVIRTKKEGSEKVLISSEPFFVVIAIENLA
NQSGIAPQLFIDVPSGVGIYGGSSNVSTIAGVQQCVPLTTAQFKCSLSLIQPSSFLNLTIPFILDSAILGVNL
LSGESVLPTLTINFTIGQNNSRAGAMSSLPLILDAQAQYTVTSSTNEAEYVAYSLTSSSLSGQGPALKYTVKV
TNSLVAGTTIPNTTLYIYWPTYFKSVKGQVPLLVVTQDTSPSAPCKHYNTVPQEALRFSNASLPQQHNSTTYN
LPSQYTTDSKSFTYGVIVCDIKNLAPQSSVDVIILSQLWSTSVLTSLMLSIEAAVVSGTSLPNFLGGKPSDSV
TVLLSPGVKEFPVACFPIWIIIVAVIGGFLCLALLGTLIACIFLIYRFVRKSASYDPNADNPDNDDDFVYKQR
TIPPIAGLAESRPTSFQYLSMTGDHAPSQDQETIRKEKEKEMEEELQLQAKLTHTMTRLVKQPQSKEPSEAPS
DDPFETTL*

>EmITGA5_(m.240814)
MTPFLEPLAKTVHVLCIVMACCHAQSIDTKEPIIRTSPDLTSTDYFGYSAVLHQTPVATVVLIGAPNGTAPGS
AVNNTGLVYVCPVTNPGTCAGLTTFTRSSLATDTLLYDRSNNSASEQKSGQFLGGTIISKRGLVVICGHRYFK
PQYTPNGRCFVSNSSLIGFQQYAPCSSSGDRCQTGASASIGNASGVAFPLLGSPGHNGWSGTASRVTVASGAL
RTTTRLSGTAELGYQGYTVASGHILQKTTEDFLVSVPRLNNMGVVNLVMNSATVTVVSQPLQGTQLSEYYGFS
IITADLTGDGYDEVLVGAPYYSPVQNPEAGRVYVYRNNAGSLQFVKQLCGSAENYGRFGHAMTNLGDINGDGL
DDVAISAPFASGGGKVFIYNGVVTTTISSTHSQVIQGNALQSTINLFNLTSFGTSLSSGVDIDKNTYNDLAIG
AYNSGQVFILRTRPTALVAVSLTANPTLVQLTNGLYPSCNLTGTTYACFNLTACLTYTGNGVSNLLNLTVTII
GDTSNQALGLASRLFIGTNSSISTFVTTVGTTKNIQSCLSLPVYIKNEILDKLNGFTMQMNVSVQDFAPPSQN
GNGTLTNLTGYPVLSVQGTSSVQVTNIDRGNCSSGTCIPIPNLAIQFLNISYEVSNGTGQSLVAQQTTNLNLW
FNISNSGQNAFATVLTFLVPKSQLYFIRCDPNLAYLSTIQDYSTALFLCTCQVANPLQGGNYSVIAVRLEPTA
NIDVTQGVIPLMFNVSSQNPENQTTVQDNSVSVQMNITAASGLSVDPIGIARPEQIILSTTTNTTSVNPLGPS
ILTTFTVRNAGPSTIPLVQLNIYWPLNSSETGSYYYLVPTSLQALTTTFFTQCDTTYLNLIASSVNATQAPSS
SGGGNKRRAASLSTSTAQDINKTIDCQLTPSSCVRMQCNISQLIQSQVQITINAALDLRYYTAENIKLTFIPY
VNVSIEGNGANYIVQTSSMKSATANIKVLKRTTASSDSQTGRPAWVGWVIGGICVLLILIIVGLIVIVTVVFI
KKRKAMKDKFGDELKWKTVQVTDNHTDSVRQLIPDN*

>EmITGA6_(m.45476)

HHDPIFVAEKELHCTDEREVAMAKASIYSSAIYLVLTTVCTSVTSPQTIDSKQPIIRTSPDRTNTDYFGYSLA
LHQTPTSTVIIIGAPNGTAPGSAANNTGLVYVCTLTPGTCTGLPAFTRSNLSTDKLLYDTSGNSGEQKAGQFL
GGTIVSKRGLVVVCGHRYFKPQYNPTGRCFVSN

>EmTalin_(m.6774)
MATQTVSLKINITKTNNIKTMQFEESMMVFDACRLIRERVPDAVQGQPTECGLFKPDEDPTKGRWLEMGRTLE
YYHLKSGDMLEYRKKIRPLRVRTLDGSIKTVLVDDSNTVAELTKTVCSRIGLANHEEFSFTVDEETSEMTLRR
QHTLARDQKKLDKLKKELHTDDELNWLNSDKSLREQGISETAVLTLRKRFFFSDQNVDRNDPVQLNLIYVQSK
NAIIDGTHPCTREEAVQFAALQCQIQYGNHNEAKHKPGFLNLDEFLPEEYVKFKGIEKLIWTDHRKLHNLTEL
NAKFRYIQLCRSLRTYGVTFFLVKEKLKGRNKLVPRLLGITRESIMRVDETTKEVLKTWPLTTVRRWAASPNS
FTLDFGDYSESFYSVQTTEGETISQLIAGYIDIIMQKKQPVQNDMADDDDTAVVVDDEVLPNTAVAFRYTGSN
QGTGELDNTANMVMPQQAMTDEGAMFHASGSAQFAEEIGAPGKKHQPHQDSQTAQLLAAQQGLLANIGSAQQA
IGAINKDLLSQAQLPQLGSDSASIKWKQTTLDVSRQNVSSAVAAMLASTASIITLTQGDPMDTNYTAVGSAVT
TISTNLTEMAKAVRLLAALSASQLEGDDLLKAARALAAATAALLNAAQPENMENRQQLLMTSGDMAMSGSQLL
GLVGEQEVDQGTQDALVAMAKAVATATAALVTNAKNVAAKCDDQALQNQVIVAAKQTALATQGLIACTKVLAP
CINSPLCQEQLIEACKLVAAAVEKIVLAAQAACKDGDALRDLGAAATAVTTALNDLIQQIKEGVRMEAGQYDE
ACEAILAATDRLFSSMGNAQEMVKQAKLLAEATSALVNAIKLESENENDPDARRRLLDAARALADATSKMVEA
AKGAARNPGNEQAQEALRKAAEYLRAVTNAAASNALKKKAIRKLEIAAKQTAAVSTQLIAAAQGAGASNRNEA
SQSQLISHCKAVAEQISQLIQSVRASVANPDSPSAQLGLINASMNMIPPAGKMVAAAKAAVPTVGDQAAALQL
GNFAKATASALADLRTATSKASEMCGSLEIDSAIDTVRSLSQEMGEAKMEAQTGQLLPLPGETVESCALELAA
TSKTVGSSMAQLLTAASQGNENYTGMAARDTASALRILGNAVRGVAAGTKNRQTQEYILTTAQQVMDQSCALL
VEAKAAVEDPNAPNKQQRLAQAAKAVSQALNQVVNCLPGQIEFDQAIKAIAQASLTLQAEKFPDASGASYQTL
QSNLSSAAAALNATGSEVVAAARATPEQQAIATVKFAHCYEELLKAGLTLAGASKDKESQNEMLGYLRNISVS
SSKLLLAAKALSADPNAPNAMNQLAAAARTVTDAINSLLNLCSSSGPGQKECDNALRNIEAVAPVLDNPNEPV
SELSYFDCLDMVIEKSKMLGEAGTLITSHAKKGSIEEFGKAVESTASAVCVLTEAAAQAAYLVGISDPSSTAA
IPGLVDQNQFARCNQAIATACQTLLSTSSTQQQVLASATVIAKHTSLLCNACKQASSKTSNPVAKKHFVQAAK
EVANSTANLVKNIKALAADLSEENRQACASTTRPLLEAVEALTTFASSPQFASTPARISEQARVAQLPIVQSG
KNVIKSSSSLLTSAKSLAINPQDPPMWQLLAAHTKAVTDSIKALILAIRDKCPGQKECDSAIDGLNATINQLD
QAILSAMNQQLHPNASSSLQGFQEQLLQAVGDIGEHVKPIATAAKGEAEKLGHQVTAMCNVFPSLAGAAIGAA
SKTTSSQLQISLLEQTKTVTESALQLVYAAKEAGGNTKSTAVHGKVDEAAILVQTAVSELTQTLEKAGSETGI
ITAMVDEIKKAMARVQESPGEVSKTFADYQTDTFTYCKAITKNAQEMVVKASSVSQELPTFSRELTNAYSQLV
DTTQCALATIDSQNIASRLSQNVRALGEACIELVFAGGTLQTSPDDQAARRELTDNAKSVTEKVSYVLATIQA
GAVGTQACNSAIATIMGLVGDLDTTTMFCTAGALHSEDKLGTFAEHRVNILETAKVLVDDTKKLVSSAAGTQE
QLAEAAIQAVKTITAEAEHVKLGAASLATEDMEAQLLLLQAAKDVANALSDLIGATRSAAGKSVQDAAMEQLK
SSAKVMVAKVSNLLKTVKNVEDEAAKGVRSLENAIEAIASDLQEFESSNPPKSQATAEDLIRSTKGITLASAK
AVSAGNSCRQLDISACANLARKAVTELLETCKSAAYKAENGELKAKTLMTGRECATSFKALLELVHQIVLKPT
YEKKQSLPTFSKEVATWVGDVVQVAEQLKGSDWVDLRDPNVIAENELLQAAASIEAAAKKLSELQPRREVRAD
ESLTFEEQILEAAKNIASATSALVKSASAAQRELVAQGKLSSKPQSEDSQWSEGLVSAAKLVAAATSNLCEAA
NMMVQGHAQEDKLIAAAKSVAASTAQLLIACQVKADARSENNRRLQMAGQAVKKATETLVAAAQQAAVEGGRA
DGASGGAASIQVNLGGKVMNRFRQELEIAEQIAAKERELEQARLQLTKIRKGQN*

>EmTalin2_N-terminus_(m.34486)
DLACPSHSSIKMATAKVAIHIHLVDTNDVRSVMFDETMLVCYACSFVREKFLPSNSKDGSASEYGLFKPDEGS
PQWLNVGSMIKDYGLRDGDTLQYKRKVLPYSVTLSDGRTLKFKLDNSRTVAENVKTVCSEAGLPNDYEYSFET
TNLAVKSSKKQDTKAKKRSSSSAKNVWLKANKTFAAQGVDENCILTLKKRFRIVDAPISLGDSTALNAVYAQC
KDDITSGLHPCTEDEAIQLAALQCYVRFGKNRPVSIKIAEFLPLDYVARKDVEQSVLLAHSKLNEMTEAECKF
FYIQLCRSLHTYGATYFLVNEKRKAKKKLAPTLLGFKASEIIRVDKVTKQVVTSWPIENVRRWMGTEDFFKID
FGTASQLIYTVQTTEGLEISEHLSELTDIIEKIKKNNAGNKKDGDREMCKKNDTESQLQTADTICSAEYDLDS
GPFDDAQFSNGCLSAEYPDCPSPRMDCPSPRIMDCPSPRITDCPSPMPPEVGNLLPVTAPGELQNSSSTPQCI
SPSPEEQRMMLPPLQITTEPTLGPTAHSDLPNTALAGYLTLPLPHKKSTAKAGNESDPAEEIQRLQMLILKLQ
MELQMAMDQVNYAEQRAELAEKKAALAEERAKRAEHSFRLSFQMTNQ

>EmPaxillin_(m.5894)

MELDDLLEDLQQALPPEAQAYSSSNTAKWERGSAEMSVSTPSHPQQFDEAQYSEVRRGPIYTPKSPPNSVSPP
PTKPPRTAPAASEGLSELDSLLAMLGDTQAANQEKPPHDTGLSRPTISAFVDELTQLENQVSNKTSASKFTTG
GIPMAAPVTASSATKELDDLMANLSKFEPSTVSVQDDKAPKGAQPKTRSALESLNSMLGSLEEDMSKRHGVST
MAKGTCAACNKIILGKVVNALNMQWHPEHFTCASCDAELGQVTYYESNGRPYCEKDYNELFAPRCAYCNGPIL
EKVMRALDRTWHPEHFFCTLCGKHFGTDGFHEKDGKAFCRECYYEKFAPRCKRCEKAIMEGFITALNAQWHPD
CFTCKVCNVSFPRGNYFDHEGEPHCEIHYHAARGTLCASCQKPVVGKCVSAMGKKFHPEHFTCAFCLKLLNKG
TFKEHRSNPYCQACYIKLFG*
